# Unraveling Microglial Spatial Organization in the Developing Human Brain with DeepCellMap, a Deep Learning Approach Coupled to Spatial Statistics

**DOI:** 10.1101/2024.11.28.625932

**Authors:** Theo Perochon, Zeljka Krsnik, Marco Massimo, Yana Ruchiy, Alejandro Lastra Romero, Elyas Mohammadi, Xiaofei Li, Katherine R Long, Laura Parkkinen, Klas Blomgren, Thibault Lagache, David A Menassa, David Holcman

## Abstract

Mapping cellular organization in the developing brain presents significant challenges due to the multidimensional nature of the data, characterized by complex spatial patterns that are difficult to interpret without high-throughput tools. We developed DeepCellMap, a deep-learning-assisted tool that integrates multi-scale image processing with advanced spatial and clustering statistics. This pipeline was designed to map microglial organization during normal and pathological brain development but can be adapted to any cell type. Using DeepCellMap, we capture the morphological diversity of microglia, identify strong coupling between proliferative and phagocytic phenotypes, and show that distinct spatial clusters rarely overlap as human brain development progresses. Additionally, we uncover a novel association between microglia and blood vessels in fetal brains exposed to maternal SARS-CoV-2. These findings offer insights into whether various microglial phenotypes form networks in the developing brain to occupy space, and in conditions involving haemorrhages, whether microglia respond to, or influence changes in blood vessel integrity. DeepCellMap is available as open-source software and is a powerful tool for extracting spatial statistics and analyzing cellular organization in large tissue sections, accommodating various imaging modalities. This platform could open new avenues for studying brain development and related pathologies.

## 1 Introduction

Recent advancements in digital pathology and multi color fluorescence microscopy, including traditional techniques such as haematoxylin and eosin (H&E) staining [1] and immunohistochemistry [2], as well as more sophisticated multiplex imaging approaches [3, 4], have transformed our ability to map cellular organization directly within tissues. Coupled with automated image analysis, these methods have provided invaluable insights into key biological processes, spanning fields such as cancer biology, immunology, and developmental neuroscience. While fluorescence and brightfield microscopy have been widely employed, their potential is enhanced when integrated with advanced spatiotemporal cell mapping techniques, which enable the comprehensive analysis of cellular organization, dynamics, and morphology in complex tissue environments.

Microglial cells, originating from extraembryonic yolk sac progenitors, migrate into the developing brain from the 4th postconceptional week (pcw) and colonize the telencephalon through migration and cycles of proliferation and apoptosis [5]. During this process, microglia display marked morphological heterogeneity, reflective of their diverse roles in brain development and neurodevelopmental topography establishment [5, 6, 7, 8, 9, 10]. Understanding how various microglial morphologies are organized within the developing brain and the spatial relationships between their distinct morphologies can provide critical insights into their functional roles. Despite their importance, there is a lack of high-throughput, automated tools capable of mapping these interactions, limiting our ability to fully characterize microglial organization during brain development, particularly in humans. In this study, we have developed an integrated computational pipeline combining advanced image processing with deep learning to systematically identify spatial relationships between microglial cells in tissue. Our method is designed to recognize cell morphologies embedded within the tissue environment, and reveal their intricate relationships using advanced spatial and clustering statistics. This framework enables us to map and quantify the organisation of microglia and other macrophages, providing a high-resolution view of their spatial arrangement and heterogeneity.

Microglial organization is known to be disrupted in various pathological conditions [11, 12]. For instance, hypoxic conditions drive microglial proliferation, leading to altered spatial distributions and cellular organization [13]. Additionally, maternal infection with SARS-CoV-2 during pregnancy has been linked to fetal cortical hemorrhages, where compromised blood vessel integrity may disrupt microglial organization [14]. However, it remains unclear to what extent microglial spatial dynamics are affected under these pathological conditions. Understanding how microglia respond to such perturbations may help elucidate their role in injury and repair mechanisms, particularly in relation to blood vessel integrity.

Historically, the study of microglial morphology has been largely descriptive, relying on manual quantification methods that are time-intensive and prone to bias [5, 7]. Automating the recognition of microglial morphology and extracting spatial statistics to analyze their interrelationships allows uncovering novel organizational patterns that are otherwise missed by manual approaches. Recent developments in automated image analysis, particularly with the advent of deep learning algorithms, have significantly improved the segmentation of cells and tissue regions [15, 16]. Both supervised and unsupervised machine learning models have emerged as effective tools for whole-slide detection of microglia, particularly in brightfield microscopy, where segmentation accuracy is critical. While these advances have shown great promise in mouse brain sections, their application to human post-mortem tissues is still in the early stages [17, 18].

Accurate cell segmentation is a critical first step in the meaningful analysis of tissue organization. In fluorescence microscopy, where cell types are often color-coded, segmentation alone can reveal cellular distributions. However, in brightfield microscopy, an additional classification step is required, which must consider both morphological features and spatial information related to the cell’s local neighborhood. This is particularly important in developing tissues, where various cell phenotypes, such as microglial cells, frequently intermingle. Current methods, primarily designed for postnatal non-human tissues, face scalability challenges due to small sample sizes and reliance on manual segmentation. These approaches often struggle with the complexity and diversity of cell morphologies encountered during development, where there is significant interplay between different cell types [9, 19].

To gain deeper insights into the spatial relationships between distinct morphologies [20, 21], advanced spatial statistics can be leveraged to uncover associations between different cell phenotypes and to analyze their spatial distributions within tissue. Traditionally, spatial analyses in digital pathology have been limited to methods that quantify the accumulation of specific cell types in regions (e.g. tumor islets) [22], or that evaluate cellular neighborhoods based on local cell type stoichiometry [23]. However, deeper spatial statistics that examine cell type proximity (called cell-to-cell association or coupling, see Supplementary Material for specific definitions) [24], remain under utilized [25]. These frameworks are often difficult to generalize to complex cell shapes, like microglia, and to quantify the accumulation of cells at anatomical region boundary (that we call cell-region association, see Supplementary Material). Moreover, these statistics require careful tissue-specific considerations (e.g., cell density, regional boundaries) and statistical processing to ensure accurate biological interpretation [26].

A promising approach to characterize cell-to-cell and cell-to-region spatial relationships involves a recent methodology called levelset analysis that partitions the tissue into sub-regions with respect to the distance to a given cell type or a region. Then, the statistical analysis of the number of a second population of cells (e.g. another morphological type of microglia) in each sub-region enables the robust characterization of how certain cell types accumulate around others or in proximity to specific tissue regions [26]. However, such analysis must also account for uncertainties arising from the deep learning-based classification of cells.

To address these challenges, we introduce here DeepCellMap, a novel computational approach that we summarized into an open-source Python package developed for robust morphological cell classification using deep learning, coupled with advanced statistical characterization of spatial relationships between different cell morphologies/classes and their distribution within tissues. This pipeline was developed on microglial cells during development. DeepCellMap integrates a generalized Ripley’s method that accounts for classification uncertainties in the deep neural networks, enabling a detailed analysis of cell distributions in tissues and with respect to the boundary of a brain region. To further investigate cell clusters, their organization, and the spatial dynamics of mixed microglial phenotypes, this method also features an optimized Density-Based Spatial Clustering of Applications with Noise (DBSCAN) algorithm [27], with semi-automated parameter calibration. Compatible with both brightfield and fluorescent microscopy images, DeepCellMap stands out as a versatile tool for analyzing complex tissue structures.

In this study, we first describe DeepCellMap and then apply it to histological brain images that we labelled with immunohistochemistry and immunofluorescence from normal human fetuses and fetuses whose mothers were exposed to SARS-CoV-2 during pregnancy [28, 29]. By fusing machine learning with advanced spatiotemporal statistics, this approach enables the characterization of microglial organization over time in both healthy and diseased conditions. The flexibility of this method also allows for its adaptation to diverse datasets of dynamically changing tissues, offering a powerful tool for broad applications in biomedical and clinical research.

## 2 Results

### 2.1 Cell morphology classification and advanced spatial statistics

#### Cell detection

To map the distribution and spatial relationships between various cell classes or morphologies in tissue samples, derived from different imaging modalities such as brightfield immunohistochemistry or high-resolution confocal microscopy, we developed DeepCellMap, a fully automated analysis pipeline processes labelled images as input (Figs. 1A and S1). Given the typically large size of these images (many Gigabytes), a pre-segmentation step divides them into smaller, manageable tiles (Figs. S2 and S3). In each of these tiles, cell shapes are automatically segmented (Fig. 1B1) using image thresholding and morphological operations, ensuring the precise identification of cellular boundaries and features (Figs S4 for method and Fig S5 for validation).

**Figure 1.**
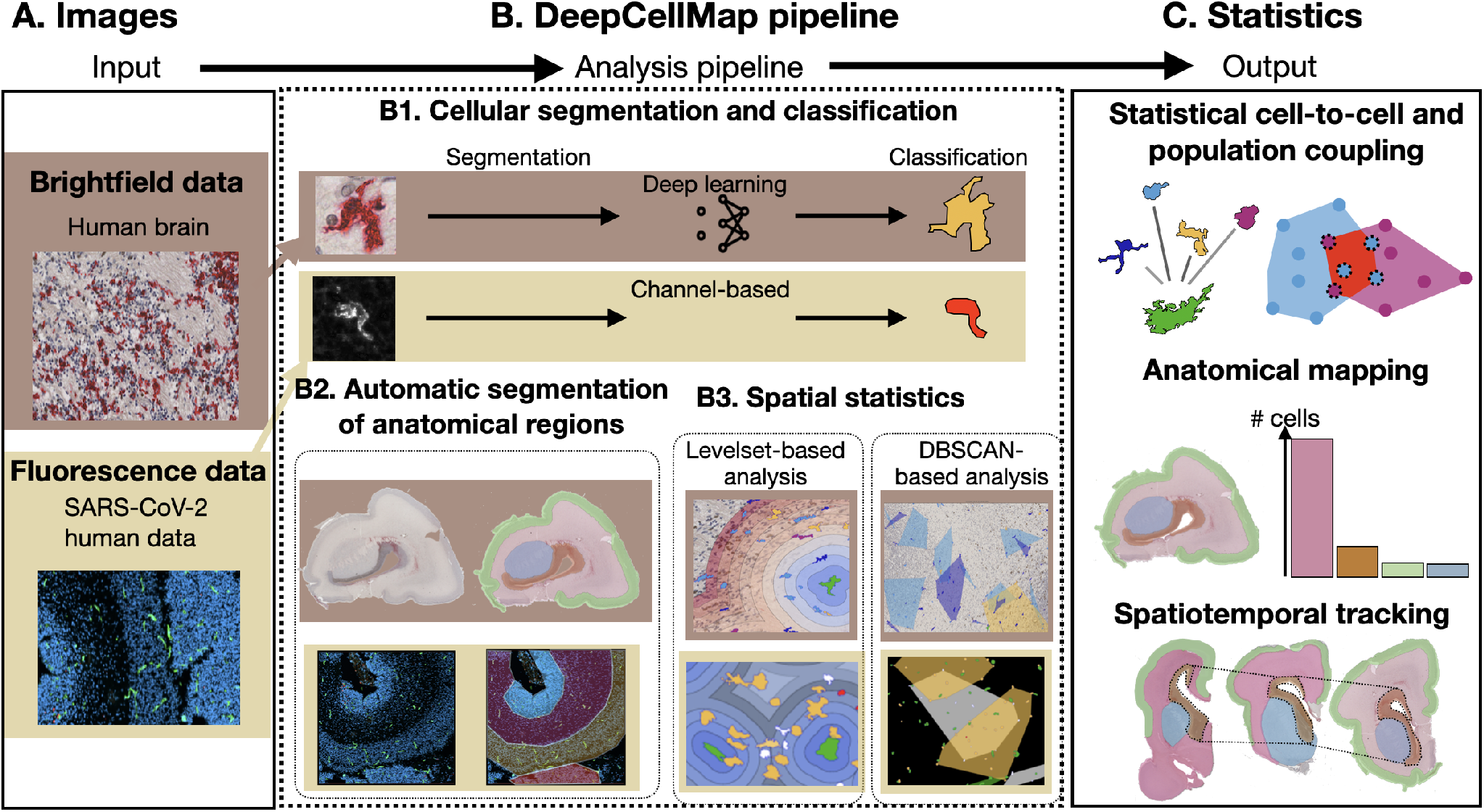
DeepCellMap principle. DeepCellMap is an open-source Python platform that segments and classifies cells, and extracts spatial patterns within tissues. **(A)** Input can either be brightfield (e.g. immunohistochemistry data from human developing brain) or multicolor fluorescent images (e.g. SARS-CoV-2 labelling). **(B)** first series of functions is dedicated to cell segmentation. The classification of cell morphologies in brightfield images uses a trained deep-learning network (U-Net), while the classification in fluorescent images is based on colour channel separation (**(B1)**). To track the temporal changes of cellular distributions in different regions of the tissue during development, DeepCellMap embeds algorithms to automatically delineate specific tissue regions based on the global cell density **(B2)**. From extracted cell types/morphologies and positions within the tissue, DeepCellMap computes spatial statistics to probe cell-to-cell and cell-to-region associations with levelset-based analysis, and the overlap between the different cell clusters with DBSCAN-based analysis **(B3). (C)**. DeepCellMap provides detailed anatomical mapping of different cell types/shapes and enables the measurement of the accumulation of cells in different anatomical regions of the tissue. The spatial statistics allow the in-depth characterization of the association between different cell types (cell-to-cell and cluster overlap) and the evolution of these metrics across time.

#### Cell classification

Microglial cells display an array of morphologies in homeostatic, pathological, and developmental conditions in humans and rodents. Microglia have been described as amoeboid, migratory, rod-shaped, phagocytic, ramified, intermediate/hyper-ramified, multinucleated, activated, reactive, dark, dystrophic, aggregated, bulbous, and more [30, 31, 32, 33, 7, 5, 34, 35]. Some of these morphologies evolve from one to the next across developing tissues and are tied to function as shown in Fig. S6A. Some morphologies coupled with functional markers inform us about what these cells are doing such as proliferative, dying and phagocytic microglia. To detect these cells, position, local neighborhood and shape are key parameters.

Following cell segmentation, DeepCellMap classifies microglia into distinct morphological categories that likely reflect their types. In fluorescence imaging, cell types correspond to specific labels and color channels. For example, in our SARS-CoV-2 dataset, different cell types are associated with specific (combinations of) color channels. However, in brightfield imaging, identifying different cell types from the segmented masks is less straightforward.

Although there have been some attempts to automatically determine cell types using machine learning in well-defined imaging conditions [18], this process is still largely performed manually, particularly in developmental stages where diverse morphological types coexist within tissues [9, 19]. DeepCellMap integrates a supervised deep learning classifier based on a U-Net architecture (see Methods) to classify microglia into 5 distinct types (see Figs. S7, S8, S9 and S10). The classification relies primarily on morphological features and information about the local cellular neighborhood in the tissue [36].

The classifier achieved an F1-score of 81% on the training dataset.More specifically, the proportion of well-classified cells in each class were proliferative : 85%, amoeboid : 85%, aggregated :83%, phagocytic, 75% and ramified 75%.

#### Delineation of brain regions

Automatic classification enables the collection of a large number of cells across different brain regions. The third step in the DeepCellMap pipeline involves the automated delineation of brain regions (Fig. 1B2). Distinct differences between cell densities exist between regions and we developed an automated method based on the local density of detected cell nuclei (see Methods). Cell nuclei segmentation was performed using the well-established deep learning algorithm, CellPose [16], which facilitated accurate mapping of cell density across the imaged tissue. By applying automatic thresholding and morphological operations, we were able to successfully reconstruct four primary anatomical regions within the brain tissue: the striatum, neocortex, the ganglionic eminence, and the cortical boundary.

#### Mapping cell-to-cell spatial association

To map the spatial relationships between the cell morphologies/classes identified by the deep learning classifier (Fig. 1B3), as well as between cells and specific tissue regions, we developed a statistical framework that accounts for the local cell density and corrects for potential misclassifications from the previous deep-learning algorithm. Many existing methods for analyzing cell distribution in tissues focus on defining homogeneous cellular neighborhoods and extracting statistics, such as the accumulation of specific cell types within given regions [23, 37, 38]. However, few approaches offer a statistical characterization of spatial relationships across varying distances between cells [39]. Here, we build on the Statistical Object Distance Analysis (SODA) framework [40, 41], which corrects for non-significant spatial association of cell populations that may arise from random distributions, providing key metrics such as the percentage and mean distance between spatially associated cells. To examine the spatial distribution of one cell morphology *B* relative to another *A*, SODA employs a levelset method to map the spatial neighborhood of *A* cells, where the 0-level contour defines the boundary of *A* cells. The surrounding area is partitioned into subregions (*ω*_*i*_, i = 1..N), and SODA adjusts for cell density variations to control for random spatial proximity (null hypothesis). This enables robust quantification of significant spatial associations between different cell types, or accumulation around a given tissue region, offering a deeper understanding of cellular organization and interactions.

Uncertainty in cell type classification can introduce errors that propagate into the spatial association analysis. For instance, in a scenario where type *B* is frequently misclassified as type *C*, any observed association between types *A* and *C* would, in fact, reflects the true association between *A* and *B*. To mitigate these effects, DeepCellMap incorporates classification uncertainty by weighing the cell counts in each level set region based on the output probabilities from the deep learning classifier (see Methods). Furthermore, we utilized the confusion matrix from the training dataset as an *a priori* correction, adjusting for potential misclassifications between cell types during the cell-to-cell spatial association analysis.

To validate the ability of DeepCellMap to accurately measure the level and distance of association between different cell types, while accounting for potential classification errors, we designed synthetic simulations (Fig. S11). In these simulations, two cell types *B* and *C* were spatially associated with a third cell type *A* across a broad range of association parameters (level and distance). To model potential misclassification between cells, we created three scenarios: no confusion, intermediate confusion, and high confusion, where the classification error between *B* and *C* was set to 0%, 30%, and 45%, respectively (see Methods). The reconstruction errors, defined as the absolute differences between the association level and distance estimated by DeepCellMap (with and without correcting for misclassification) and the simulated parameters are presented in Fig. S12 for the three confusion scenarios.

Overall, applying DeepCellMap without correction of classification errors leads to incorrect estimates of the association score (up to 40% error in case of intermediate confusion according to Fig. S13A and 45% in case of high confusion (Fig. S14C) and can lead to errors of up to 400 pixels for the evaluation of the association distance in both scenarios (Fig. S13B,D and FigS14B,D). We observed that the accuracy of DeepCellMap decreases as the standard deviation of the association distance increases, thus as associated cells are distributed across a larger number of level set regions, performance declines slightly. Nevertheless, while the algorithm tends to slightly underestimate the level of association, it maintains high accuracy in estimating distances, with an average error of less than 5% across the entire range of parameters tested. These results demonstrate that DeepCellMap reliably characterizes spatial associations between different cell populations, even in the presence of classification uncertainties and errors.

#### Cell cluster overlap

Level-set analysis is highly effective for probing cell-to-cell and cell-to-region spatial associations, but it has limitations in describing the relationships between the spatial territories occupied by different cell types within tissues. To address this issue, DeepCellMap incorporates a semi-automatic DBSCAN-based algorithm, where one of the key parameters is automatically optimized (see Methods). To validate this function, we performed synthetic simulations where we varied the percentage of points organized in clusters and measured the performance of the optimized DBSCAN algorithm to retrieve the simulated proportion of clustered points (Fig. S15). DeepCellMap was able to recover a large proportion of clustered points, with a slight underestimation probably due to the difficulty of taking into account points at the edge of Gaussian clusters.

Therefore, the present clustering analysis enables the robust detection of cell clusters, and computes the proportion of cells distributed in clusters *versus* isolated. The method also quantifies the degree of overlap over time and between different cell type clusters. Combined with brain region delineation based on relative cell densities (as described above), this clustering analysis allows for spatiotemporal tracking of the colonization of various cell types during development (Fig 1C).

### 2.2 Microglial colonization in the developing human brain

The colonization of the developing brain by different morphological types of microglia cells remains difficult to evaluate and could benefit from an automated identification of their spatial organization within different regions of the brain [5, 7]. Using immunohistochemical labelling of microglial cells and brightfield imaging of post-mortem brain tissue at different developmental stages (Methods), we used DeepCellMap to characterise distributions and spatial associations of different microglial phenotypes (morphological and functional) in different brain regions.

#### Microglial quantification in distinct anatomical regions

Using DeepCellMap, we automatically delineated and segmented the striatum, neocortex, cortical boundary and the ganglionic eminence (Fig. 2A1-A2) and extracted microglial counts by morphological class in each region using a segmentation pipeline based on Cellpose and Otsu methods (Fig. S16 and Fig. S17). We found (Fig. 2B) over three time points 17, 19 and 20 pcw that the total number of microglia was not continuously increasing over the covered time period. For example in the neocortex (pink labelled), the number first decreases from 10928 to 6926 and then increases to 14262. The number of microglia in the other regions was increasing. We then quantified the proportion of the 5 classes of microglia in the same regions (Fig. 2C) using the deep learning classification approach integrated to DeepCellMap. We observed from 17 to 20 pcw, an increase number of amoeboid microglia in the ganglionic eminence and a net increased proportion of phagocytic cells (see also Fig. S18). The large proportion of amoeboid cells in the ganglionic eminence aligns with their role in migrating to colonise nearby structures such as the striatum. We also found that ramified cells are dominant in the striatum, neocortex and cortical boundary, most likely because these regions are maturing at a faster rate than the ganglionic eminence, which remains mostly populated by amoeboid cells (Fig. S19). Overall, DeepCellMap allows the classification of microglial classes according to morphology, the automated segmentation of anatomical areas according to cell density and the further characterization of the total and class-specific microglial cell counts in the different brain regions during development.

**Figure 2.**
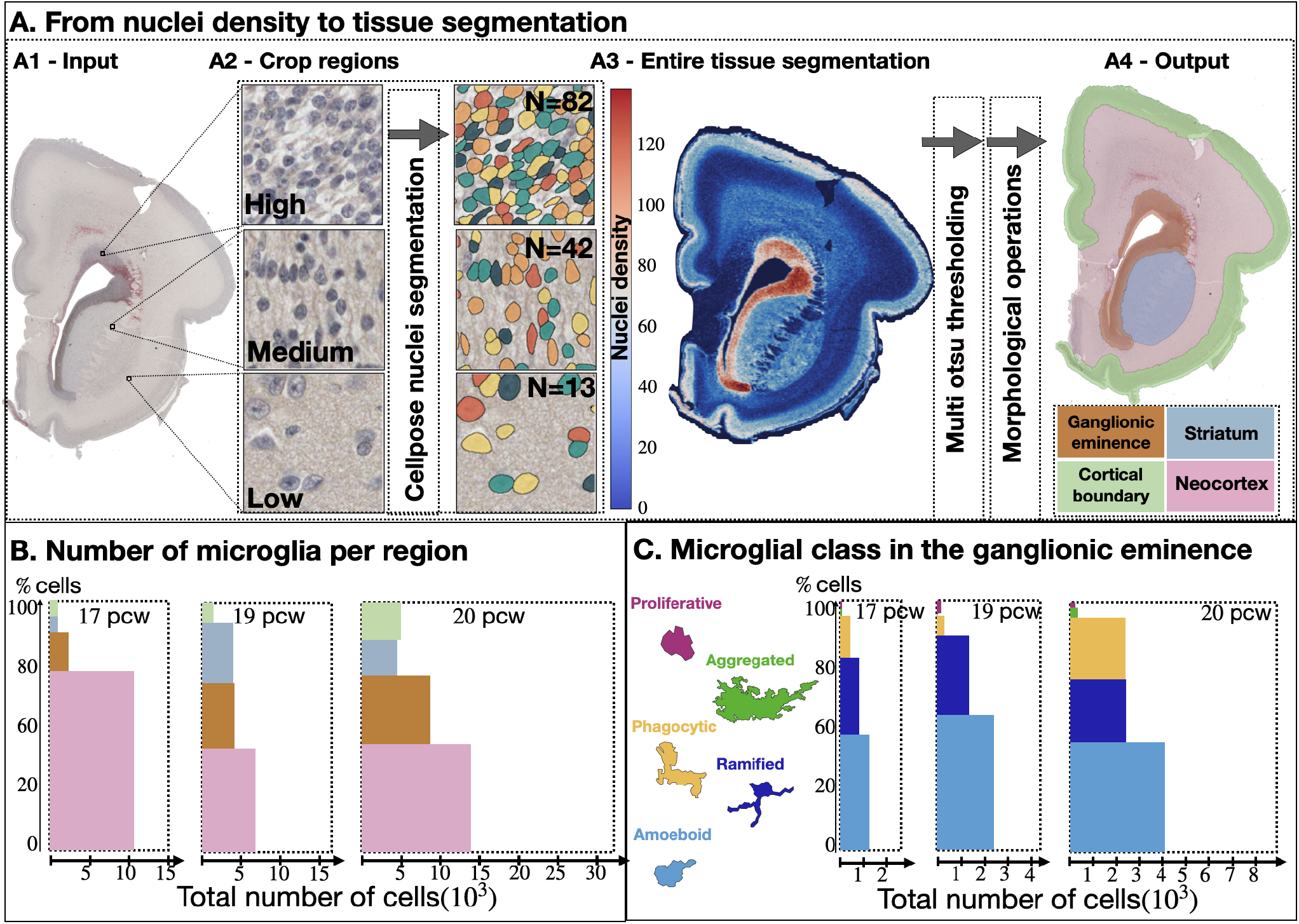
Tracking microglial colonization in different brain regions. **(A)** Detecting cell nuclei performed by CellPose segmentation algorithm [16]. **(A1)**-Image of a fetal tissue section at 17 pcw. **(A2)**-Cropped regions (256×256) showing three different densities (high, medium and low), where we identified 82, 42, and 13 cells respectively. **(A3)** Nuclei density over the entire tissue (left color bar). This step is followed by the Otsu-thresholding and morphological operation pipeline (Figs. S16 and S17). **(A4)** Automatic delineation of four brain regions based on nuclei density: striatum (blue), neocortex (pink), ganglionic eminence (orange), cortical boundary (green). **(B)** Percentage and number of microglia estimated from the deep learning algorithm for three pcws (17, 19 and 20) (n=3). **(C)** Distribution of microglial morphologies in the ganglionic eminence.

#### Spatial associations between microglial phenotypes

To study the spatial associations between microglial morphologies, we first applied the level set algorithm in DeepCellMap after choosing a region of interest and defining the states to be classified (Fig. 3A-C). Briefly, this method allows us to map the spatial neighborhood of a given cell class (morphological type) and extract the number of cells from another given class in level set-defined regions. Importantly, the DeepCellMap implementation of levelset analysis corrects for the uncertainties in classification and the expected accumulation of cells in different level sets for a random distribution (Fig. 3D).

**Figure 3.**
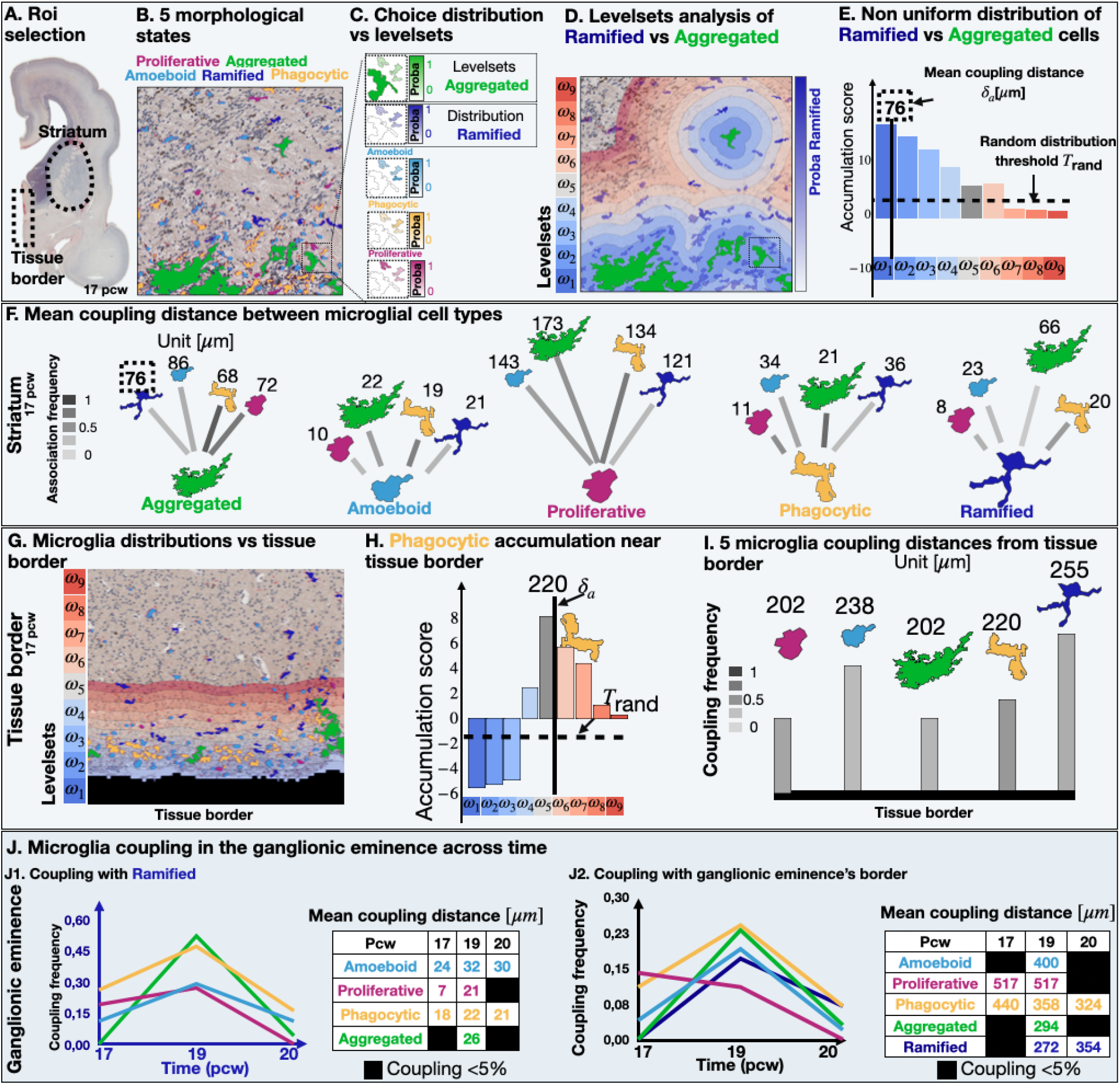
Microglial distributions according to their neighbors (based on level set segmentation. **(A)** Choice of 2 regions of interest: Striatum and tissue border in the fetal brain at 17 pcw. **(B)** Five microglial morphological types are identified in developing tissue: proliferative, aggregated, phagocytic, amoeboid, and ramified, 2 of which are chosen. **(C)** Choice of the “aggregated” microglia as the center of levelset construction vs “ramified”. **(D)** Ramified localization color-coded by their coding probability (right blue scale) in the level set representation of the aggregated cells (green), obtained from a gradient distance. The distances are divided into 9 subregions (*ω*_1_, …, *ω*_9_), color-coded (left scale).**(E)** Statistics of ramified cell accumulation in the levelsets. The Accumulation-score is defined as the ratio 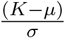, where *K* is the generalized Ripley function based on U-Net output probabilities (Methods) and the expected mean *µ* and the standard deviation *σ* of ramified cells in each region *ω*_*i*_. The statistical threshold 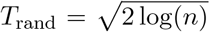 (horizontal dashed line), where *n* is the number of level sets, is used to identify the *ω*_*i*_ regions where the microglial distribution deviates from the uniform one. The mean coupling distance δ_*a*_ = 76*µm* between ramified and aggregated measures the mean distance between the two populations, with weights accounting for the deviation in each region *ω*_*i*_ of microglial distributions (see method). **(F)** Coupling result shown as trees, where the root (defining level sets) is a chosen microglia type, and the leaves (microglia distributed in the level set) are the 4 others, quantified by the mean coupling distance in *µm* (above) in the striatum. The distance lines are color-coded by the frequency of association (left scale bar). **(G)** Microglial cells in the level sets generated by the edge of the tissue or a region. **(H)** Distribution of phagocytic cells in the level sets *ω*_*i*_, with the same threshold as above. **(I)** Coupling distances and association frequencies for each morphological type with respect to the edge of the tissue. **(J)** Spatiotemporal coupling of four microglia types vs ramified (level set segmentation) in the ganglionic eminence (interactions where coupling frequency is below 5% are discarded) **(J1)**, and coupling of the 5 microglia types with respect to the edge **(J2)** across time.

First, we report here that all microglial morphologies are spatially associated with one another but at different levels. In particular, three microglial morphologies are most frequently associated with each other (phagocytic, aggregated and proliferative) whilst ramified and amoeboid are less coupled with other cell morphologies. The level set analysis also allowed an estimation of the mean distance between associated single cells. For example, we found that the mean distances weighted by the level set (see Method) is 76 *µm* for ramified versus aggregated (Fig. 3E) in the striatum (Figs S20, S21 and S22). Interestingly the mean distance from all morphologies to the aggregated class was ∼75.5*µm*. The phagocytic class was the closest on average to the aggregated class with a mean distance of 68 *µm*. When we performed this analysis on all groups, we found that amoeboid, phagocytic, and ramified had mean distances to the other groups of 18, 26, and 29 *µm* respectively. This is in contrast with proliferative that were on average at the largest distance of 143 *µm* from all other classes (Fig. 3F).

Second, we estimated the mean distances between the 5 microglial classes and the border of the brain region (Figs 3G-I and S23): we found that proliferative and aggregated microglia were the closest to the border with a mean distance of 202*µm*, the others being at a distance of the border of ∼238*µm* on average. At this stage, we concluded that the levelset analysis reveals a certain variability of coupling, that departs from a uniform distribution, but this variability depends on the morphological class. Moreover, in the ganglionic eminence, the average coupling distance between amoeboid and phagocytic versus ramified (Fig. 3J1), was quite stable across time, with a mean distance of 29*±* 4 *µ* and 61 *±*2 *µm* respectively. For other cell classes (proliferative and aggregated), the coupling frequency was less than 5% for 1 and 2 time points respectively, but when the frequency was higher, the coupling distances with ramified cells were of the same order of magnitude between the ganglionic eminence and the striatum for 17 pcw. However, the distance between the boundary of the ganglionic eminence and each population was changing over time, but the coupling frequency remained quite low, showing that this effect could be marginal (see frequency of coupling tables) (Fig. 3J2).

To conclude, our key findings here suggest that three microglial morphological classes are most frequently coupled with one another (phagocytic, aggregated and proliferative) whereas ramified and amoeboid are less coupled in general. Whilst the coupling distance remained stable for one time point between the striatum and the ganglionic eminence, it doubled when we considered the ganglionic eminence border versus the cortical tissue border. Importantly, on average, the cell-to-cell coupling distances were much smaller (*<*100 *µm*) than the average cell to region border coupling distances (200 *µm* on average and could reach 500 *µm*). In the context of our knowledge of the rodent literature, microglial proliferation may be associated with increased apoptosis which could explain the strong coupling between our phagocytic and proliferative classes. Microglia are phagocytes and it is precisely during these temporal windows that we expect an increase in cell death as the brain prunes extranumerary cells. Dying cells influence the distribution of microglia and this is the first time that the coupling association between proliferative and phagocytic and aggregated microglia is reported in humans. We also observe an overall increase in phagocytosis genes as development progresses (Fig. S6B-D) which mirrors our histology data. Aggregated and phagocytic microglia accumulate very closely to the border and a drastic decrease of association at 20 pcw is visible, possibly linked to distinct territories being colonized by the different types. Finally a decreased accumulation from the ganglionic eminence border may suggest that cells could be migrating from the ganglionic eminence towards other regions of the brain.

#### Identifying distinct territories occupied by microglia during brain colonisation

To further assess whether microglial morphological types could be non-uniformly distributed, we developed and applied our semi-automatic DBSCAN clustering algorithm in DeepCellMap (Methods). We obtained an ensemble of clusters for each morphological state delimited by their convex hull, as illustrated with the ganglionic eminence (Fig. 4A-B).

**Figure 4.**
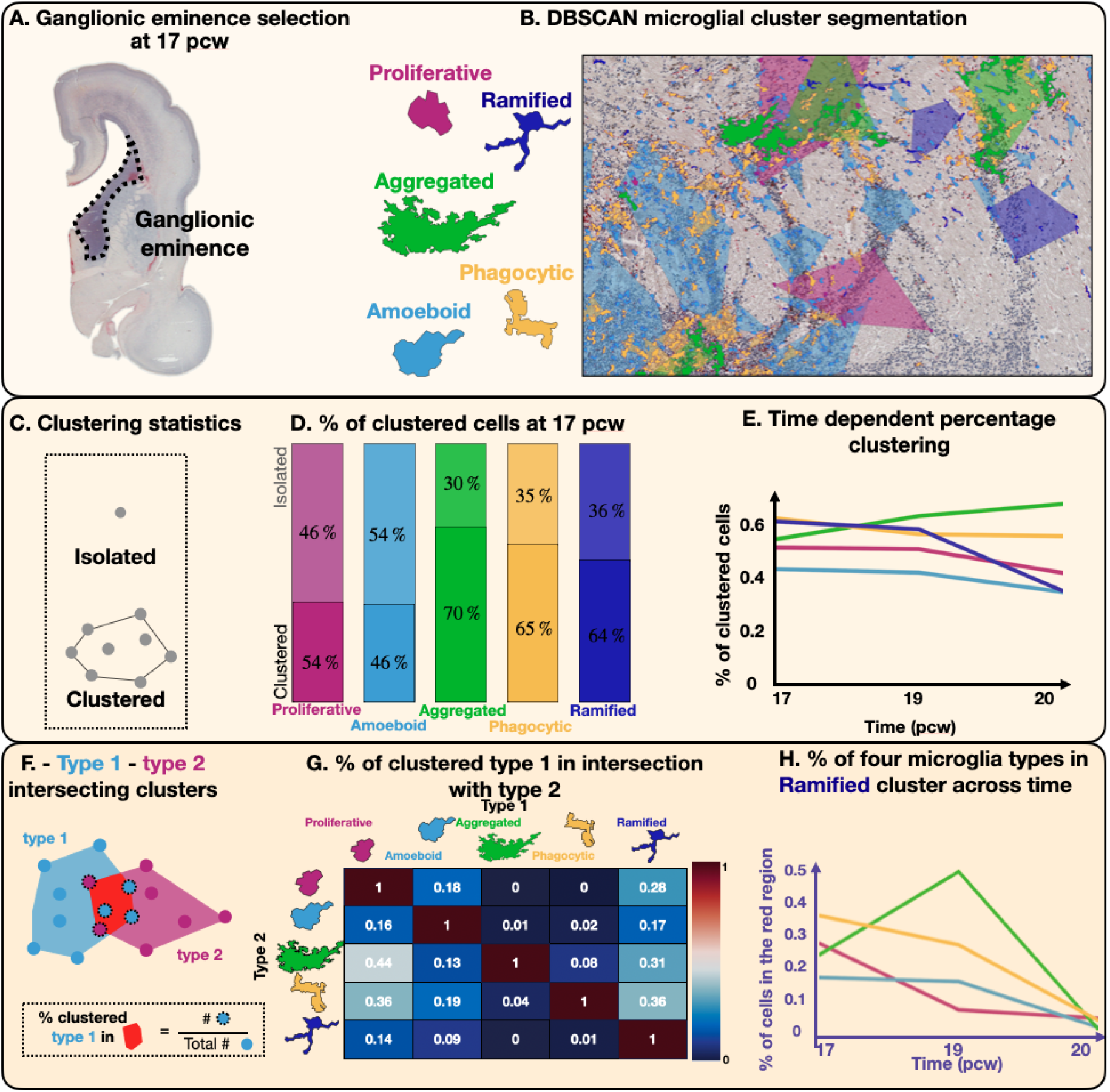
Overlapping of cell clusters in the ganglionic eminence. **(A)** DBSCAN application to the (x,y) coordinates of the cells’ center of mass. **(B)** Graph of *Min*_*Sample*_ versus *∈* used to select the value of the radius *∈* that maximizes the number of clusters. **(C)** Schematic representation of the procedure to select stable clusters by removing 10% of the cell located on the edges of the convex hull while the area of the new convex hull remains *>* 60% of the initial surface. **(D)** Clustering of the five morphological states using DBSCAN algorithms, generating convex hulls around each clusters. Color code is the same as microglial morphological types: proliferative (pink), amoeboid (light blue), aggregated (green), phagocytic (yellow), ramified (dark blue). **(E)** Scheme of isolated vs clustered cells. **(F)** Percentage of isolated vs clustered cells for the five populations at 17 pcw in the ganglionic eminence. **(G)** Time dependent clustering percentage at 17,19 and 20 pcw. **(H)** Scheme of intersecting clusters from two different populat1i4ons (type 1 vs type 2) and associated metric. **(I)** Cluster mixing rate matrix between the 5 microglial morphological states in the ganglionic eminence at 17 pcw. **(J)** % of four microglia types vs Ramified cluster across time at 17,19 and 20 pcw.

We observed significant variability in the percentage of cells organized in clusters *versus* isolated between the different morphological types, with aggregated cells being predominantly organized in clusters with 70% of cells found in clusters (Fig. 4C-D) unlike amoeboid cells where the fraction of cells organized in clusters drops to 46% (see also fig. S24 for the other regions). We also investigated the changes over time in the fraction of clustered cells (Fig. 4E) for the 5 microglia types in the four identified anatomical regions (striatum, neocortex, cortical boundary, ganglionic eminence): we found quite homogeneous fractions of clustered cells in the ganglionic eminence (Fig. 4E), suggesting that specific classes tend to group in specific territories.

To further study the mixing of different microglial classes, we estimated the fraction of the clustered cells A in the intersecting area formed by clusters from A and an other morphological category B by the total area of these clusters (Fig. 4F). We then estimated in the ganglionic eminence the fraction of cells from one type in the clusters of the other types, as summarized in the matrix representation of Fig. 4G. Interestingly, we found that 44% of proliferative territories (not isolated cells) were mixed with aggregated territories (clusters). Also, there were no clustered phagocytic and aggregated cells in proliferative clusters and no clustered aggregated in the ramified clusters. This suggests that only certain morphological types, when distributed in clusters, can mix but others do not, a situation that should be further explored. Finally, we explored the time evolution of mixing between the different microglial types (Fig. 4H). This analysis allows to follow in time how cell clusters are overlapping. We chose ramified cells as a reference and computed the fraction of the four other microglial types located inside ramified clusters. We found that from pcw 17 to 20, the percentage of all groups inside the cluster of ramified decays below 10% (this trend is similar for the striatum Fig. S25, but not for the neocortex Fig. S26). In the cortical boundary, this ratio is also around or less than 10% (Fig. S27).

To summarise, and in all 5 morphological classes and the 4 regions we analysed across time, our findings suggest that there is no strong repulsion or attraction of clusters by other clusters. Furthermore, as development progresses, the overlap between the different classes is reduced so that each morphological group occupies a specific territory. This is consistent with brain development but not reported in humans. As the structures mature, we report here the switch to a more ramified microglia phenotype for example and this is captured by our single-cell data too (Fig. S6). From the mouse, we know that microglia are driven to occupy the space during embryonic development by proliferation but as the proliferative potential drops, contact inhibition which refers to the physio-regulatory mechanism of arresting cell division when two cells come into physical contact, may provide a mechanism through which microglia occupy space with no overlap [9]. This is consistent with the distinct morphologies occupying specific niches in humans from our data. Furthermore, the underlying neurodevelopmental landscape may drive a specific phenotype such that we see the most striking separation/non-overlap of clusters for example in the striatum (a relatively mature structure by this stage) as compared with the ganglionic eminence where there is more overlap, is a transient structure at this stage.

#### Microglial classes correlate positively with proportions obtained using scRNAseq analyses

We identified 10 clusters (Fig. S6C) and annotated microglia and other cell types according to gene lists obtained from previously published datasets (SI table Fig. S28). Our purpose here was to demonstrate whether proportions obtained from DeepCellMap within similar pcws correlated with single-cell obtained proportions of the same classes of microglia. We therefore chose the three most functionally distinct classes - homeostatic, phagocytic and proliferative. Amoeboid and aggregated microglia though morphologically distinct, may have dual functions - migratory or phagocytic/activated. Albeit it being a restricted temporal window, we are able to show with the Spearman’s correlation that the proportions of microglia calculated by DeepCellMap in 3 classes (ramified/homeostatic, phagocytic and proliferative) in the forebrain correlate positively with proportions of microglia calculated from our single-cell RNAseq data (homeostatic, phagocytic and proliferative) matched for 3 pcws weeks 11,12 and 14 (r=0.64, p=0.06) (Fig. S6E). If we take the proliferative and ramified/homeostatic, the correlation is even stronger (r = 0.85). This shows a good positive correlation between DeepCellMap and single-cell-transcriptomic proportions in two independent datasets.

### 2.3 Robust association between microglia and blood vessels in response to SARS-CoV-2 in human fetal cortex

To show the versatility of DeepCellMap, we used our pipeline in fluorescence imaging of microglia in neocortical tissues of patients infected with SARS-COV-2, labelled using antibodies against SARS-CoV-2, microglia (IBA1), blood vessels (Laminin) and lysosomal cells (CD68) (Methods). Using image processing tools (Methods), we detected and classified four types of cells, to the channels from which they are segmented: blood vessels from channel laminin (green), microglia from channel IBA1 (white), lysosomal cell from CDS68 channel (red) and phagocytic microglia from the superposition of CD68 and IBA1 channels (Fig. 5A).

**Figure 5.**
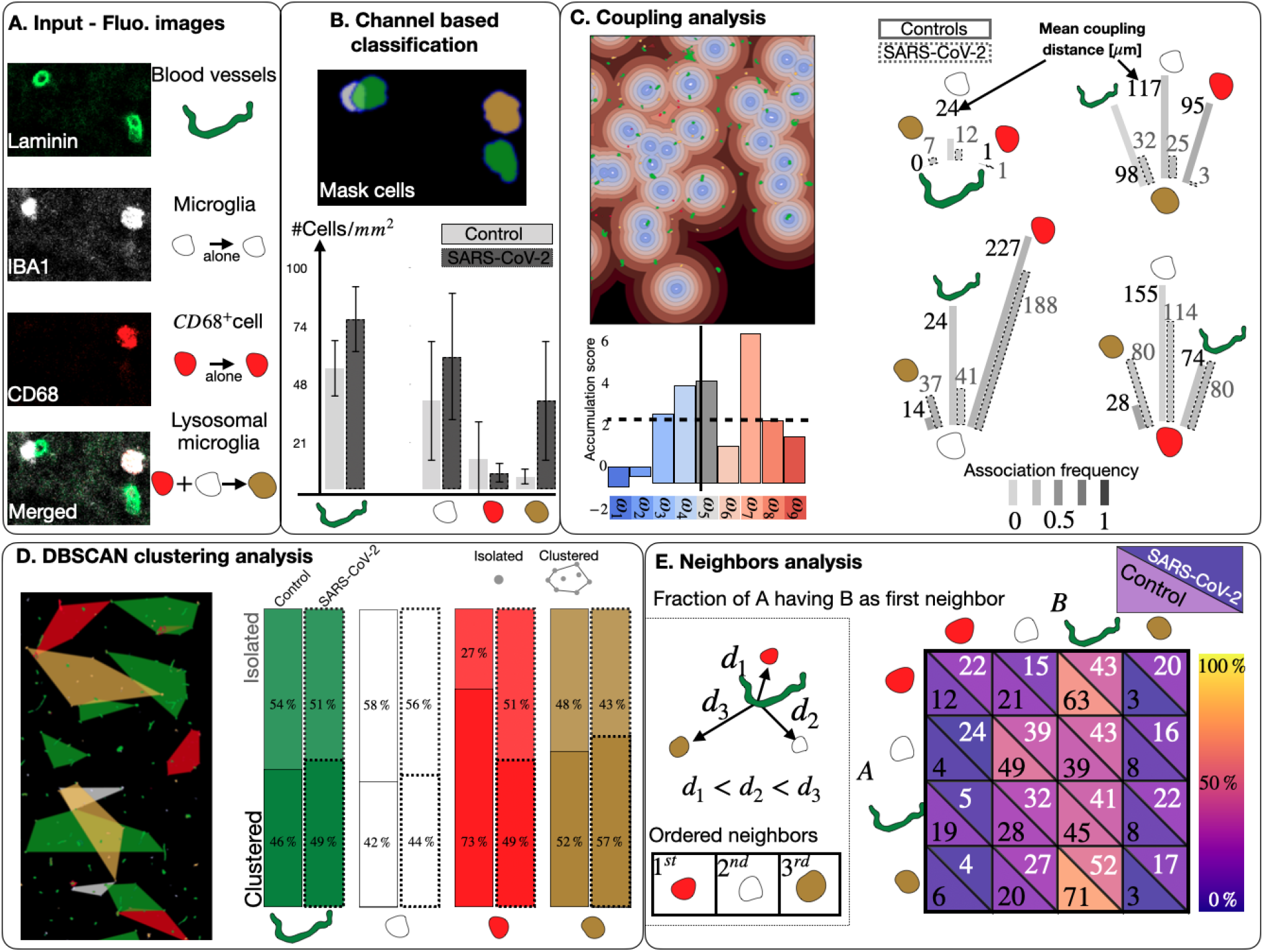
DeepCellMap applied to SARS-CoV-2 (COVID19) fetal human brain confocal data. **(A)** 3 channels : Laminin (blood vessels), IBA1 (microglia/macrophages) and CD68 (Lysosomal marker). **(B)** Cells are classified according to the channels from which they are segmented. We segmented four types of objects : blood vessels from channel Laminin (green), microglia from channel IBA1 (white), lysosomal cell from CD68 channel (red) and phagocytic microglia from the superposition of CD68 and IBA1 channels (n = 3 controls and n = 4 SARS-CoV-2 exposed sample. Error bars correspond to standard errors). **(C)** Coupling frequency and mean distances. **(D)** Fraction of isolated vs clustered cells computed from the DBSCAN algorithm. **(E)** Fraction of A cells having B cells as first neighbors. Differences between control and SARS-CoV-2 are represented in the matrix.

In fluorescence microscopy, the morphological diversity of microglia is reduced from the classes we observed in brightfield during normal neurodevelopment. IBA1 cells were mostly amoeboid/round microglial cells consistently across multiple regions including the neuroproliferative zones, the cortical plate, the subplate and the marginal zone of the telencephalic wall (Fig. 5B). We extracted cell counts for each object identified above and normalised these to tissue size to obtain a 2D density in cells/mm2. We found that in SARS-CoV-2 samples, an increased blood vessel density, phagocytic microglia/macrophages and resting microglia compared to controls which was consistent with figures obtained using manual counts in a larger dataset (*n* = 3 controls and *n* = 4 SARS-CoV-2 exposed samples), as shown in Fig. 5B [42]. Importantly, about 50% microglia were IBA1+CD68+ in SARS-CoV-2 samples, compared to 9% in controls (Fig. 5B).

To test whether any of identified cell organization departed from a uniform distribution, we then measured the spatial association between the different cells (microglia, lysosomes CD68+ cells and lysosomal microglia) and blood vessels (Fig. 5C). We observed that the distance between spatially associated cells, and between cells and blood vessels was significantly reduced in SARS-CoV-2 samples compared with controls. We then quantified the number and overlap between the different cell clusters in SARS-CoV-2 (Fig. 5D). We measured an important decrease of CD68+ cells clustering in SARS-CoV-2 samples (49 % of cells in clusters) compared with controls (73 %). We also observed a substantial overlap between lysosomal microglia (IBA1+CD68+) clusters and blood vessels. Nearest-neighbour analyses (see Methods) confirmed the tight association between lysosomal microglia and blood vessels, in both SARS-CoV-2 and controls (Fig. 5E).

Overall, the spatial statistics computed by DeepCellMap (spatial association, clusters’ overlap and nearest-neighbor analysis) all point to a stronger association of lysosomal microglia, with the vasculature in SARS-CoV-2 samples, suggesting that these microglia could be sending or receiving signals which may alter vessel integrity which has yet to be probed further as a hypothesis. Overall, DeepCellMap is a versatile tool that can be extended to fluorescence confocal images to extract robust metrics about the spatial distribution of cells in control and disease conditions.

## 3 Discussion

To investigate the spatial distribution of cells in digital pathology and extract organizational patterns, we developed a computational pipeline, available as the open-source platform DeepCellMap. This platform integrates deep learning for cell classification with advanced spatial statistics. DeepCellMap corrects for possible confusion between cell morphotypes during automatic classification, and local cell densities during statistical parameter estimations, ensuring robust analysis.

In the age of digital pathology, DeepCellMap allows the rapid extraction of information about core inflammatory processes to probe specific hypotheses. It can be adapted in clinical and scientific settings, identifying basic counts in different classes of cells, statistical associations between pairs, and cluster analyses which places it along the continuum from a basic diagnosis based on an inflammatory profile to a platform that allows the probing of robust patterns/characterisation of cell populations for scientific studies. Because of its dual potential and our expanding it to fluorescent images, it can be widely used.

### Dynamics of human brain microglia colonization

With DeepCellMap, we capture here the morphological diversity of microglia during human brain development and establish a novel, thorough, and adaptable pipeline to group them into morphological classes. We know from the rodent that the exponential expansion phase of microglia during murine development is largely postnatal and more linear prenatally following the growth of the brain [9, 43]. From our data, although restricted by the temporal windows we had investigated, we capture the decrease in proliferative microglia as development progresses. We also identify that whilst in some regions the numbers keep increasing (the striatum, the ganglionic eminence), in others (e.g. the neocortex), they undulate consistent with previous literature [5, 44, 9] suggesting that the trend of expansion and refinement of the population will vary by anatomical region.

### Microglia morphological types lay in distinct microdomains

In terms of the occupation of space by microglia, and as development progresses, the overlap between the different classes is reduced so that each morphological group occupies a specific territory. This is consistent with brain development but not clearly reported in humans. As anatomical structures mature, we report here the switch to ramified microglia for example and this is captured by our single-cell data as well. From the mouse, we know that microglia are driven to occupy the space during embryonic development by proliferation but as the proliferative potential drops, contact inhibition which refers to the physio-regulatory mechanism of arresting cell division when two cells come into physical contact, may provide a mechanism through which microglia occupy space with no overlap [9]. This is consistent with the distinct morphologies occupying specific niches in humans from our data. Furthermore, the underlying neurodevelopmental landscape may drive a specific phenotype such that we see the most striking separation/non-overlap of clusters for example in the striatum (a relatively mature structure by this stage) as compared with the ganglionic eminence where there is more overlap, is a transient structure at this stage.

### Strong association between different morphotypes is a marker of apoptosis

The strong coupling association between morphotypes such as proliferative and phagocytic/aggregated is in line with the rodent literature that microglial proliferation may be associated with increased apoptosis [45, 46]. Microglia are phagocytes and it is precisely during these temporal windows that we expect an increase in cell death as the brain prunes extranumerary cells. Dying cells influence the distribution of microglia and this is the first time that the coupling association between proliferative and phagocytic and aggregated microglia is reported in humans. We also observe an overall increase in phagocytosis genes as development progresses (single-cell data) which mirrors our histology data. Overall, DeepCellMap provides new metrics which are informative regarding the overall organisation of a very complex cell type during human development.

### Tight association of microglia with blood vessels in SARS-COV-2 infected patients

DeepCellMap revealed the association with blood vessels in SARS-CoV-2 samples and this is particularly significant for two reasons: first, SARS-CoV-2 samples exhibited weakened blood vessels due to the loss of claudin-5 tight junctions, and second, these samples contained hemorrhages [29]. The increase in microglial numbers and their association with blood vessels may suggest that microglia may either be responding to, or contributing to, changes in vascular integrity. This raises important questions about the role of microglia in areas of hemorrhage or in blood-brain barrier disruption in other diseases [47]. Specifically, are microglia aiding in the repair of weakened vessels, or are they contributing to the loss of claudin-5. This warrants further investigation.

### Conclusion and perspectives

In terms of applications, DeepCellMap applied to neurodegenerative disease tissues can be help to understand concomitant pathology (alpha synuclein, tau, TDP43) [48] to try and fully characterise associations between specific microglial classes and neuronal pathology. This can lead to the potential targeting of a specific microglial class or the class with the strongest association with the pathology for therapeutics. Furthermore, another layer may be added for those diseases that look pathologically similar but are clinically distinct, hence why a tool like DeepCellMap would be very valuable in characterising the inflammatory profile to distinguish between diseases.

From an imaging point-of-view, 3D reconstructions of microglia in thicker tissue sections would provide valuable insights into the migratory phenotype, which remains particularly challenging to define in humans. This approach would allow us to better understand the role of migration in microglial colonization during development and how this process may be altered by SARS-CoV-2 infection, further enhancing the capabilities of DeepCellMap.

By computing a wide range of statistics on the spatial relationships between different cell states or types, it becomes possible to map more complex cellular assemblies involving multiple cell types, similar to approaches used for molecular assemblies in super-resolution fluorescence microscopy [41, 49]. The development of such multivariate spatial analysis is especially relevant given the rise of spatial transcriptomics and multiplex imaging techniques [50], such as multiplex immunohistochemistry [51], imaging mass cytometry [52], and co-detection by indexing [23], which can identify dozens of cell types *in situ*. These advanced imaging and mapping techniques will ultimately enable the deconstruction of complex spatial networks of cell types within tissues, providing robust and precise predictive models of clinical outcomes or, more broadly, revealing the biological principles that govern tissue organization.

## 4 Methods

### 4.1 Immunohistochemistry data collection

We focused on the forebrain along the frontal axis which includes the cortex, the striatum, border regions and the ganglionic eminence. The striatum’s development begins at the basal telencephalon as early as the 7th pcw and spans the entire developmental period until 25-28 pcw [53, 54]. The ganglionic eminence by that stage resolves and after 30 pcw, the corpus striatum seems to have reached its mature form. Nissl and PAS-AB labellings were used for the delineation of the anatomical boundaries of these structures (Figs. 1 and S1).

#### Histological slides

We labelled and selected high-resolution histological slides across human development from the 10th pcw-term for this study. Immunohistochemistry steps were consistent with the methods below for tissue-processing for a total of 31 cases. All demographics are summarised in supplementary table 1 (Fig. S29). These cases were covered by the appropriate ethical approvals from the University of Zagreb, the Croatian Institute for Brain Research and the Oxford Brain Bank (Rec approval: 23/sc/0241, South Central Oxford C). These data including the SARS-CoV-2 dataset summarised below can be made available upon request. Exclusion criteria were congenital abnormalities, genetic disorders, brain trauma, periventricular leukomalacia, and hypoxic ischaemic encephalopathy except for the SARS-CoV-2 samples that had haemorrhages described elsewhere ([29]. Additional exclusion criteria were infection, and brain trauma.

#### Immunohistochemistry

Paraffin-embedded blocks were cut into thin sections of 10 *µ*m on a microtome for immunohistochemistry. Brightfield immunohistochemistry experiments were performed using antibodies against microglia with the following dilutions: rabbit (019-19741, Wako) or mouse (ab283319, Abcam); mouse anti-Ki67 (GA62661, Dako Omnis, Agilent), rabbit TMEM119 at 1:500 (ab185333, Abcam, mouse PG-M1 at 1:400 (GA61361-2, Dako Omnis, Agilent), mouse P2RY12 at 1:1000 (ab180366, Abcam) and rabbit SOX2 at 1:1000 (sc365823, Santa Cruz). The first step was deparaffinization of formalin-fixed paraffin embedded sections in xylol solution and rehydration in descending concentrations of diluted ethanol (100%, 96%, 70%). Antigen retrieval was done by heat induced epitope opening using citric acid buffer (pH = 6.2). Thereafter, sections were pre-treated with methanol and hydrogen peroxide to block endogenous peroxidase and phosphatase activity. Sections were blocked with a solution of 5% Bovine serum albumin + Tween20 (0.1%) + normal horse serum (5%) in 1X PBS and then incubated with primary antibodies overnight. The next day, secondary antibodies were applied using either the Immunopress duet kit (MP7724, Vector labs, UK) with anti-mouse epitopes visualised with DAB chromogen in brown and anti-rabbit epitopes visualised with alkaline phosphatase in magenta or the Envision kit (mouse/rabbit) (K500711-2, Agilent, UK). Sections were counterstained with haematoxylin or methyl green and coverslipped with permanent mounting medium before imaging.

#### Brightfield slide-scanning

Imaging was done using high-resolution histological slide scanners: the Aperio Imagescope (Oxford, UK) and Hamamatsu Nanozoomer (Zagreb, Croatia) 2.0 RS with a 40× objective (numerical aperture of the lens = 0.75) at 0.45 *µ*m × 0.45 *µ*m pixel resolution.

#### Microglial morphologies selected

We sampled microglial morphologies based on previously published literature during human development [31, 32, 5, 7, 33, 34, 30, 35, 55]. In brief, microglial morphologies described here were proliferative, amoeboid, aggregated, phagocytic, ramified (Fig. S1).

For the coupling of morphology with function, we refer to the representative examples provided in fig. S6A. The dataset consists of a set of 24 histological images of human fetuses at various stages of brain development. Images were processed coronally through the frontal axis of the brain from 10 post-conceptional weeks until term. All sections were histochemically labelled with haematoxylin and eosin (H&E) according to standard methods and assessed by a neuropathologist for histology.

### 4.2 Fluorescence data collection

#### Human tissues

We extended our brightfield pipeline to allow the processing of human fetal images using fluorescence microscopy. Demographics of the cases are provided in supplementary table Fig.S29. Ethical approval was granted by the Human Developmental Biology Resource (HDBR) jointly funded by the MRC and Wellcome Trust (Rec number (Newcastle): 23/NE/0135, Newcastle and North Tyneside ethics committee and Rec number (London): 23/LO/0312, Fullham ethics committee).

Human fetal tissues were obtained from the Human Development Biology Resource (HDBR), provided by the Joint MRC/Wellcome Trust (grant #MR/R006237/1). The HDBR provided fresh tissue from fetuses aged 9–21 post-conception weeks (pcw): we selected here from 26 haemorrhagic samples, 7 samples with elective terminations (3 with no histological abnormality recorded and 4 with cortical haemorrhages whose mothers were positive for SARS-CoV-2) for the purposes of demonstrating how DeepCellMap can be applied to these tissues. The processing of these samples has been specified elsewhere [1]. Briefly, all tissues were fixed for at least 24hrs at 4°C in 4% (wt/vol) paraformaldehyde (PFA) in 120 mM phosphate buffer (pH 7.4). Brains were then sucrose-treated (15 and 30% sucrose solution sequentially for 24 h each), OCT-embedded, and then 20 *µ*m thick sections were cut using a cryostat.

#### Immunofluorescence

Sections were then stained in a solution containing 10% BSA and 0.1% Triton, using the primary antibodies CD68 (1:100 DAKO M0814), IBA-1 (1:100 Abcam ab5076), pan-laminin (1:100 Sigma L9393), rabbit IgG isotype control (1:50 Abcam ab172730 [EPR25A]), SARS-CoV-2 spike protein (1:100-250 Genentech GTX632604 [1A9]) and SARS-CoV-2 nucleocapsid protein (1:50-150 Sino Biological 40143-R001). Life technologies secondary antibodies, used at 1:1000, were donkey-anti-goat Alexa fluor 488 (A11055), anti-mouse Alexa fluor 568 (A10037) and 555 (A31570), anti-rabbit Alexa fluor 647 (A31573) and 488 (A21206), and goat anti-rat 555 (A21434). Sections were all stained with DAPI (Sigma D9542) and mounted in Mowiol (Merck Biosciences).

##### 4.2.1 Fluorescence data and confocal imaging

Highly-resolved confocal imaging was subsequently performed using a Zeiss LSM 800 inverted microscope and a Zeiss Plan-Apochromat 20 × 0.8 objective, or a Zeiss AxioScan slide scanner and a Zeiss Plan-Apochromat 20 × 0.8 M27 objective at 0.312 um/pixel as resolution. More details have been specified elsewhere [29]. We illustrate our image processing pipeline in Fig. 1.

### 4.3 Single-cell RNA-seq analysis of an independent dataset

We analysed a total of 6584 microglial cell transcriptomes from the forebrain, midbrain and hindbrains and focused our analyses on the forebrain, diencephalon and telencephalon regions for the purposes of this work. These data were derived from [56]. The temporal windows considered were 10-15 pcw or the late first/early second trimester matching part of our histological temporal windows. Normalization and dimensionality reduction were done using Sctransform (version 0.4.0) and RunPCA function of Seurat R package (version 4.1.4) with default parameters. To harmonize the data, we used harmony R package (version 1.0.3) to regress out the effect of each sample with *max*_*iter*_ set to 20 and the first 30 principle components. The Uniform Manifold Approximation and Projection (UMAP) plots were made using RunUMAP function of Seurat R package with dimensions set to the first 20 and using the harmony-derived dimensionality reduction. The number of dimensions for UMAP plots were defined using Elbowplot function implemented in Seurat R package. The scRNAseq clustering was done using FindNeighbors (k.param=15 and dims = 1:20) and FindClusters (resolution = 0.5, algorithm = 2) functions of Seurat R package. The resolution was defined using the *Clustree* R package (version 0.5.0). Annotation of clusters was done based on canonical and functional markers from [57, 58, 59, 60, 61, 62], see SI table 1 Fig. S28. We then calculated the proportions of 3 microglial classes extracted from our single-cell analysis and correlated these with the proportions of the same classes obtained using DeepCellMap matched for pcw(s) 10 to 14 (see Fig S6B-D).

### 4.4 Image processing

#### Pre-processing

Images were manipulated at different scales. First, tissue was extracted using Otsu thresholding and morphological operations on the downscaled image (factor 1/32 for brightfield and 1 for fluorescence data). A subdivision of the original images into tiles of size 1024×1024 pixels characterised by their row and column numbers allowed a manipulation of the regions of interest at high resolution (Fig. S3). Thus, each region of interest is characterized by its coordinates (top left tile and bottom right) in the image grid. Cells are identified by the quadruplet (*row, col, x, y*) describing the tile (*row, col*) to which their centre of mass belongs and their coordinates (*x, y*) in that tile.

#### Cell detection and segmentation

Given the arbitrary division of the image into tiles and the possibility that cells may be present on several tiles, the following procedure is applied to ensure that all cells are taken into account and not counted more than once. During the detection procedure on a tile, the 8 neighboring tiles are considered and the segmentation algorithm is applied to the resulting 3×3 tile image. The centers of mass of detected cells with part of their body in the central tile are calculated; if this center of mass is in the central tile, then the cell belongs to this tile. The segmentation algorithm proceeds in several steps, applied to each tile whose tissue percentage exceeds 5% (Fig. S4A). First, *Image binarization* is performed with the Otsu adaptive thresholding from the eosin color-deconvolution of the RGB image [63]. The algorithm evaluates the histogram of the eosin image and finds two critical thresholds allowing to separate three regions. Pixels belonging to the third region are assigned a value of 1, while the others are set to 0 (Fig. S4B1). Small isolated fragments of the binary mask are removed by using a disk of radius *r* = 1. (Fig. S4B2). Then, a *morphological dilation* with a disk of radius *r* = 3 is applied to the binary mask to connect the different binary masks belonging to the same cells (Fig. S4B3). *Holes in the cell mask are filled* by a background reconstruction (Fig. S4B4), and finally, a *size filter* is applied to cell masks, and cells with a size smaller than a threshold *T* = 700 pixels) are filtered out (Fig. S4B5).

During this segmentation procedure, two parameters can be adjusted by the user: the radius *r* of the dilation disk (between 2 and 4), and the maximum size of the cells, which depends mainly on the resolution of the image.

The output is a binary mask of segmented cells over background. Five parameters are associated with each segmented cell: row and column of the corresponding tile, coordinates (*x, y*) of the cell centre of mass, and cell size (sum of mask pixels equal to 1). For brightfield images, the result of the DL classification into the different types of microglial cells is also listed.

#### Validation of cell detection

To validate how DeepCellMap detects microglial cells in brightfield imaging, we randomly selected *N* = 20 tiles of size 3000×3000 from the dataset. For each tile *N*_*i*_, a certified expert counted the number 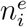 of microglial cells. This number was then compared to the number 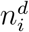 of cells automatically detected by DeepCellMap (Fig. S5. We computed the overall error by the following formula:

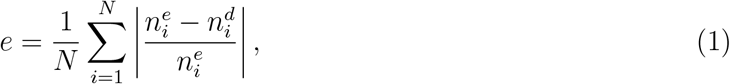

and obtained an overall error *e*_*mean*_ = 15 ≤with std error *e*_*std*_ = 2.4 ≤over *N* = 20 annotated tiles.

Since the detection algorithm is sensitive to size filtering of detected cells, the ‘Laboratory cell segmentation’ notebook has been created to calibrate this critical value as best as possible according to the dataset.

### 4.5 Deep Learning classification of microglial cells

To ensure that the training database of annotations in each class covers a maximum of the inter-class spectrum, cells are randomly selected from images at different times: further several tiles are also randomly selected, cells are detected on a tile, and a fraction of these cells is added to a database of unlabeled cells. Cells are patches (size 256^2^) whose center is the cell’s center of mass.

An ergonomic annotation tool allowed the expert to label each cell belonging to one of the five morphological states. This made possible to build up a base of annotations in each class. We illustrated in Fig. S30 this dataset using the ramified microglia cell type.

The RGB images of the cells and the labels assigned to the cells were given as input to the UMAP dimension reduction algorithm in order to visualise the inter-class heterogeneity of the cells in the training database and to manually correct annotation errors (Fig. S8). After the training set has been built up, the masks contained only one cell in the centre, but other cells may be present in the periphery. These cells were added to the mask with an “other” label (Fig. S9) to prevent them from being considered as background by the model. RGB images and completed masks were then shuffled and split into training (75%), validation (15%), and test (15%) sets. The test set was used for the evaluation of the final model.

During training, images belonging to the training set were submitted to image augmentation procedures (brightness changes, flips, crops, and rotations) and were applied to each image to expand data diversity and make the model more robust [64].

A convolutional neural network (CNN) based on U-Net architecture was selected for cell classification. The network consisted of a contracting and an expansive path. The contracting path consists of three applications of the following steps : two 3×3 convolutions (with zero padding), each followed by a rectified linear unit (ReLU) and a 2×2 max pooling operation with stride 2 for down sampling. For the fourth layer of the contracting path, a dropout of rate 0.5 was added between the 2 convolutions and the max pooling operation to constrain the fully connected layers and to reduce overfitting. Then two 3×3 convolutions (same configuration with ReLu activation and zero padding) are applied before a dropout (rate 0.5) that ends the contracting path. Every step in the expansive path consists of a 2×2 up-sampling of the feature map followed by a 2×2 convolution (“up-convolution”) that halves the number of feature channels, a concatenation with the correspondingly cropped feature map from the contracting path, and two 3×3 convolutions, each followed by a ReLU activation. At the final layer a 1×1 convolution is used to map each 64-component feature vector to the classes we identified: background, proliferative, amoeboid, aggregated, phagocytic, ramified, and detected.

“Adam” optimizer was used for optimization, the initial learning rate was set to 0.003, batch size was set to 3. The CNN was trained on an off-the-shelf NVIDIA GeForce GTX 1080 with 8 GB GPU memory, for 40 epochs, training time took about 10 h.

For each pixel of the image, the output of the CNN algorithm is the probability that this pixel belongs to one of the 5 pre-defined classes of microglial cells (Fig. S10B3). The overall probabilities of segmented cells were obtained by averaging the probabilities of mask’s pixels. There are some aggregated cells (*<* 1%) whose cell body was larger than the patch, in this case the mask probabilities of the cell in the patch were assigned to the full mask of the cell. Finally, microglial morphology was assigned by selecting the label for which the maximum probability was achieved.

### 4.6 Automatic delineation of tissue regions

To identify the components of the tissue with similar nuclei densities, we first segmented automatically cells’ nuclei. In fluorescence microscopy, we used the color channel corresponding to nuclei labeling. In brightfield microscopy, the identification of nuclei is less obvious and we tuned CellPose algorithm [16]) to segment the nuclei contours in different types of images (Fig. 2A). The output of the algorithm is the number of nuclei present in each image patch of size 256×256, allowing to reconstruct the heatmap of the underlying tissue density. We focused on 4 regions: striatum, neocortex, ganglionic eminence, and the cortical boundary (Fig. S16 and Fig. S17).

For each tissue, the histogram of nuclei densities was then separated into 3 or 4 regions using the Otsu-multi-thresholding algorithm, which optimises the choice of thresholds. The different tissue classes obtained with Otsu thresholding then underwent several elementary morphological operations to distinguish each of the physiological regions identified in the images: striatum, neocortex, ganglionic eminence, and the cortical boundary (Fig. S31).The cortical boundary and the neocortex (which cover regions several hundred thousand pixels wide in the images), were subdivided into 4 sub-regions in which the calculations were carried out and recombined by weighting the results as described in figure S31.

### 4.7 Level set analysis of cell-to-cell spatial association

To measure the spatial association between different populations of cells (or cells to regions) and account for the uncertainties associated with the deep-learning classification of cell types, we implemented a generalised version of the statistical approach developed in [40].

The method can be described as follows: we first specify K cell types **A** = {*A*_1_, …, *A*_*k*_, … *A*_*K*_. During the deep learning classification, each cell type *A*_*k*_ (1 ≤ *k* ≤ *K*), is classified as type *A*_*j*_ with probability *p*_*jk*_. We highlight that these classification probabilities can be estimated from the results of the deep-learning classification on training dataset.

To compute the potential accumulation of *A*_*m*_ cells around *A*_*l*_ cells, we first select the cells with a maximum classification probability in the type *A*_*l*_ and map the domain of interest Ω (typically a predefined region-of-interest within the tissue slide) around the population *A*_*l*_ with a series of levelset regions ⋃ _1≤*i*≤*n*_ *ω*_*i*_, with *ω*_*i*_ = {**x** ∈ Ω|*r*_*i*_ ≤ |**x** − ∂*A*_*l*_| *< r*_*i*+1_} which is the region of Ω that contains points **x** at a distance comprised between *r*_*i*_ and *r*_*i*+1_ (*r*_1_ = 0 *< r*_2_ *<* … *< r*_*n*_) from the contour ∂*A*_*l*_ of *A*_*l*_ cells.

To measure the spatial association of type *A*_*m*_ of cells to type *A*_*l*_, we use the ensemble of cell coupling (center of mass typically) {*u*_*j*_}_1≤*j*≤*N*_, with 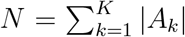 the total number of cells, and the probabilities 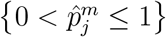 of being classified as a *A*_*m*_ type, and measure the total number of *A*_*m*_ points within the levelset *ω*_*i*_ according to the estimator:

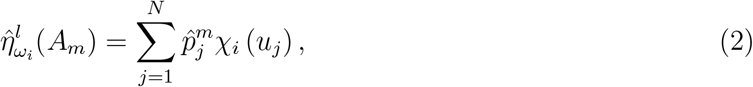

where the indicator function *χ*_*i*_ (*u*_*j*_) = 1 if *u*_*j*_ ∈ *ω*_*i*_, and 0 otherwise.

#### Correcting for cell misclassification

Automatic cell classification leads to potential errors, and a correction of the statistical analysis of cell-to-cell association is thus needed to account for misclassification: indeed, in the worst case scenario where cells of type *A*_*m*_ would be misclassified as type *A*_*n*_, the association of *A*_*m*_ cells to *A*_*l*_ cells would actually reflects the association of *A*_*n*_ cells to *A*_*l*_ cells.

To derive the correction term, we consider the *K × K* confusion probability matrix **P**, with elements (*p*_*nm*_)_1≤*n*,*m*≤*K*_ the probabilities for a *A*_*m*_ cell to be classified as a *A*_*n*_ cell Since each *A*_*m*_ cell within level set *ω*_*i*_ has a probability *p*_*nm*_ to be classified as a *A*_*n*_ cell, the measured accumulation 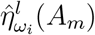 of *A*^*m*^ cells within *ω*_*i*_ around *A*_*l*_ cells is equal to

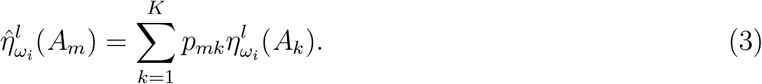

with 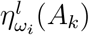 the *ground-truth* accumulation of *A*_*k*_ cells in *ω*_*i*_ around *A*_*l*_ cells. Note that we have *η*^*l*^ (*ω*_*i*_(*A*_*l*_)) = 0 because we are estimating the association to *A*^*l*^ cells.

If 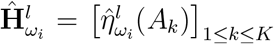 and 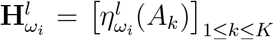 are respectively the vectors of measured and *ground-truth* number of cells within levelset *ω*_*i*_ around *A*_*l*_ cells, we can rewrite the equation in a matrix form as:

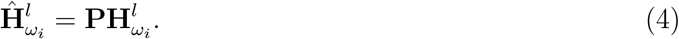

and we can invert the *ground-truth* which can be estimated from the measured accumulation of cells based on the relation

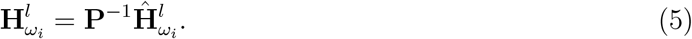

Based on the previous estimation of the accumulation of the different cell types in levelsets around a given population *A*_*l*_ of cells, we aimed at determining the cell types that significantly accumulate in each level set. Such estimation will allow us to characterize the spatial association between the different types of cells.

To assess whether or not the accumulation of *A*_*m*_ cells in levelsets *ω*_*i*_, 1 ≤*i* ≤*n* around *A*_*l*_ cells is significant, we followed the methodology given in [65] and computed the *n*-dimensional Ripley vector

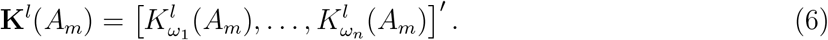

With

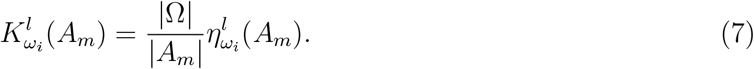

that we rewrite

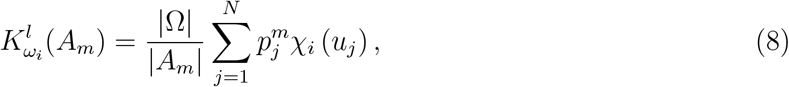

where 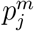 is the probability that cell position *u*_*j*_ belongs to the *A*_*m*_ population of cells. It can be computed from the classification probabilities 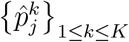 and the confusion matrix **P**:

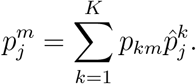

The estimated total number of *A*_*k*_ cells is given by

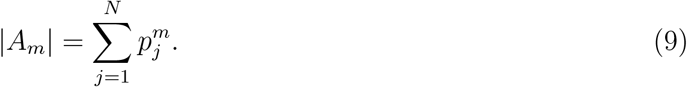

When cells *A*_*m*_ are coupled to cells *A*_*l*_, we expect a significant accumulation of the *A*_*m*_ cells’ positions in the neighborhood of *A*_*l*_, *i*.*e*. in a subset of the levelset regions *ω*_*i*_ for 1 ≤ *i* ≤ *n*. To rule-out a fortuitous accumulation of *B* cells due to chance, it is necessary to characterize the distribution of **K**^*l*^(*A*_*m*_) under the null hypothesis of complete spatial randomness where the *A*_*m*_ cell coordinates would be randomly and uniformly distributed over Ω according to the Homogeneous Poisson law. Under the null hypothesis of *A*_*m*_ randomness, each function 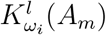 is normally distributed [40]

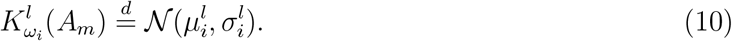

To compute the mean 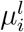 and standard deviation 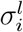,we highlight that, if *A*_*m*_ cells are randomly distributed, *χ*_*i*_(*u*) is a Bernoulli variable with mean

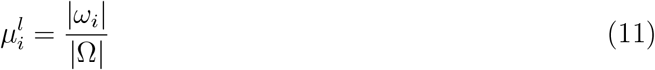

and variance

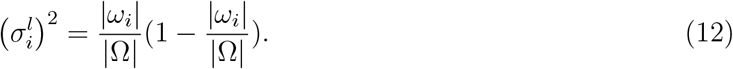

Therefore, we obtained that the mean is given

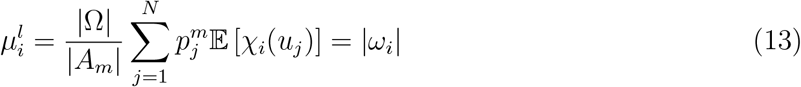

and the variance is given by

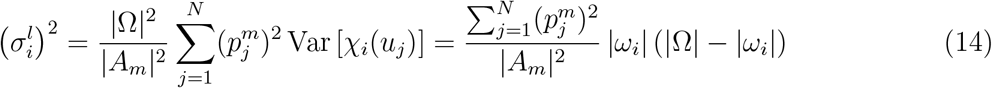

Under the null hypothesis, the vector **K**^*l*^(*A*_*m*_) is Gaussian:

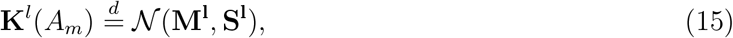

with the mean 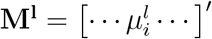,and the covariance matrix is

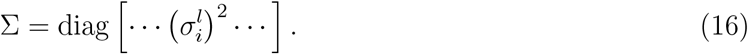

The accumulation of cells within the levelset region *ω*_*i*_ is statistically significant when the mean exceeds a threshold

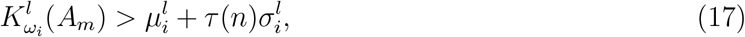

where 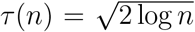 is the universal statistical threshold used for determining the relevant signal in a vector containing Gaussian noise [66].

From previous statistical thresholding (eq. 17), we can now estimate the subset of level set regions 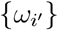 where the observed accumulation of *A*_*m*_ cells is statistically significant.

To quantitatively describe the association of the set of cells *A*_*m*_ to the set *A*_*l*_, we computed the number 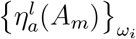 of spatially associated *A*_*m*_ cells inside each level set *ω*_*i*_:

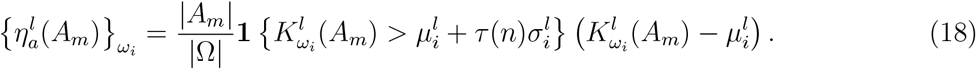

which corresponds to the significant overcount of *A*_*m*_ cells inside *ω*_*i*_ above the expected number of randomly distributed cells. We then defined a probabilistic association index p-ASI which corresponds to the ratio of *A*_*m*_ cells that are spatially associated to *A*_*l*_ cells:

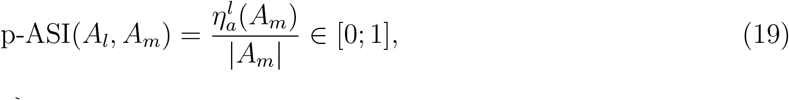

with 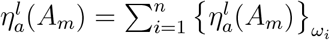 the total number of associated *A*_*m*_ cells.

We also derived the probability-weighted coupling distance

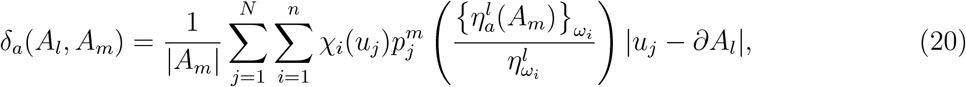

where the ratio 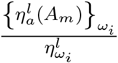 is the proportion of *A*_*m*_ cells within *ω*_*i*_ that are spatially associated to *A*_*l*_ cells. It is the probability that cell *u*_*j*_ is spatially associated and not randomly distributed, and |*u*_*j*_− ∂*A*_*l*_ |is the Euclidean distance of *u*_*j*_ to *A*_*l*_ cells’ contours. Finally, the association p-value is given by

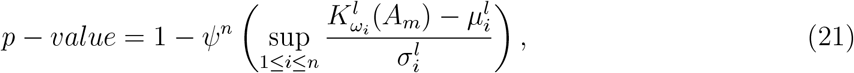

where *ψ*(.) is the cumulative density function of a normalized Gaussian random variable.

#### Validation levelset analysis with synthetic simulations

To validate the accuracy of DeepCellMap in characterizing the spatial association between two cell types, we generated synthetic simulations with various spatial distributions. For a sake of simplicity, we considered three cell types: A, B and C, and cells were reduced to points. A positions were uniformly distributed in a square domain Ω. Among the *B* and *C* positions, a part 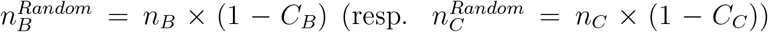 of cells are randomly distributed in Ω, while the remaining part 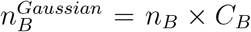 (resp.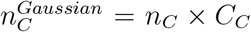) are positioned around A cells with a Gaussian association distance *d* ∼ *N* (*µ*_*B*_, *σ*_*B*_) (resp. *d* ∼ 𝒩 (*µ*_*C*_, *σ*_*C*_). To efficiently sample the coupled *B* and *C* cells, the distance transform from

A cells is computed across the entire image, a random association distance *d* is drawn from𝒩 (*µ, σ*), and a coordinate in the distance map at distance *d* from A cells is randomly selected (fig. S11). To assess the capability of DeepCellMap to handle cell misclassification, we used three different confusion scenarios: scenario P1 (no confusion), there are no classification errors, which allows the validation of the analysis of spatial association in a setting where there is no ambiguity in cell types; scenario P2 (intermediate) were 30% of *B* cells are classified as *C* cells and *vice-versa*; scenario P3 (high) where classification errors reach 45% of classification errors between types B and C. Parameters used for the simulations are summarized in the supplementary material.

The accuracy of DeepCellMap was measured by comparing the estimated association with 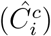 and without 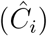 correcting for misclassification, with ground truth *C*_*i*_ between cells of type *i* ∈ [*B, C*] and cells A. The estimated association distances 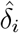 and 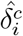 were compared to *µ*_*i*_.

### 4.8 Characterizing cell clustering and overlap with a generalized DBSCAN algorithm

To measure the rate of clustering of the different cell types, and the overlap between the domains occupied by the different cell populations, we developed an improved and automatically-calibrated *density-based spatial clustering of applications with noise* (DBSCAN) algorithm [27]. This widely used clustering algorithm contains two user-defined parameters: the radius *E* to define the neighborhood of each cell and the number *Min*_*Sample*_ which defines the minimum number of neighboring cells (points) in this neighborhood in order to consider the cell as belonging to a cluster. We used the DBSCAN algorithm with cell coordinates (*x, y*) for each microglial state (Fig. 4A) and automatically compute *E* to maximize the number of clusters (Fig. 4B):

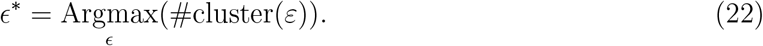

We observed empirically that the number of clusters #cluster(*ε*) has a single maximum (Fig. 4C), ensuring that our selection procedure converges. We fixed the minimum number *Min*_*Sample*_ = 4 of points inside a cell neighborhood (disk) classify it as *clustered*.

To ensure the stability of computed clusters, we defined the following criteria: after removing 10% of the cells located on the cluster’s convex hull [67], we computed the ratio between the area of the remaining cluster and the initial one, and kept the cluster if this ratio exceeds a threshold *t* = 60% (average over 100 realization). After having sorted each cell as part of a cluster or not (isolated) we computed several metrics to quantify cell clustering and the relative overlap between the clusters from different types: the fraction of clustered cells (*versus* isolated) (Fig. 4F), the mixing proportion φ_*A/B*_ between clusters of cells from type A and B. This mixing proportion is computed as the fraction of A clustered cells belonging to the convex hull of B clusters (Fig. 4H):

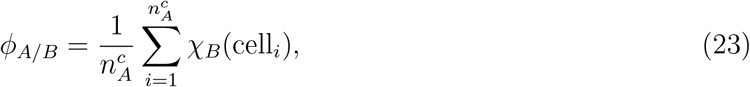

where 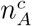 is the number of *A* clustered cells and *χ*_*B*_(cell_*i*_) = 1 if cell_*i*_ ∈ *B* clusters convex hull and 0 otherwise.

### Cell neighbors analysis

DeepCellMap allows for a traditional quantification of the relationships between each cell’s neighbors. We used various parameters to estimate the relative position of different cell populations in a region of interest:

1. For cells of type A, we define the first neighbor distance as

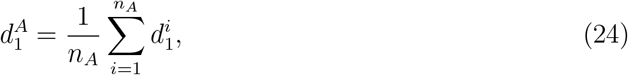

where *n*_*A*_ is the total number of cells A and 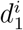 are the distances to the first neighbors of cell i (regardless of neighbor type).
2. For A cells, we define the distance to the first B neighbor as

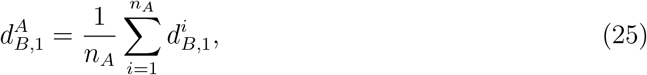

where 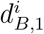 is the distance to the first *B* neighbor of cell *i*. This distance 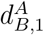 can be interpreted as the average distances from one population to another. For these last two metrics, DeepCellMap includes the possibility to calculate 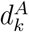 and 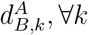.

## Supporting information

main

## 5 Ethics

The appropriate ethical approvals were granted by the School of Medicine, University of Zagreb, and are part of the Zagreb Brain Collection. The study was conducted according to the guidelines of the Declaration of Helsinki, and approved by the Ethics Committee of School of Medicine University of Zagreb (protocol number 380-59-10106-15-168/314). Part of the human fetal material for the SARS-CoV-2 samples was provided by the Joint MRC/Wellcome Trust grant #099175/Z/12/Z Human Developmental Biology Resource. Ethical approvals were also from the Oxford Brain Bank (Rec approval: 23/sc/0241, South Central Oxford C).

## 6 Data and Code availability

The software and data are available in Zenodo of the Holcman’s lab https://doi.org/10.5281/zenodo.140035 and www.Bionewmetrics.org.

## Acknowledgments

D.H. research is supported by a grant ANR AstroXite-22-CE16-0027, ANR 10LabX54 Memolife, ANR AnalysisSpectralEEG and the European Research Council (ERC) under the European Union’s Horizon 2020 research and innovation program (grant agreement No 882673). D.A.M. was supported by a Springboard grant funded by the British Council (grant agreement No 1170803491) and T.L.’s research is partially supported by grants from France-BioImaging infrastructure (ANR-10-INBS-04) and the Labex IBEID (ANR-10-LABX-62-IBEID). Experimental work was supported by K.L. who is supported by a Medical Research Council career development award (MR/S025065/1) at the MRC Centre for Neurodevelopmental Disorders, KCL, London, UK and the Croatian Institute for Brain Research and the Zagreb brain collection. We would like to acknowledge Dr Ahmed Osman at the Karolinska Institute, and Dr Jenny Dewing-Burton at the University of Southampton.We acknowledge the Oxford Brain Bank, supported by the Brains for Dementia Research (BDR) (Alzheimer Society and Alzheimer Research UK), and the NIHR Oxford Biomedical Research Centre. We would also like to acknowledge Mr Sam Perochon for his guidance on code development as well as Miss Nita Alpin and Miss Abigail Phillips for their technical assistance.

## Author contributions

D.H. co-supervised T.P. with D.A.M and T.L. D.H., T.L. and D.A.M. designed the overall project.D.A.M., L.P. and Z.K. collected fetal human tissues, labelled them with immunohistochemistry and digitised them for computational analyses. K.L. and M.M. provided the SARS-CoV-2 dataset. X.L. and E.M. performed the sc-RNA-seq analysis. Y.R., K.B. and A.L.R. analysed datasets for further profiling including spatial transcriptomic datasets. T.P wrote the code. T.P., T.L and D.H. developed the method and analyzed the data. All authors wrote and revised the manuscript extensively. Authorship was granted as per the guidelines provided by the journal.

