## Supplementary material for "Unraveling Microglial Spatial Organization in the Developing Human Brain with DeepCellMap, a Deep Learning Approach Coupled to Spatial Statistics": main

October 28, 2024

The supplementary material presents a glossary, further details about image processing, demographic details and the supplementary figures.

#### 1 Glossary

1. **SARS-CoV-2-labeling:** Antibody labelling of the COVID-19 virus.
2. **Global cell density:** the overall density of cells in the image (including all tiles).
3. **SODA:** Statistical Object Distance Analysis: Null-hypothesis statistical method to characterize the cell-cell spatial association [**?**].
4. **Level set-based analysis:** procedure to quantify cell-cell and cell-region spatial association [**?**]. The contour of a given set of cells, or the boundary of a region, is mapped with a level set function (the contour/boundary corresponding to the 0-level of the level

*[1] Group of Data Modeling and Computational Biology, IBENS, Ecole Normale Supérieure, France, [2] Croatian Institute for Brain Research, University of Zagreb, Croatia, [3] Centre for Developmental Neurobiology, MRC Centre for Neurodevelopmental Disorders, King's College London, UK, [4] Department of Women's & Children's Health, Karolinska Institutet, Sweden, [5] Department of Neuropathology & The Queen's College, University of Oxford, UK, [6] Institut Pasteur, Université Paris Cité, CNRS UMR3691, BioImage Analysis Unit, France, [7] DAMPT, University Of Cambridge, DAMPT and Churchill College CB30DS, United Kingdom. *Equal Contribution*

set function). The domain (e.g. tissue) around the segmented contour is segmented in regions that correspond to increasing intervals of values of the level set function.
Finally, the number of cells (points) from another set is determined within each region, and a null-hypothesis framework allows to reject the null hypothesis of cell randomly distributed and to determine the statistical association of cells.

5. **Cell-to-cell association (coupling):** this terminology is associated to the result of the level set-based analysis on two given cell type: the procedure determines the accumulation of one cell type around another by decomposing the image of the tissue in level sets. Two types of cells are considered to associated if the accumulation of one with respect to the other in a subset of level set regions is significantly larger than expected for a random cell distribution.

6. **Cell-to-region association (coupling):** This coupling refers to the accumulation of cells in the neighborhood (level set regions) of a region's boundary.

### 2 Image processing

We provide here several specifications associated to the image processing step.

**Tuning *CellPose* algorithm** We extracted the optimal parameters of CellPose algorithm, by testing them one-by-one, and we compared them on 10 heterogeneous tissue regions representative of different tissue environments until a qualitatively optimal configuration was obtained. We recall that there are 5 parameters:

1. Nuclei model_type

2. Approximated diameter "diameter"

3. Channels (blue)

4. Normalisation

5. Average over several models net_avg,

Once the best model had been selected, the algorithm was applied to the entire image on crops of size 256x256.

**Validating the analysis of cell-cell association: parameters of the simulations:** To validate the accuracy of DeepCellMap in characterizing cell-cell spatial association for the different cell types, we generated various spatial distributions. We considered three cell types: A, B and C. The first type of cells (A) was uniformly distributed over a square domain. Then the cells B and C were then positioned according to a labeling index and to given distances resulting in spatial association (fig. **??**). We mapped the simulation domain with level sets

around A cells, and analyzed the spatial association of B and C cells within the regions between successive level sets.
To simulate the possible confusion between the different cell types during deep-learning classification, we used three scenarios: P1-no confusion, P2-intermediate and P3-high confusion: in scenario P1, there are no classification errors, which allows validation of the algorithm in a setting where there is no ambiguity in cell types. In Scenario P2, we generated 30% of classification errors between types B and C. Finally, in scenario P3: there are 45% of classification errors between types B and C. This is summarized by the three confusion matrices:

$P_1 =$

| | $B^{GT}$ | $C^{GT}$ |
|---|---|---|
| $B^{Pred}$ | 1 | 0 |
| $C^{Pred}$ | 0 | 1 |

, $P_2 =$

| | $B^{GT}$ | $C^{GT}$ |
|---|---|---|
| $B^{Pred}$ | 0.7 | 0.3 |
| $C^{Pred}$ | 0.3 | 0.7 |

, $P_3 =$

| | $B^{GT}$ | $C^{GT}$ |
|---|---|---|
| $B^{Pred}$ | 0.55 | 0.45 |
| $C^{Pred}$ | 0.45 | 0.55 |

.

Parameters used to generate the synthetic datasets are:

1. **Image height, width:** 10000,10000.
2. **Cell type A:** Type of cell chosen to generate the cell masks (*Aggregated* in the simulations).
3. **Numbers of $n_A$ of A, $n_B$ of B, and $n_C$ of C cells**: 20, 1000, 1000.
4. **Coupling $C_B$ of B and $C_C$ of C cells around A cells.** $C_B = 0.4$ means that 40% of the cells are spatially associated to A cells, *i.e.* normally distributed $\mathcal{N}(\mu, \sigma)$ around A cells.
5. **Coupling distance $\mu_B$ (resp. $\mu_C$) of B (resp. C) cells to A cells:** Mean of the Gaussian distance of between A and associated B (resp. C) cells.
6. **Standard deviation $\sigma_B$ (resp. $\sigma_C$) of the distribution of B (resp. C):** standard deviation of the Gaussian distance between associated cells.
7. **Confusion matrix P: of the model classifying B and C cells.**
8. **Number n of images** created in each configuration: 5.

For each generated image, the outputs are

1. Mask of the cells A, B and C (see also fig**??**.A5)
2. Table of the cells with columns: **'idcell'** (cell identifier), **'tile row'** (tile row of the cell), **'tilecol'** (tile column of the cell), **'xtile'** (x-coordinate in the tile), **'ytile'** (y-coordinate in the tile), **'ximg'** (x-coordinate in the entire image), **'yimg'** (y-coordinate in the entire image), **'xtileborder'** (x-coordinate in the entire image + border), **'ytileborder'** (y-coordinate in the entire image + border), **'size'** (sum of the pixels of the cell mask), **'lengthmax'** (max length of the cell mask), **'celltype'** (ground truth), **'proba_A'** (probability that the cell belongs to A), **'proba_B'** (probability that the cell belongs to B), **'proba_C'** (probability that the cell belongs to C).

**Cluster metrics:** For each microglial morphology, we computed several metrics to quantify cell clustering and the relative position of clusters. The different metric parameters are:

1. The fraction of clustered cells

2. The mixing proportion $\phi_{A/B}$ of A clusters within B clusters. It is computed as the fraction of A clustered cells belonging to the convex hull of B clusters according to formula:

$$\phi_{A/B} = \frac{1}{n_A^c} \sum_{i=1}^{n_A^c} \chi_B(\mathrm{cell}_i), \tag{1}$$

   where $n_A^c$ is the number of $A$ clustered cells and $\chi_B(\mathrm{cell}_i) = 1$ if $\mathrm{cell}_i \in B$ clusters convex hull and 0 otherwise.

### 3 Supplementary Figures

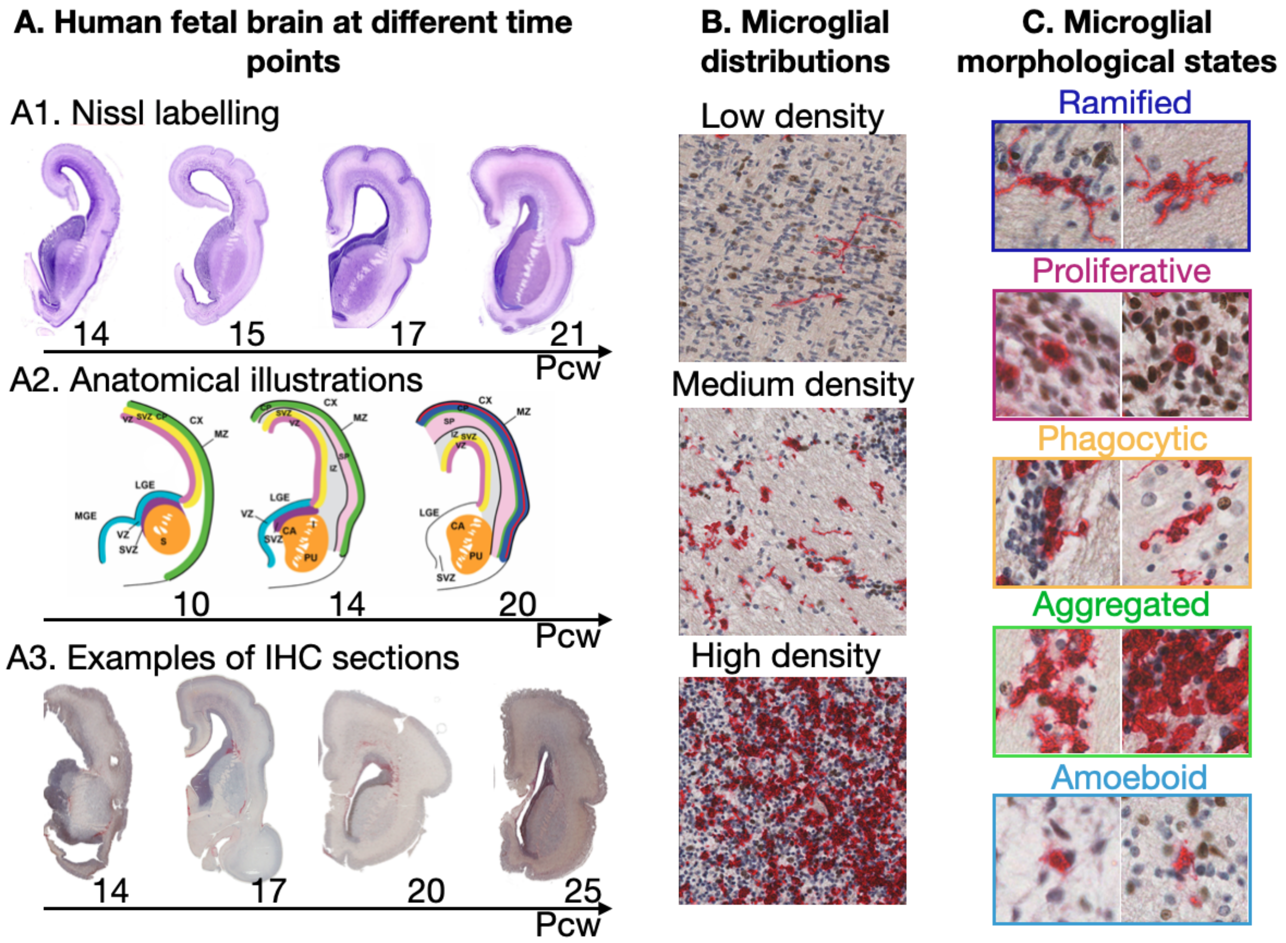

Figure S1: **Diversity of microglial morphological morphologies in human fetal tissue** **(A)** Anatomical mapping of the ganglionic eminence and striatum in human fetuses. A1. Nissl labeling in the coronal plane in fetuses aged between 14-21 pcw. A2. Anatomical sketches of fetal compartments in the same coronal plane between 10-20 pcw. A3. Example photomicrographs of microglia-labelled coronal slides using immunohistochemistry in the same plane in fetuses between 14-25 pcw. **(B)** Magnification of several regions arranged by microglia concentration. **(C)** Five different microglial morphological states: Ramified (dark blue), Proliferative (pink), Phagocytic (yellow),, Aggregated (green) and Amoeboid (light blue).

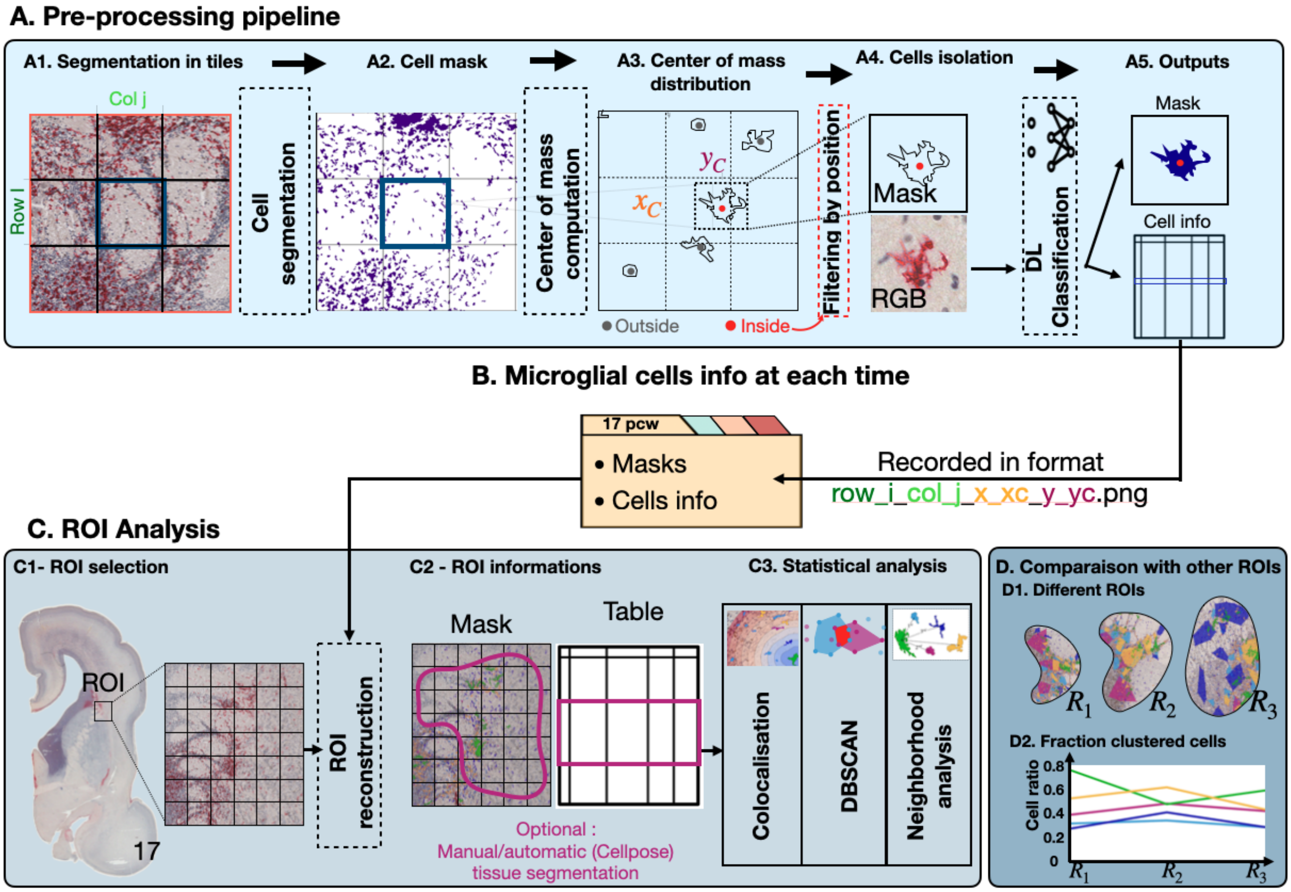

Figure S2: **Slide pre-processing and ROI reconstruction.** **(A)** Segmentation over the entire image based on slide tiling (size 1024x1024), where the slides are considered one after the other. **(A1)** Tile with 8 neighbors. **(A2)** Pixels belonging to microglial cells equal 1 and background 0. **(A3)** Center of mass $(x_c, y_c)$ computed for each cell. $x_c$ is the mean of pixels equal to 1 on x-axis (in the coordinate system of the 9-tile region). Same procedure for $y_c$. **(A4)** Cells whose center of mass are outside tile $(i, j)$ are excluded. For all the remaining cells inside tile $(i, j)$, RGB image of size 256x256 centered in $(x_c, y_c)$ is given as input to the DL classification. **(A5)** Saved features in a table ($x_c$, $y_c$, row-tile, col-tile, cell size, max length, probabilities of the cell to belong to all states, DL decision.**(B)** Cells mask saved in a database at each time. **(C)** Roi reconstruction and analysis. **(C1)** Selection of a ROI. **(C2)** Reconstruction of ROI with saved parameters. It is possible to filter the tissue manually or with pre-computed anatomical region mask. **(C3)** Statistics on the ROI. **(D)** Spatio-temporal statistics.

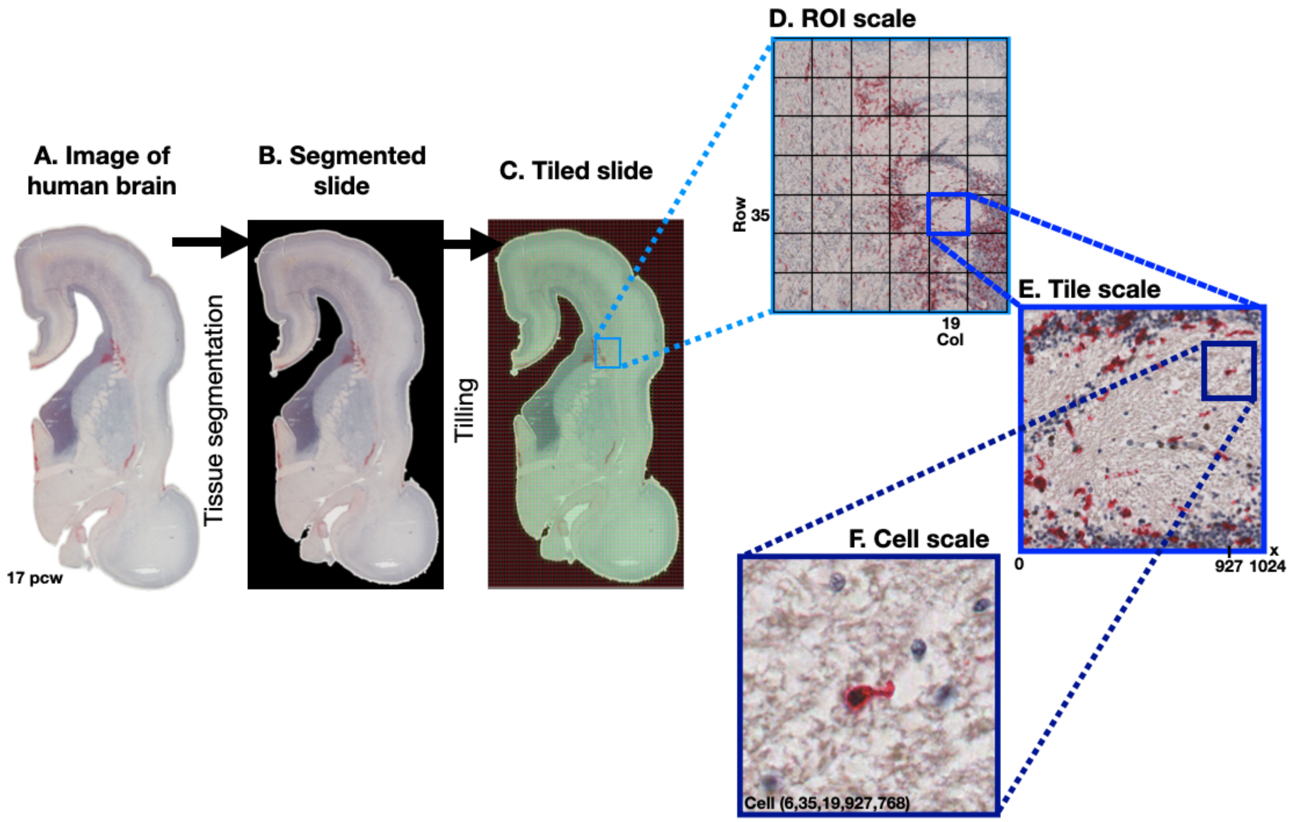

Figure S3: **Image manipulation at different scales.** **(A)** Example of an image of human fetal brain tissue at 17 post conceptional week (pcw) **(B)** Tissue extraction using otsu thresholding and morphological operations (see supplementary). **(C)** Division of the image into tiles of size 1024x1024 (tiling). Each tile is referenced by its row and column number. **(D)** Magnification of a region of interest containing several tiles. A border of size 1024 is considered around a ROI to take border effects into account. **(E)** Tile scale for which several cells are present. **(F)** Cellular scale where a single cell is located at the center of a crop of size 256x256. The cell is characterised by the tuple (slide number, tile row, tile col, centre of mass x in tile, centre of mass y in tile).

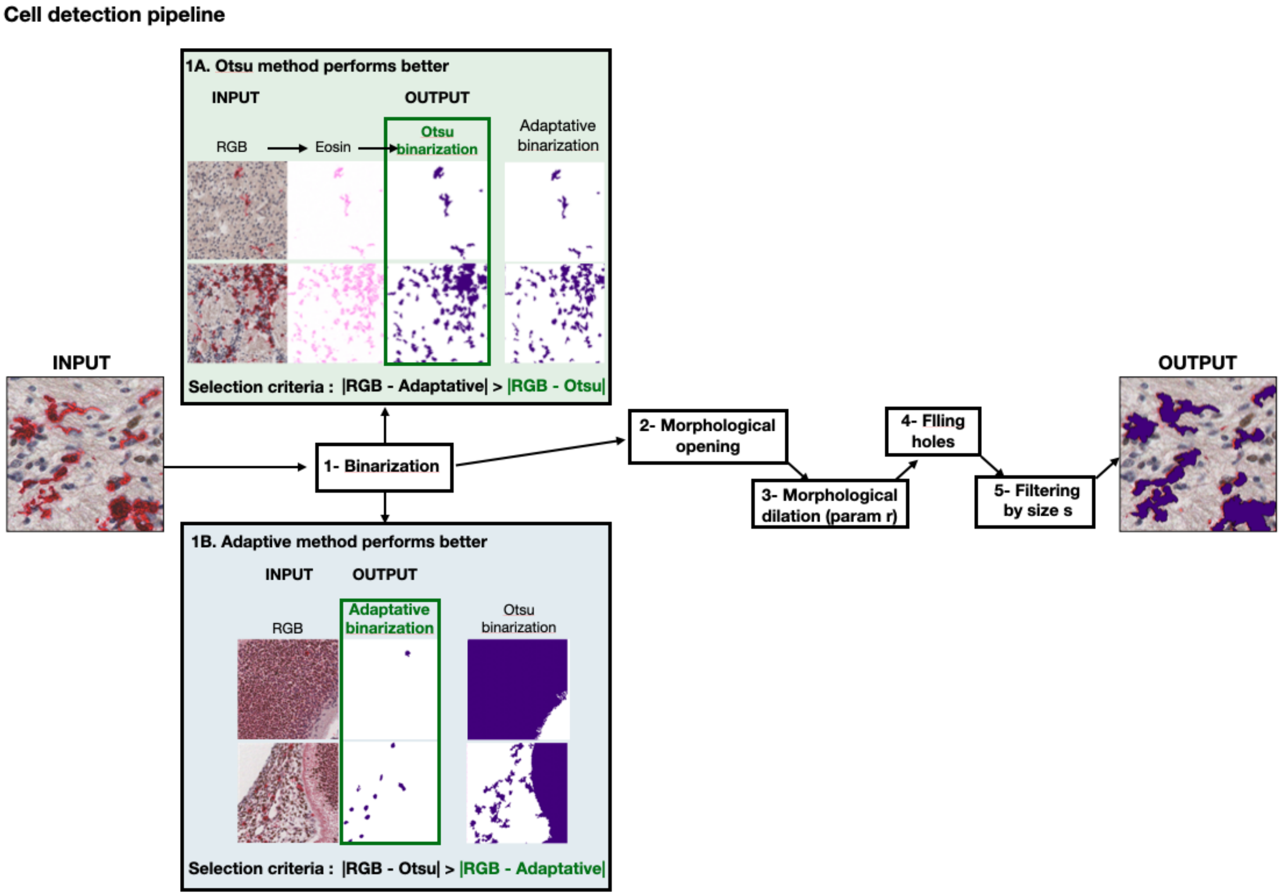

Figure S4: **Cells segmentation pipeline. (A)** Various image processing steps employed for cell detection and segmentation. **(B)** The input image (left) displays cells, which are segmented using a series of previously described processing steps (middle). This results in a binary image (right), where pixels corresponding to cell bodies are assigned a value of 1, while those belonging to the background are assigned a value of 0. DeepCellMap initiates the process by testing predefined sequences on different image sections. Users can then choose a starting sequence and iteratively refine it to determine the optimal sequence for all parts of the image.

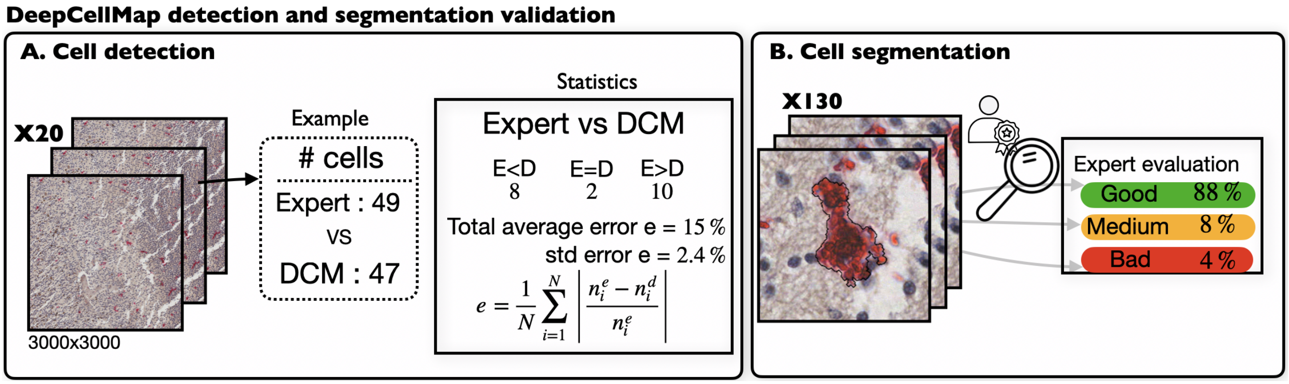

Figure S5: **DeepCellMap detection and segmentation validation.** **(A)** Dataset consisting of 20 randomly chosen images (left). Cells were identified both by DeepCellMap and an expert (middle), where the expert (resp.DCM) found 49 (resp. 47). Results on the datas where the expert found less cells than DCM in 8/20 images, the same amount in 2 and more in 10/20 (Right). The overall result differs from 15%. **(B)** Cell detection comparison between expert and DCM: expert evaluate the segmentation quality in three categories (Good, Medidum and Bad) on 130 images.

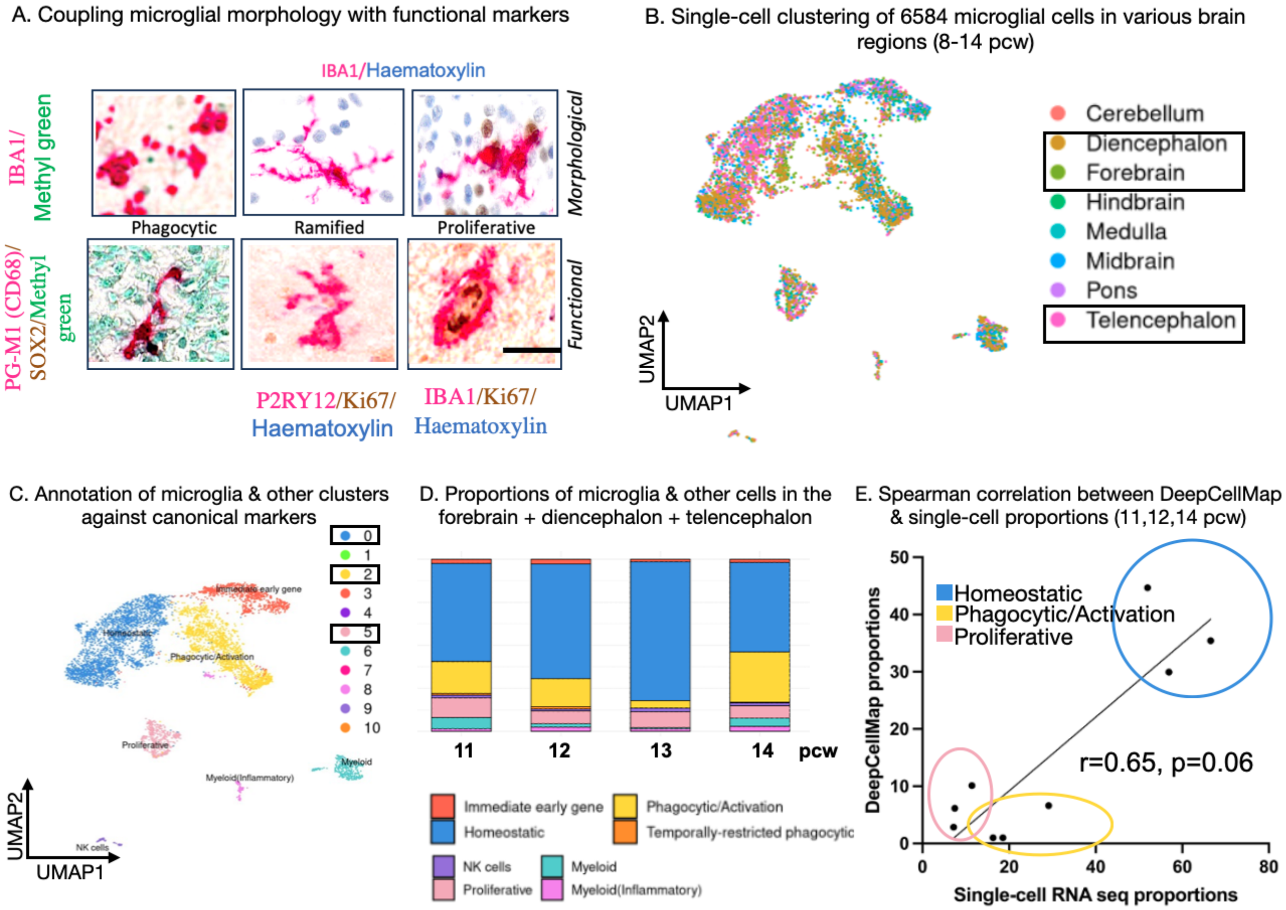

Figure S6: **Microglial morphology coupled with function in human fetal tissues using histological & sc-RNAseq analyses (A)** Representative examples of the coupling between morphology and function in developing human tissues using P2RY12 as a homeostatic microglial marker, PG-M1 as a lysosomal marker and Ki67 as a proliferative marker. **(B)** Single-cell clustering across sampled regions of the developing brain between 10-15 pcw (Braun et al., 2023). **(C)** Annotation of microglia and other clusters in the developing brain based on the gene list in (supplementary table) between 10-15 pcw. **(D)** Bar plot showing in the forebrain+diencephalon+telencephalon regions the combined the proportions of microglia and other cell types between 10 and 15 pcw. **(E)** Spearman's correlation between the proportions of microglia calculated by DeepCellMap in 3 classes (ramified/homeostatic, phagocytic and proliferative) in the forebrain and proportions of microglia calculated from single-cell RNAseq data (homeostatic, phagocytic and proliferative) matched for 3 postconceptional weeks 11,12 and 14 (r=0.64, p=0.06). If we take proliferative and ramified, r = 0.85. This shows a good positive correlation between DeepCellMap and single-cell-transcriptomic proportions in two independent datasets.

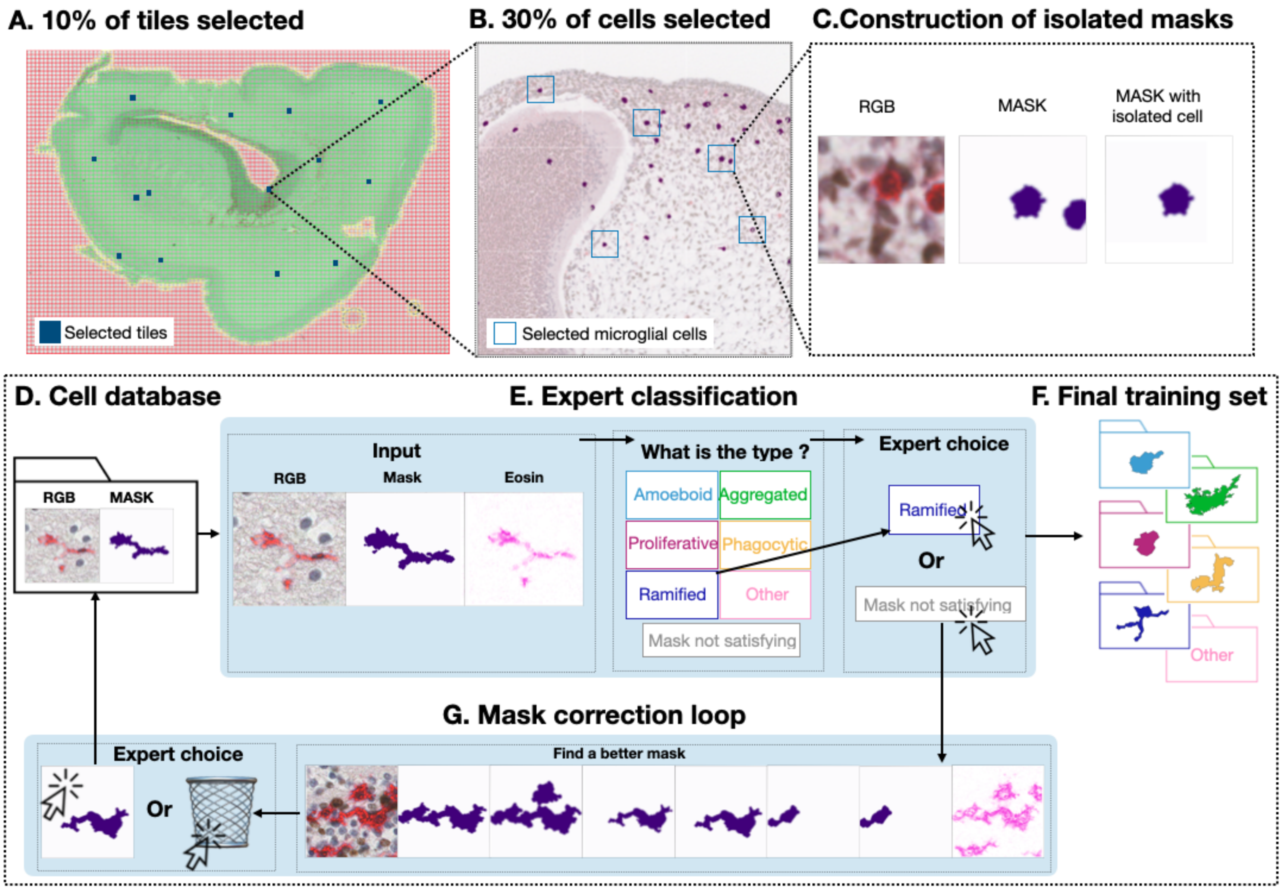

Figure S7: **Training set constitution pipeline.** Cells from the training set were randomly picked from the data using the following procedure.**(A)** $p_1$% tiles were selected (blue boxes) in each image.**(B)** Magnification of a tile where microglial cells have been segmented (purple) and from which $p_2$% are then kept. **(C)** Isolation of the mask corresponding to the cell of interest. **(D-G)** Label attribution to each cell. **(D)** Database of unlabelled microglial cells. **(E)** Manual annotations by the expert. **(F)** Finalized training set. **(G)** Decision to change mask when an incorrect one is used. If no better masks are found, this example is eliminated.

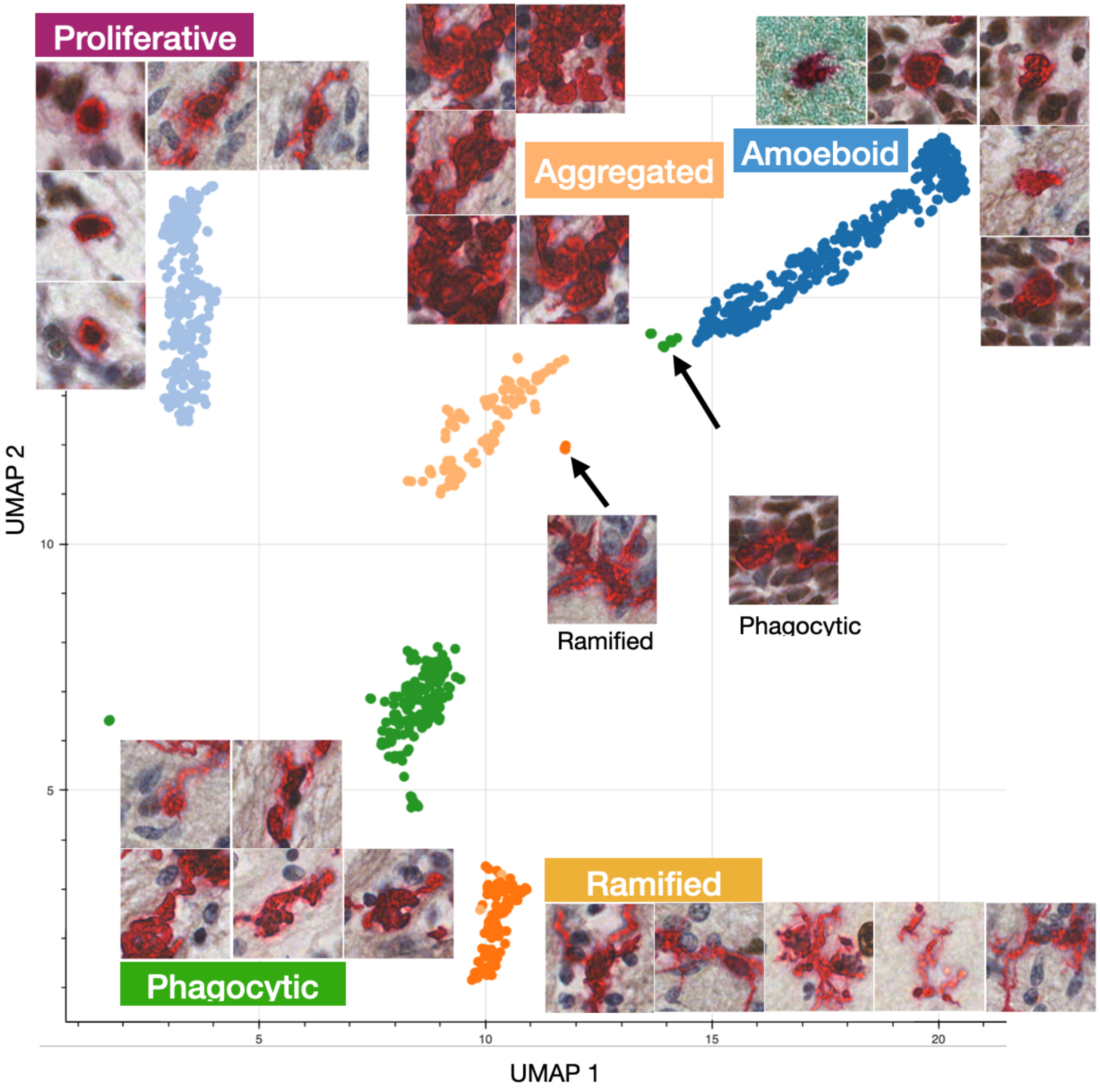

Figure S8: **Training set heterogeneity.** Distribution of the training base examples in the plane of the first two-UMAP components (dimension reduction algorithm). Each point corresponds to an image of the training set. Proliferative (pink), Amoeboid (light blue), Phagocytic (yellow), Ramified (dark blue) or Aggregated (green).

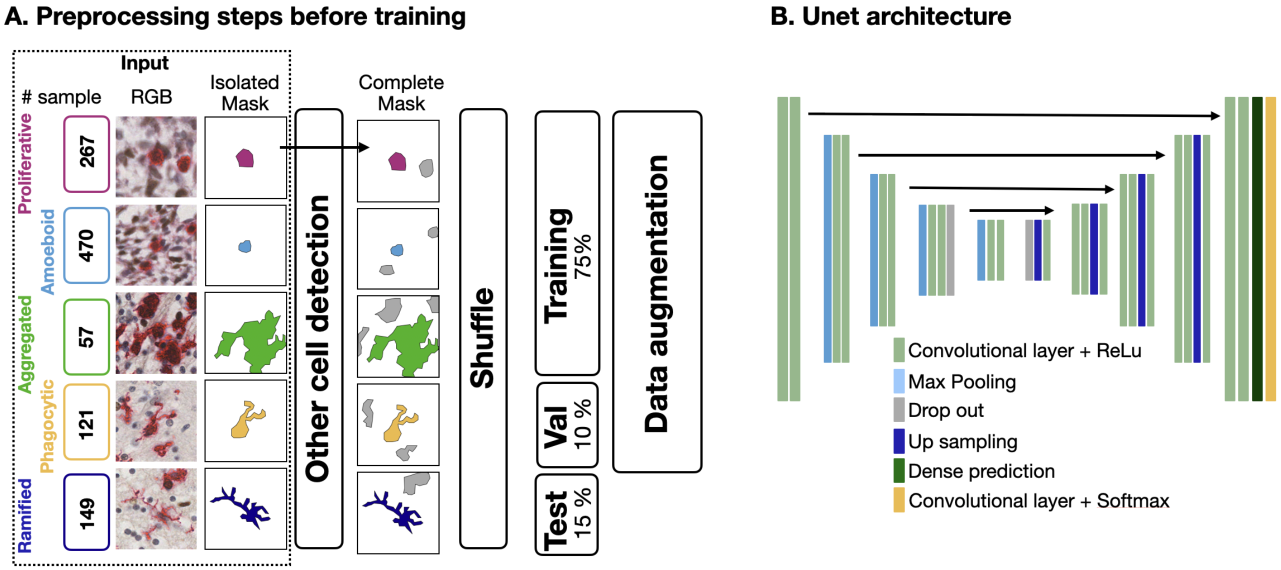

C. Hyperparameters selection

| Pooling steps | batch size | global_f1 |
|---|---|---|
| 3 | 2 | 0.69 |
| 3 | 3 | 0.64 |
| 3 | 4 | 0.66 |
| 4 | 2 | 0.73 |
| **4** | **3** | **0.81** |
| 4 | 4 | 0.76 |

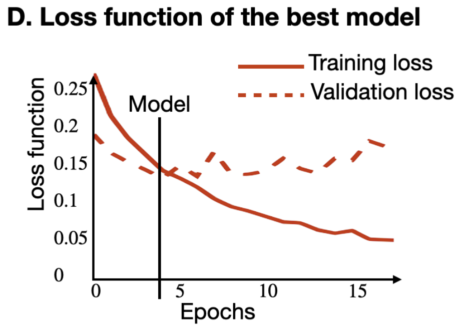

Figure S9: **Deep-learning classification details.** **(A)** Pre-processing steps before training. Input consists of several paires of RGB images and masks for each of the five microglial morphological states.For each mask, we added the mask of the neighboring detected cells with label "Other". We then shuffle the datasets and split it into Training (75%), Test (15%) and Validation set (10%). **(B)** Architecture of the U-Net based DL-network. **(C)** Hyper-parameters table showing the best choice of the parameters based on f1-score. **(D)** Training and validation loss of the best model. The final model is the one that minimises the validation loss.

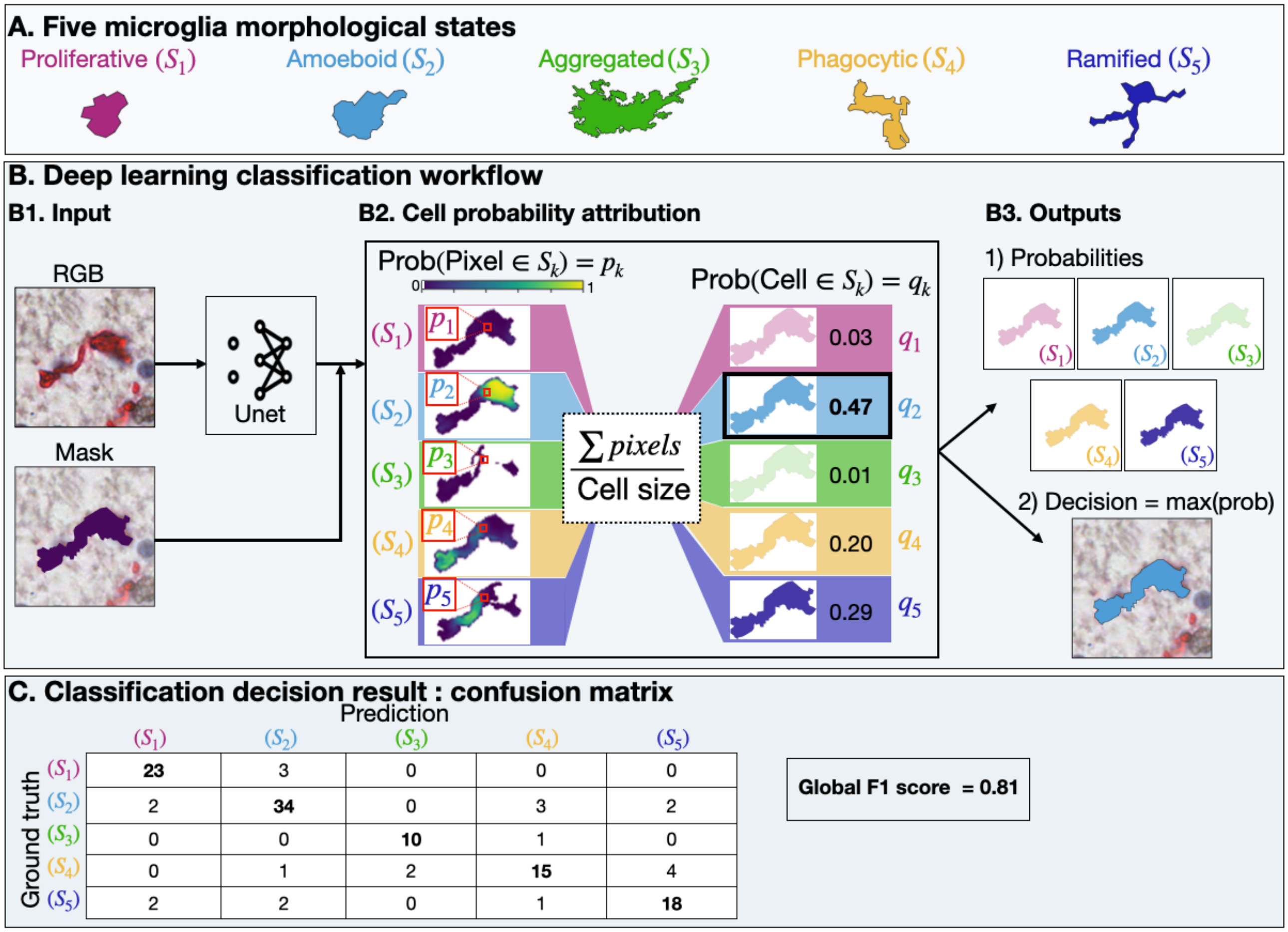

Figure S10: **Deep learning classification pipeline.** **(A)** Microglial cells are classified into 5 morphological morphologies: proliferative (pink), amoeboid (light blue), aggregated (green), phagocytic (yellow) and ramified (dark blue). This does not preclude the existence of other morphological states but our focus here was on these 5 that could be reliably identified using DeepCellMap. **(B)** Deep-learning workflow starting with an RGB image of a cell. **(B1)** Input to the deep learning model U-Net. Another processing step is to filter the image by the mask of a cell. **(B2)** Left: Each pixel has a probability $p_k$ to belong to a state $S_k$ for $k = 1..5$. Right: a post-processing step attributes a global state probability $q_k$ by averaging the probability of each pixel belonging to a given cell (right). **(B3)** The output consists of the probabilities $q_1, ..q_5$) for a microglial cell to belong to a given class and the decision of the network (maximum value of the probability $q_k$). **(C)** Confusion matrix resulting from the U-Net decision algorithm (max probability), comparing ground truths (rows) with predictions (columns) of the network applied to the test ensemble (Global f1-score is 0.81).

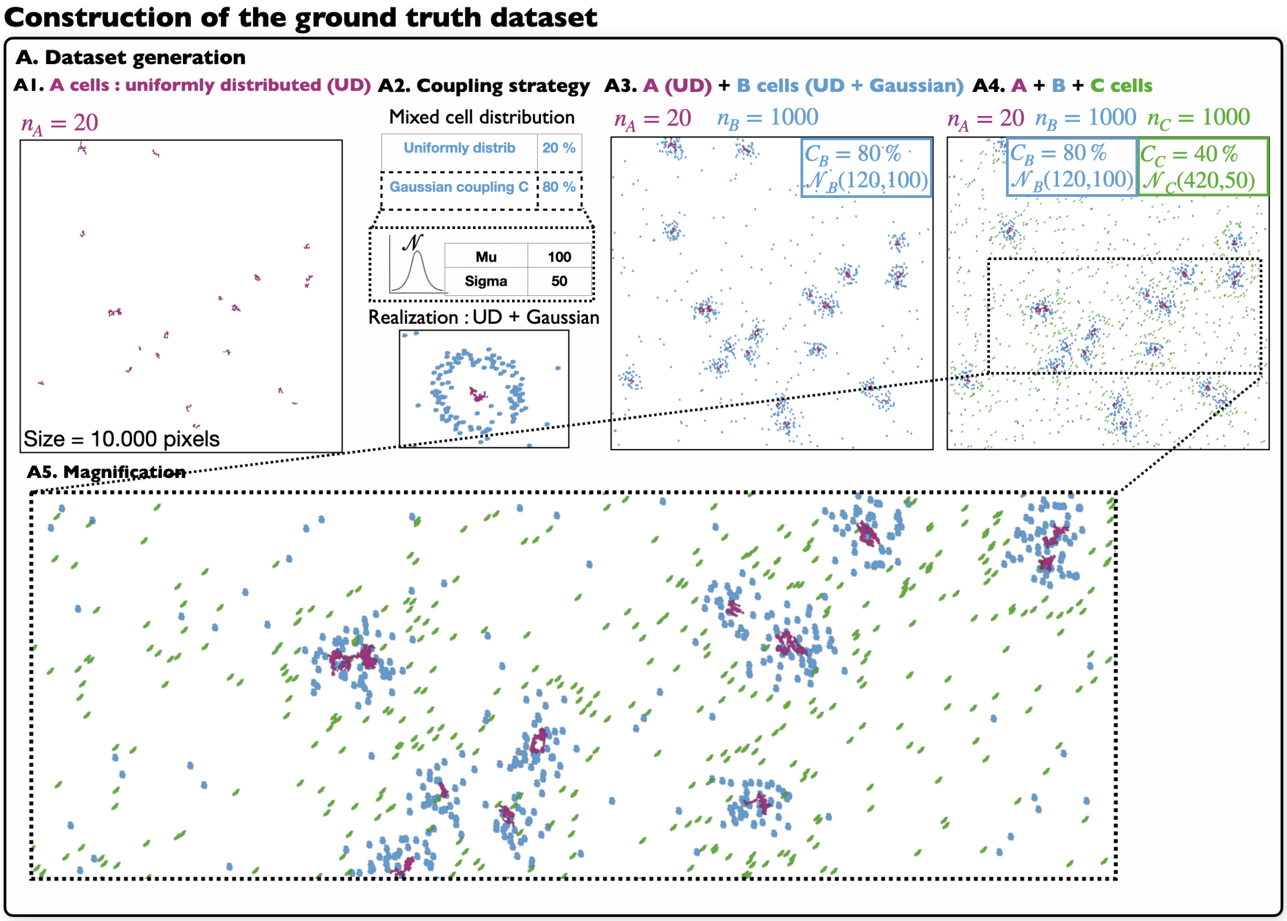

Figure S11: **Generation of a ground truth distribution.** **(A)** Dataset generation. **(A1)** A cells are uniformly distributed (UD). **(A2)** Coupling parameters of the mixed cell distribution with a fraction of cells UD and the others coupled to A cells by a Gaussian distribution with mean distance $\mu$ and variance $\sigma$, a realisation is shown (bottom). **(A3)** A cells (UD) and B cells (blue) distributed in the image with respect to the mixed parameters (UD + Gaussian). **(A4)** A cells (UD), B cells (UD + Gaussian) and C cells (UD + Gaussian) are distributed in the image. **(A5)** Magnification.

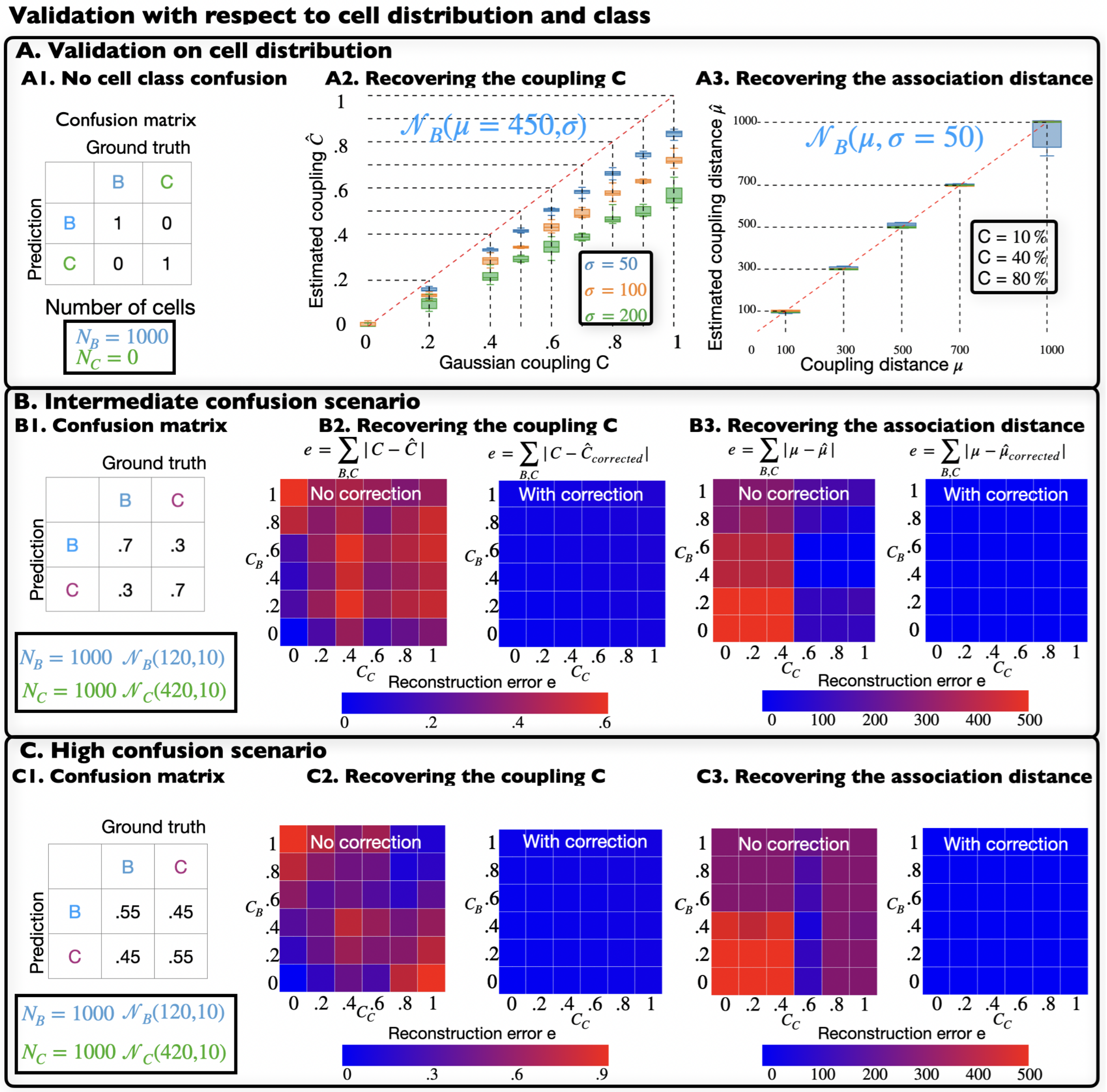

Figure S12: **Level set analysis on synthetic ground truth.** **(A)** Validation of DeepCellMap levelset analysis when there is no classification errors (scenario 1). **(A1)** Confusion matrix and number of cells.**(A2)** Recovering the coupling C for different value of $\sigma$. **(A3)** Recovering the association distance $\mu$ for different values of the coupling C. **(B)** Validation of DeepCellMap for scenario 2 (intermediate confusion). **(B1)** Confusion matrix. **(B2)** Coupling recovery errors (heatmap) for several coupling configurations of $C_B$ and $C_C$ before (left) and after (right) correcting the levelset estimators (see method). The total error account for B and C coupling reconstruction. **(B3)** Association distance recovery errors (heatmap) for several coupling configurations of $C_B$ and $C_C$ before (left) and after (right) correcting the level set estimators (see method). The total error account for B and C coupling reconstruction. **(C)** Validation of DeepCellMap for scenario 3 (high confusion). **(C1)** Classification confusion matrix. **(C2)** Coupling recovery errors (heatmap) for several coupling configurations of $C_B$ and $C_C$ before (left) and after (right) correcting the level set estimators (see method). The total error account for B and C coupling reconstruction. **(C3)** Association distance recovery errors (heatmap) for several coupling configurations of $C_B$ and $C_C$ before (left) and after (right) correcting the level set estimators (see Methods). The total error account for B and C coupling reconstruction.

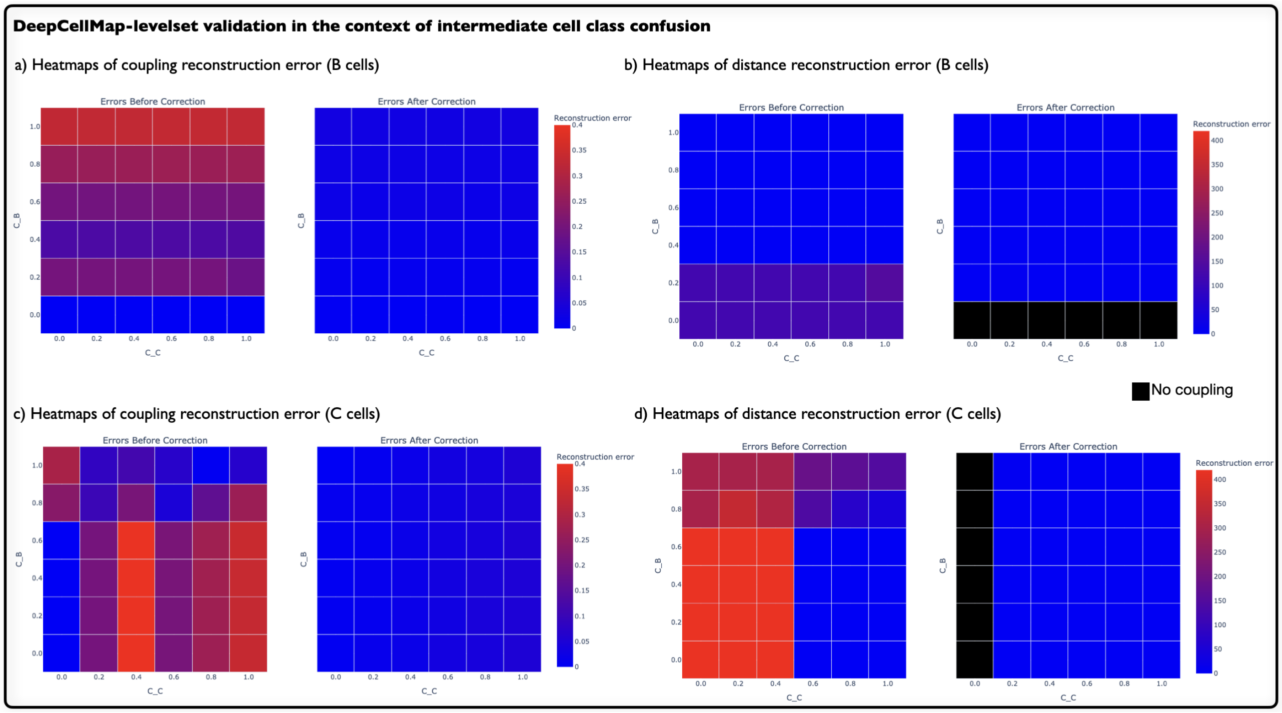

Figure S13: **DeepCellMap levelset analysis validation in the context of intermediate cell class confusion (scenario 2) .** **(a)** Coupling reconstruction errors (Heatmaps) vs coupling constant $C_B$ and $C_C$ before and after correction. The error accounts for B cells only . **(b)** coupling distance reconstruction errors (Heatmaps) vs coupling constants $C_B$ and $C_C$ before and after correction. **(c)** Coupling reconstruction errors (Heatmaps) for several configurations of $C_B$ and $C_C$ before and after correction. **(d)** Coupling distance reconstruction errors (Heatmaps) vs coupling constants$C_B$ and $C_C$ before and after correction.

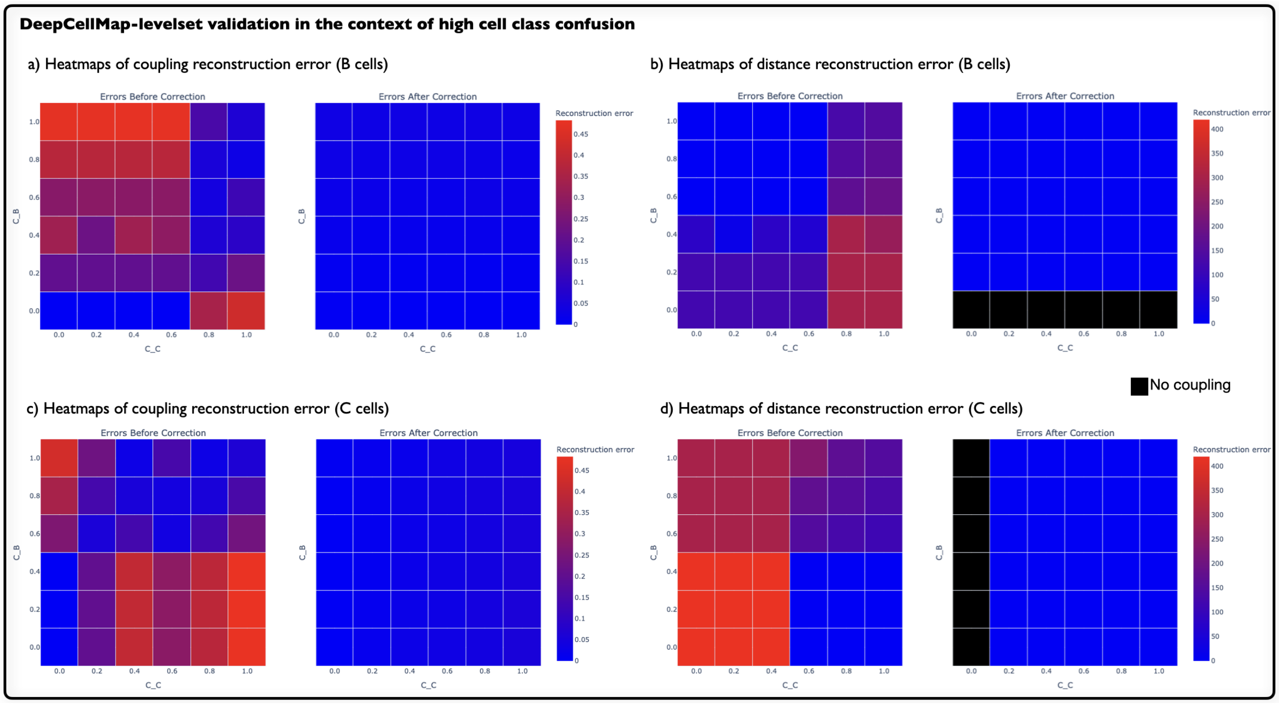

Figure S14: **DeepCellMap validation for the levelset analysis in the context of high cell class confusion (scenario 3).** See legend of Fig. ??.

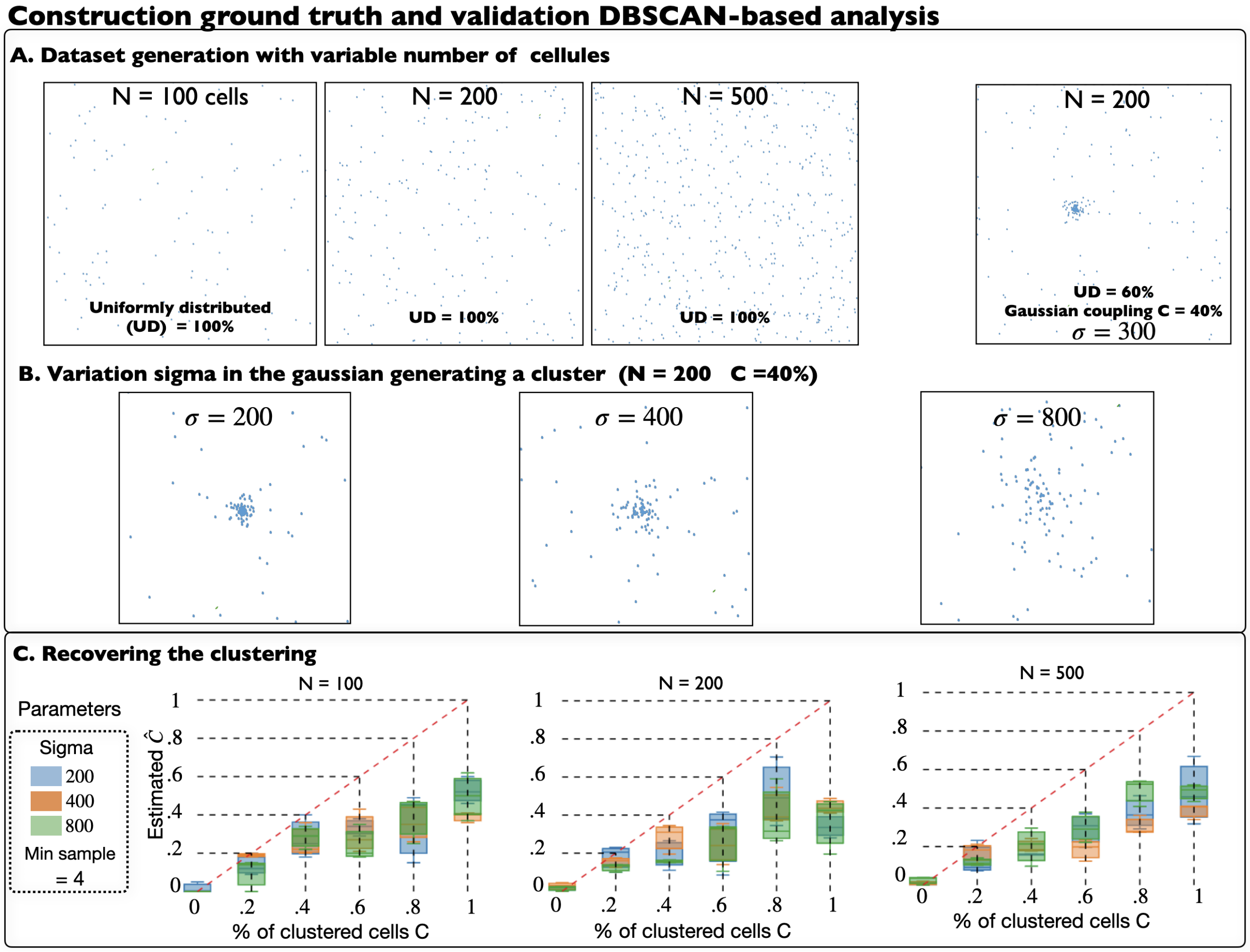

Figure S15: **Validating the optimized DBSCAN clustering analysis with synthetic simulations.** **(A)** Generating synthetic ground truth for with uniformly distributed (UD) cells ($N = 100, 200$ or $500$), or partially clustered with $C = 40\%$ of the $N = 200$ cells that are distributed in Gaussian clusters ($\sigma = 300$). **(B)** Generation of 200 cells with 40% clustered for various values of $\sigma = 200, 400$and 800. **(C)** Recovering the clustering percentage $C$ for different value of $\sigma$.

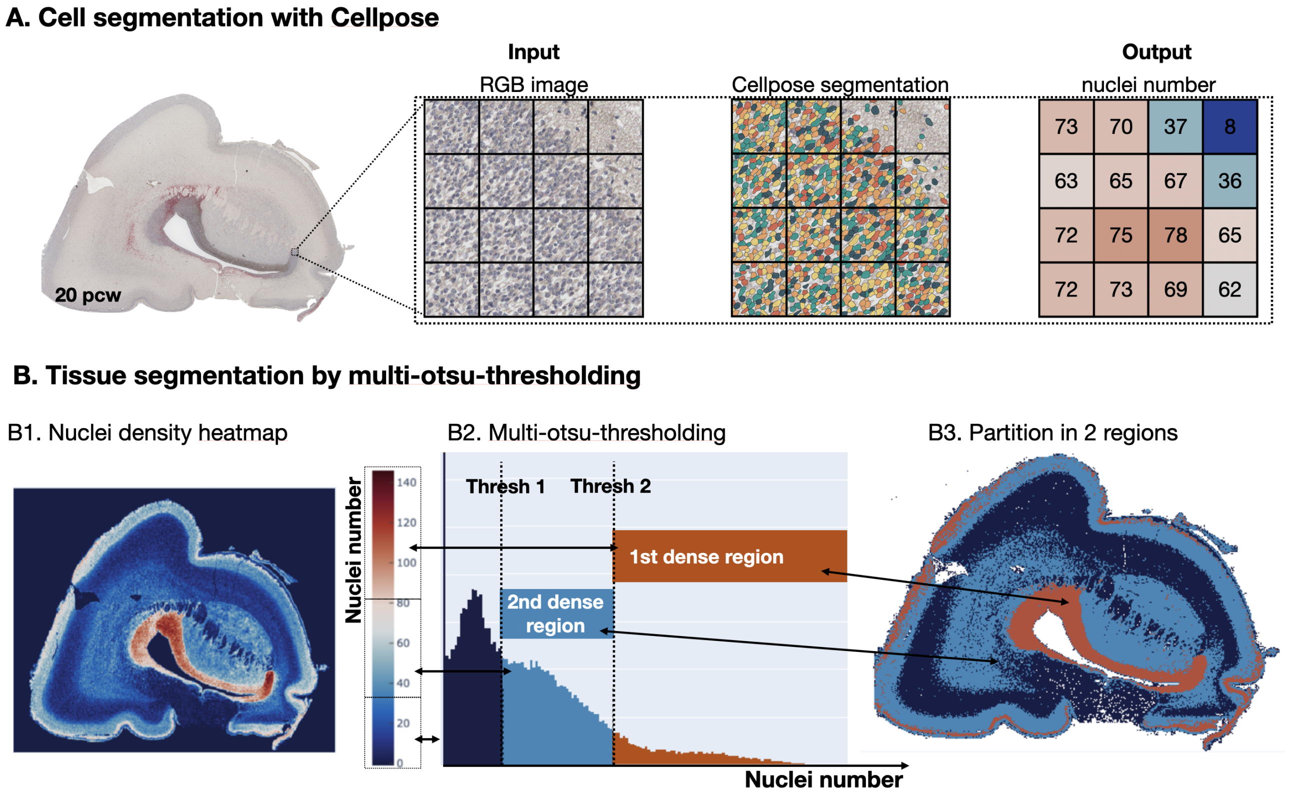

Figure S16: **Fetal tissue segmented by CellPose algorithm.** **(A)** **(Left)**- Magnification of a tile of size (1024x1024) decomposed in 16 crops of size (256x256). **(Middle)**- Input: cell organization, followed by CellPose segmentation. Right: Number of segmented nuclei for each crop. **(B)** CellPose segmentation on the entire tissue based on density. **(B1)** cellPose applied to the entire tissue. **(B2)** Multi-Otsu thresholding method to decompose the tissue into 2 regions (color bar), with two thresholds Thresh1 and Thresh2. **(B3)** Output : decomposition of the tissue in 2 regions : high density (red) and medium density (light blue).

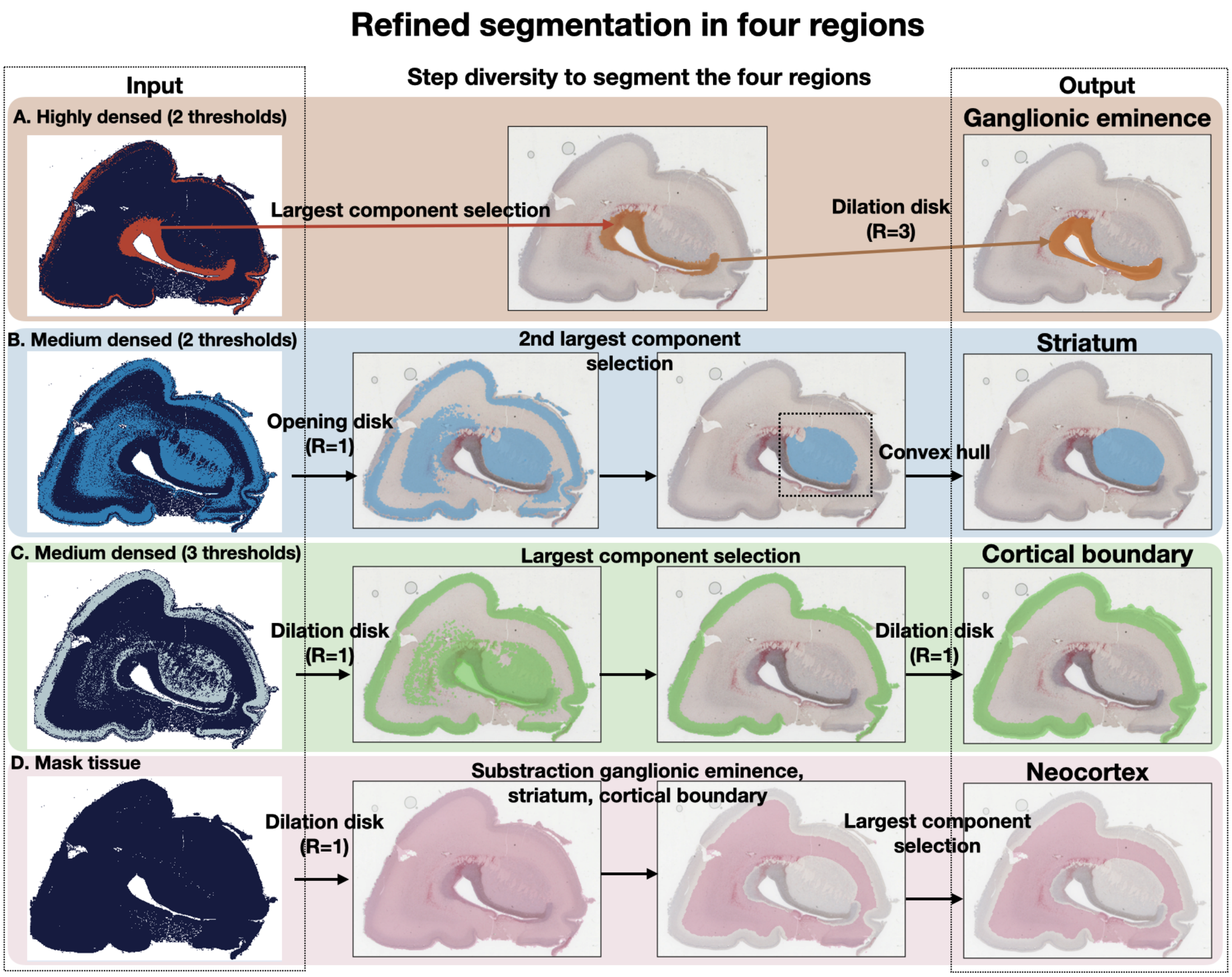

Figure S17: **Refined segmentation of the four regions from nuclei density.** **(A)** Left: input of the tissue segmented by multi-otsu thresholding with 2 thresholds (see Fig. **??**). Middle: Automated selection of the largest component, followed by morphological dilation to increase the mask. Right: selection of the ganglionic eminence. **(B)** Left: Multi-otsu thresholding with two thresholds. Middle: Morphological dilation (Radius=1) followed by the selection of second largest component (based on the surface computation) with a final convex hull estimation Right: the output is the striatum region. **(C)** Left: Multi-otsu thresholding with three thresholds. Middle: Morphological dilation (Radius=1) followed by the selection of the largest component (based on the surface comparison) with a final dilation (Radius=1). Right: the output is the cortical boundary. **(D)** Left: input mask of the tissue. Middle: substraction of the selected region in A-B-C, followed by the selection of the largest component (based on the surface comparison). Right: the output is the Neocortex.

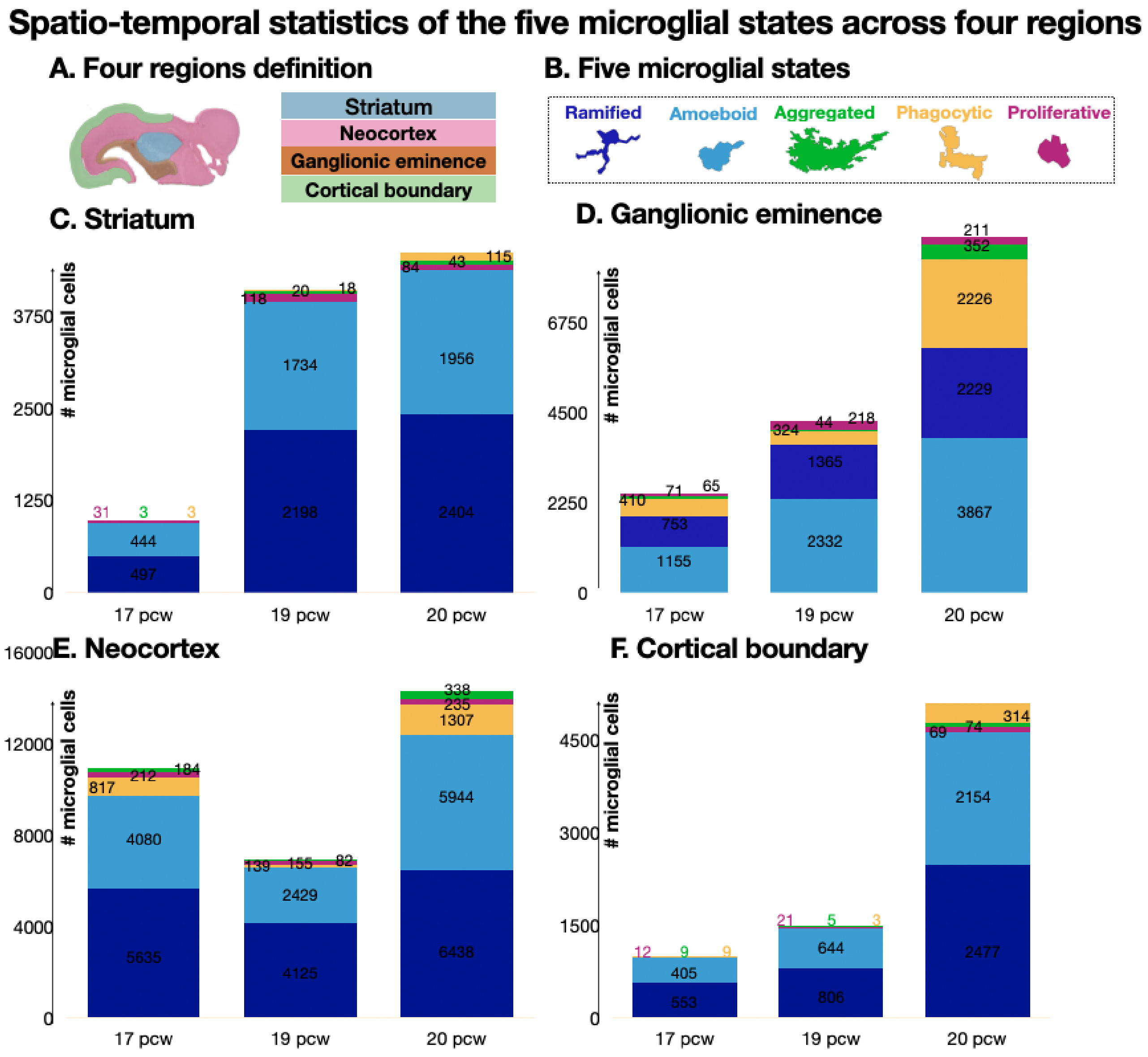

Figure S18: **Spatio-temporal statistics of the five microglial morphologies across four regions.** The numbers are estimated from the deep learning algorithm for three pcw (17, 19 and 20). **(A)** Definition of the 4 regions : striatum (blue), neocortex (pink), ganglionic eminence (orange), cortical boundary (green). **(B)** Definition of the five microglial states : Proliferative (pink), Amoeboid (light blue), Aggregated (green), Phagocytic (yellow), Ramified (dark blue). **(C)** Distribution of microglial states in the striatum, **(D)** in the ganglionic eminence, **(E)** neocortex and **(F)** cortical boundary.

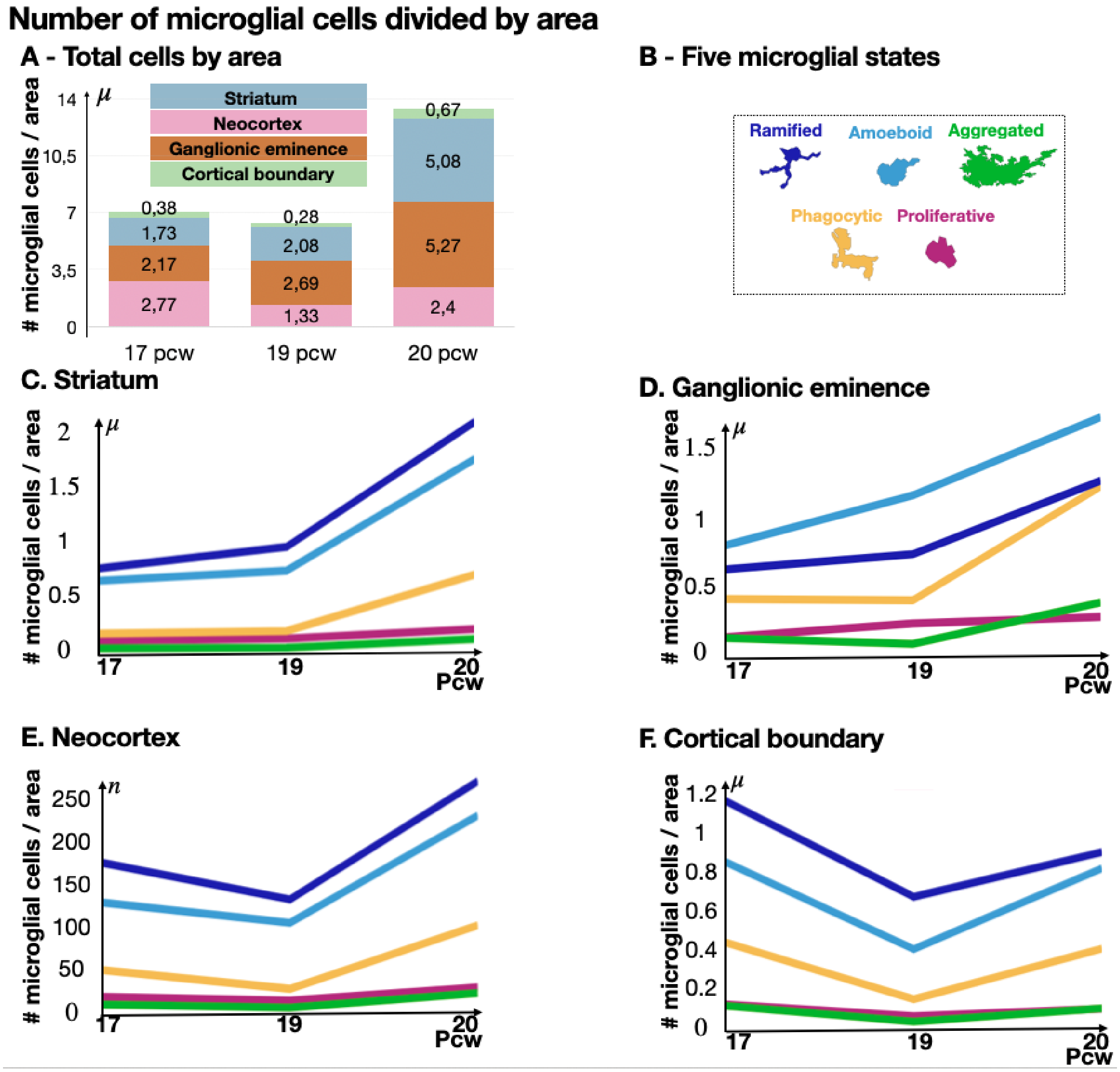

Figure S19: **Spatiotemporal concentration (number divided by area) of the five microglial states across four regions.** The numbers are estimated from the deep learning algorithm for three pcw (17, 19 and 20). **(A)** Changes across time of the concentration (total number of cells divided by area) in the four regions. **(B)** Five microglial states. Concentration changes of the five microglial states in the striatum **(C)**, the ganglionic eminence **(D)** the ganglionic eminence, **(E)** the neocortex and cortical boundary **(F)**.

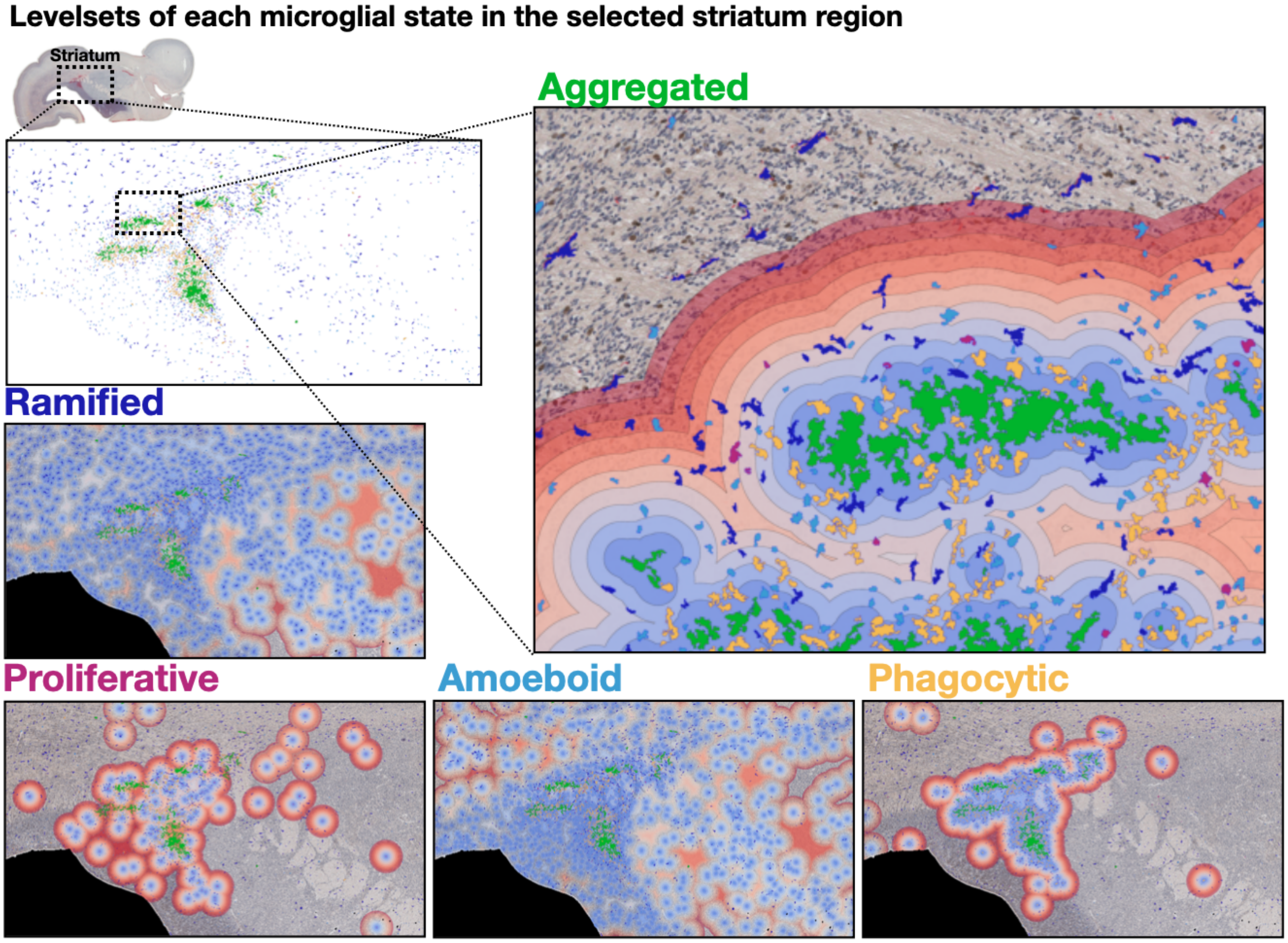

Figure S20: **Levelset analysis on the striatum.** Selection of the subsection of the Striarum. Levelsets computed for each of the 5 subtypes: Proliferative (pink), Amoeboid (light blue), Aggregated (green), Phagocytic (yellow), Ramified (dark blue).

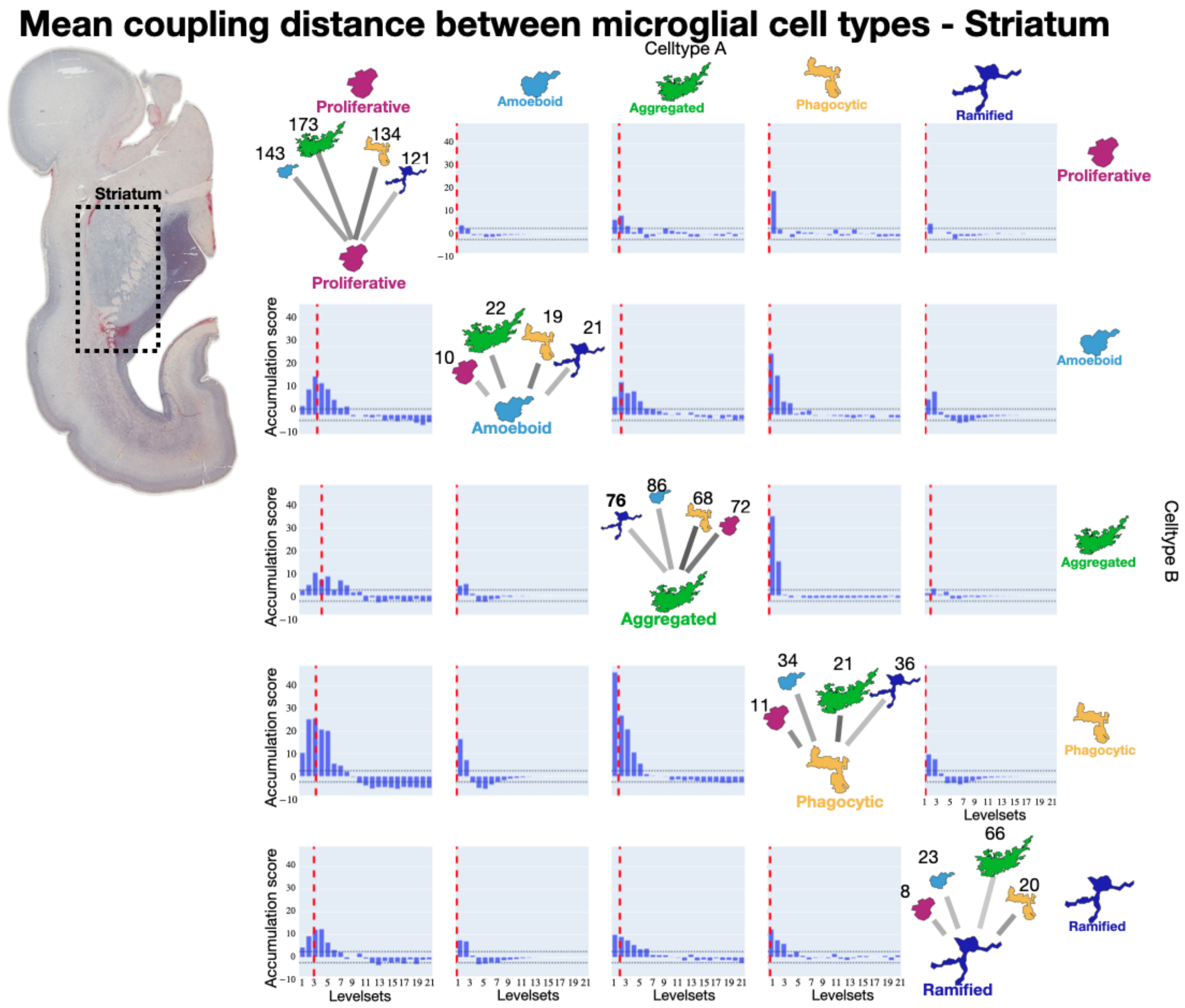

Figure S21: **Mean coupling distance matrix computed from the Striatum.** 5 microglia type accumulation matrix showing the threshold (dashed horizontal line) and the mean coupling distance (vertical line).

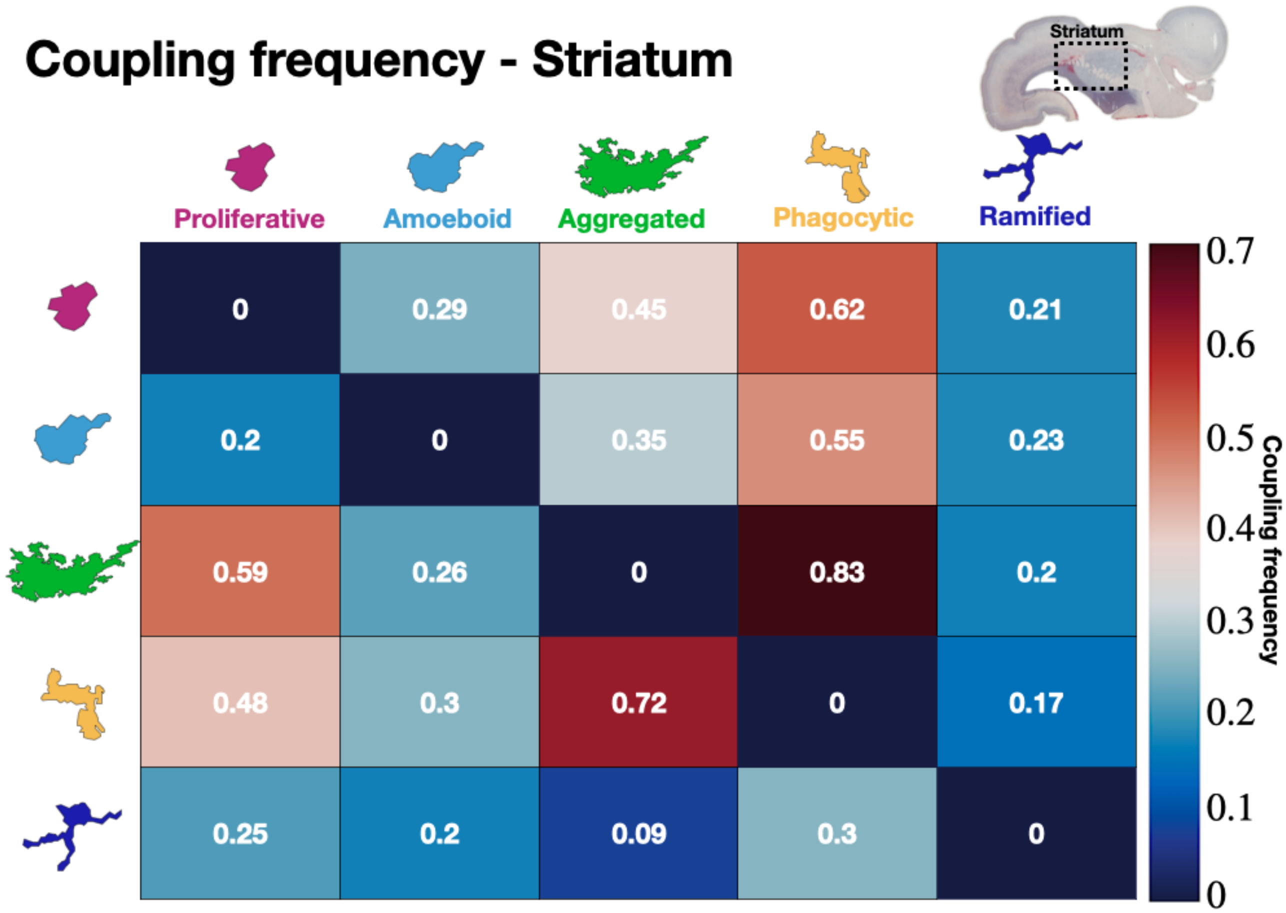

Figure S22: **Coupling frequency.** Matrix of association between the 5 microglial types in Striatum at 17 pcw. Aggregated and phagocytic share the highest coupling frequency, suggesting that there are specifically coupled.

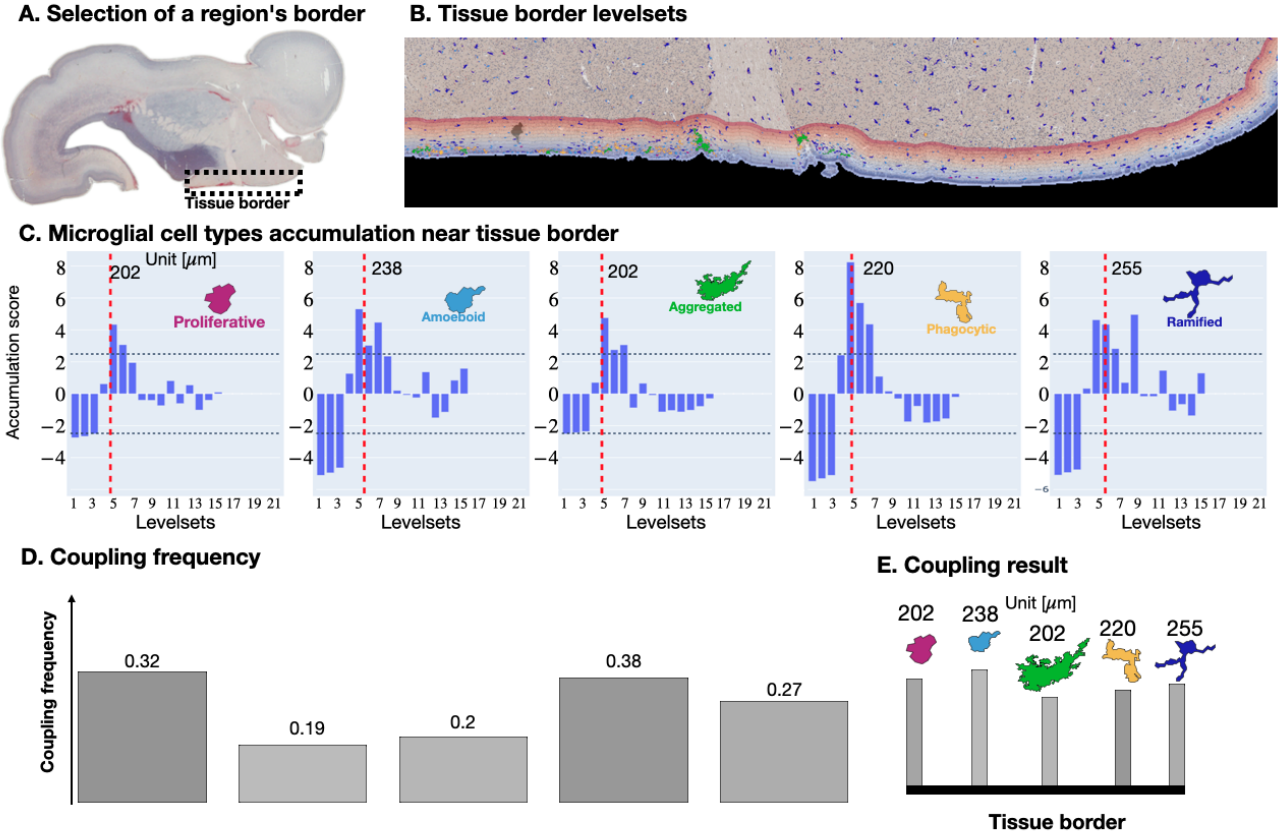

Figure S23: **Coupling with a region or tissue border.** **(A)** Selection of a region's border. **(B)** Microglial cells in the ROI and levelsets defined by tissue border. **(C)** Accumulation score of the five microglial states in the levelsets. **(D)** Coupling frequency of each cell with the border. **(E)** Coupling result shown in the form of trees, where the root (defining level sets) is the border of the tissue (in the example) or the border of a region. The leaves are the microglia distributed in the levelsets, quantified by the mean coupling distance in $\mu m$ (above). .

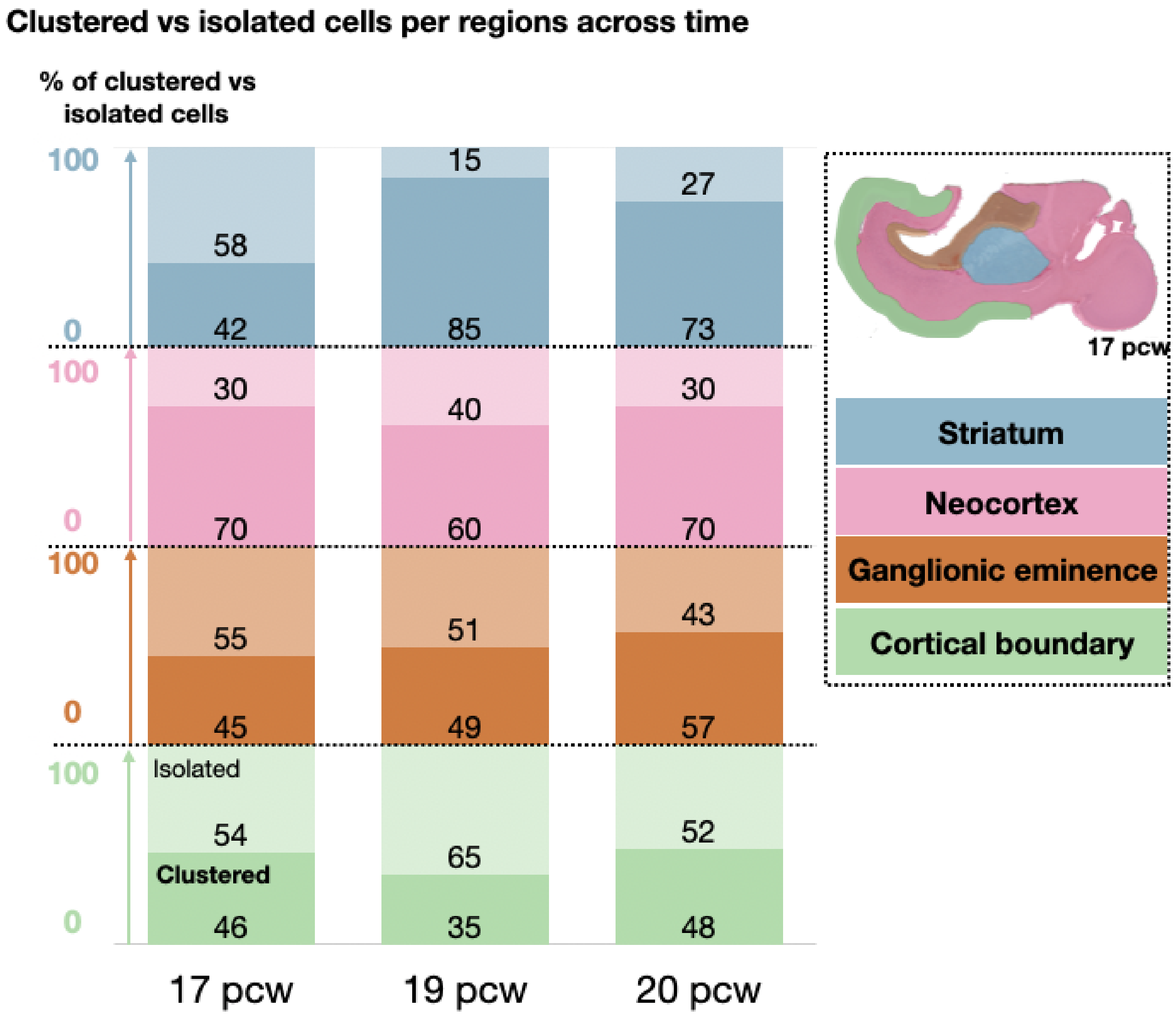

Figure S24: **Clustered vs Isolated cells per region across time.** The four regions are striatum (blue), neocortex (pink), ganglionic eminence (orange), and cortical boundary(green). The percentage of clustered (bold) and isolated (light) cells are shown for three post-conceptional weeks 17, 19 and 20.

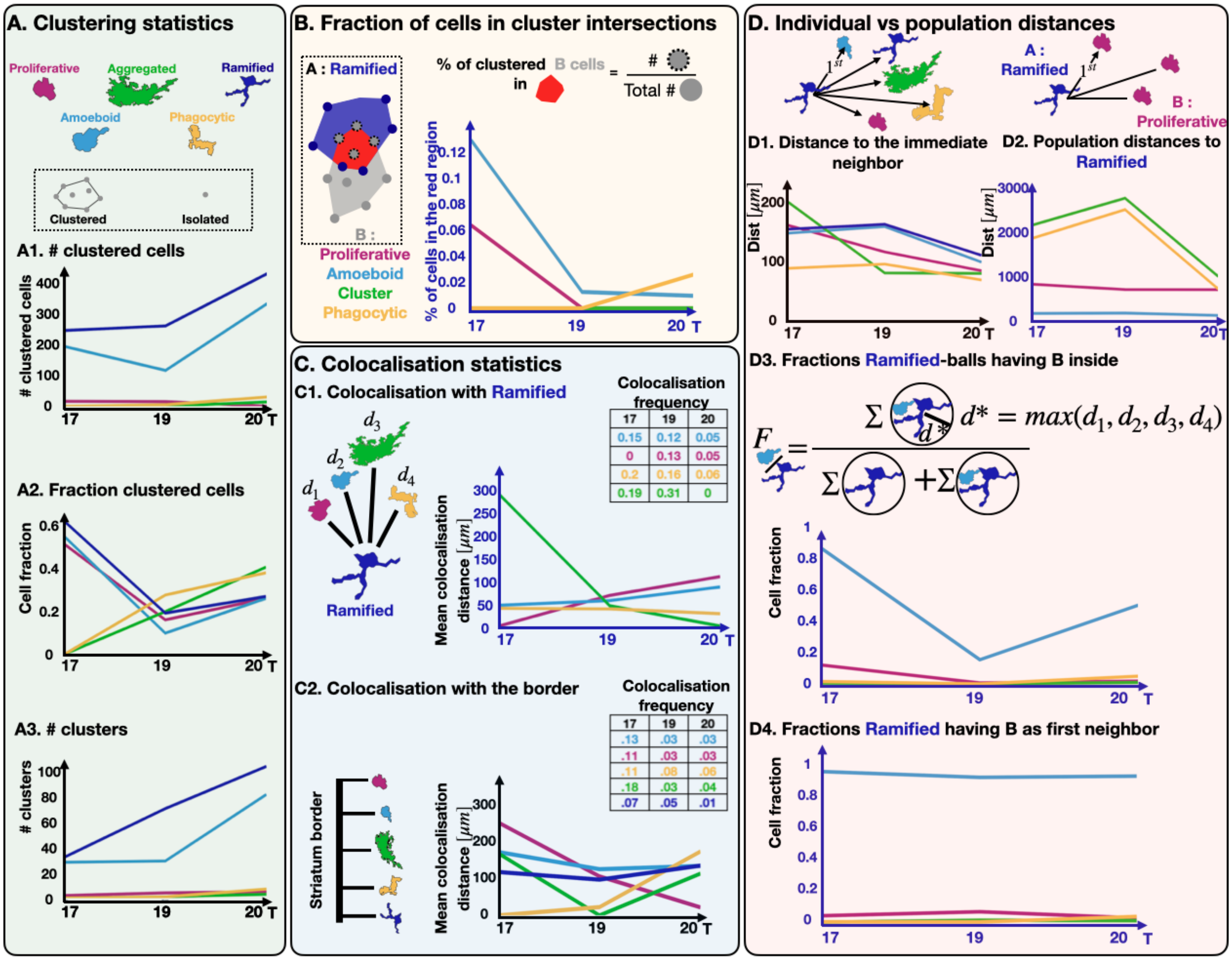

Figure S25: **Spatiotemporal statistics of microglia in the striatum.** **(A)** Clustering of the different microglia types: Proliferative (pink), Amoeboid (light blue), Aggregated (green), Phagocytic (yellow), Ramified (dark blue). Clusters are computed using DBSCAN (MinSample = 4). (A1) Time evolution for the number of cluster cells, (A2) fraction of clustered cells, (A3) number of clusters. **(B)** Mixing quantification, computed as the fraction of clustered B cells present in A clusters: Proliferative (pink), Amoeboid (blue), Aggregated (green), and Phagocytic vs Ramified (yellow). **(C)** (C1) Average coupling distance between Proliferative (pink), Amoeboid (light blue), Aggregated (green), Phagocytic (yellow) vs Ramified (blue axis). (C2) Average coupling distance between the edge of the ganglionic eminence and each population. Frequencies of coupling are summarized in tables. **(D)** Individual versus population distances. We chose Ramified as the reference. (D1) Mean distance ($\mu m$) to the first neighbor that can be one of the 5 microglial types. (D2) distance ($\mu m$) to the shortest of the second population, averaged over all ramified cell position. (D3) Fraction of balls of radius $d^*$, centered around a microglia type containing at least one cell type inside. The distance $d^*$ is chosen at each time as the maximum among the mean coupling distances (d1, d2, d3, d4). (D4) Fraction of cells (Ramified cells) having another cell type as first neighbor, independently of any fixed ball.

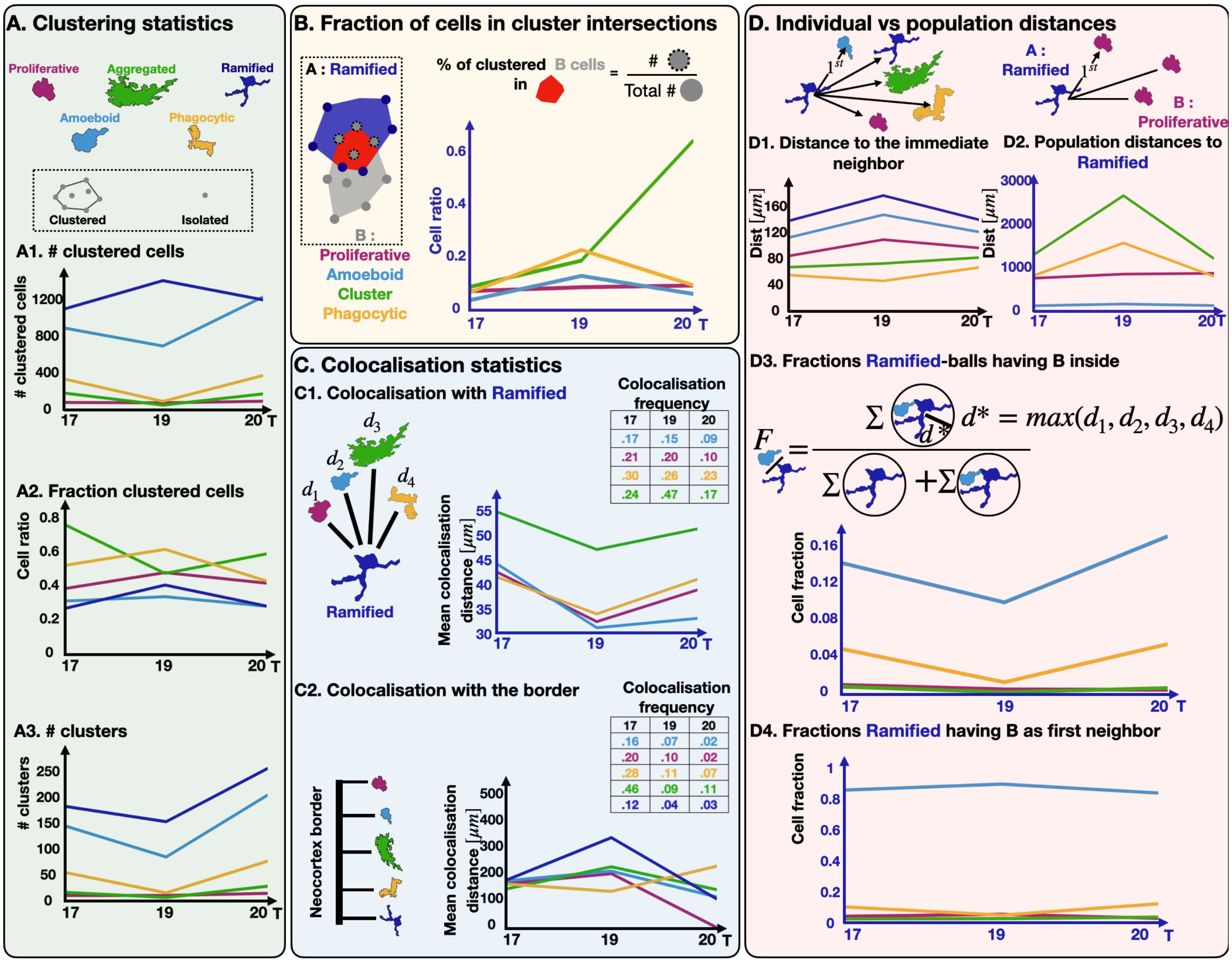

Figure S26: **Spatiotemporal statistics of microglia in the neocortex.** **(A)** Clustering of the different microglia types: Proliferative (pink), Amoeboid (light blue), Aggregated (green), Phagocytic (yellow), Ramified (dark blue). Clusters are computed using DBSCAN (MinSample = 4). (A1) Time evolution for the number of cluster cells, (A2) fraction of clustered cells, (A3) number of clusters. **(B)** Mixing quantification, computed as the fraction of clustered B cells present in A clusters: Proliferative (pink), Amoeboid (blue), Aggregated (green), and Phagocytic vs Ramified (yellow). **(C)** (C1) Average coupling distance between Proliferative (pink), Amoeboid (light blue), Aggregated (green), Phagocytic (yellow) vs Ramified (blue axis). (C2) Average coupling distance between the edge of the ganglionic eminence and each population. Frequencies of coupling are summarized in tables. **(D)** Individual versus population distances. We chose Ramified as the reference. (D1) Mean distance ($\mu m$) to the first neighbor that can be one of the 5 microglial types. (D2) distance ($\mu m$) to the shortest of the second population, averaged over all ramified cell position. (D3) Fraction of balls of radius $d^*$, centered around a microglia type containing at least one cell type inside. The distance $d^*$ is chosen at each time as the maximum among the mean coupling distances (d1, d2, d3, d4). (D4) Fraction of cells (Ramified cells) having another cell type as first neighbor, independently of any fixed ball.

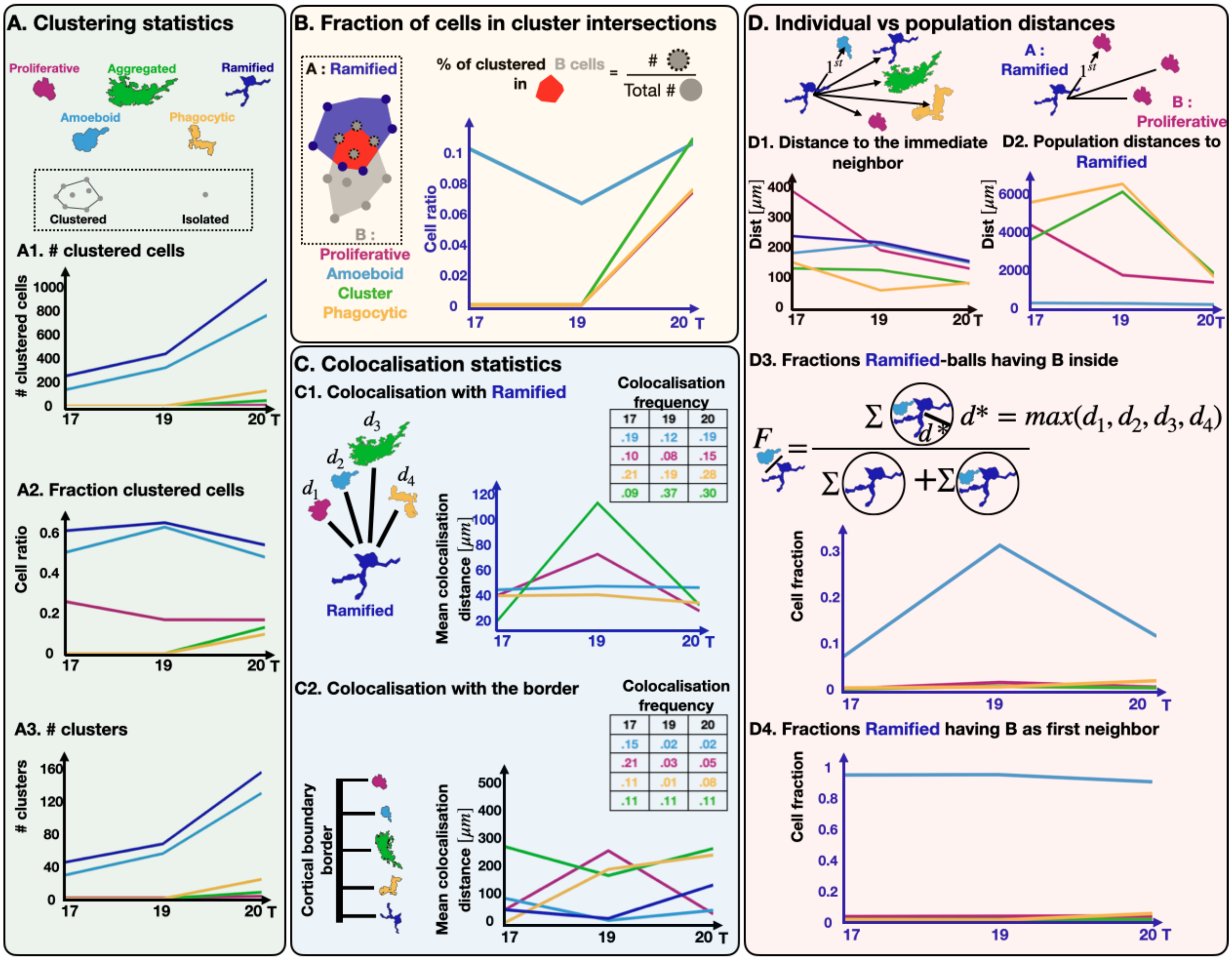

Figure S27: **Spatiotemporal statistics of microglia in the cortical boundary.** **(A)** Clustering of the different microglia types: Proliferative (pink), Amoeboid (light blue), Aggregated (green), Phagocytic (yellow), Ramified (dark blue). Clusters are computed using DBSCAN (MinSample = 4). (A1) Time evolution for the number of cluster cells, (A2) fraction of clustered cells, (A3) number of clusters. **(B)** Mixing quantification, computed as the fraction of clustered B cells present in A clusters: Proliferative (pink), Amoeboid (blue), Aggregated (green), and Phagocytic vs Ramified (yellow). **(C)** (C1) Average coupling distance between Proliferative (pink), Amoeboid (light blue), Aggregated (green), Phagocytic (yellow) vs Ramified (blue axis). (C2) Average coupling distance between the edge of the ganglionic eminence and each population. Frequencies of coupling are summarized in tables. **(D)** Individual versus population distances. We chose Ramified as the reference. (D1) Mean distance ($\mu m$) to the first neighbor that can be one of the 5 microglial types. (D2) distance ($\mu m$) to the shortest of the second population, averaged over all ramified cell position. (D3) Fraction of balls of radius $d^*$, centered around a microglia type containing at least one cell type inside. The distance $d^*$ is chosen at each time as the maximum among the mean coupling distances (d1, d2, d3, d4). (D4) Fraction of cells (Ramified cells) having another cell type as first neighbor, independently of any fixed ball.

| **Supplementary Table: Microglia and myeloid functional gene lists** | | |
|---|---|---|
| **Functional group** | **Genes** | **References** |
| Microglial molecular markers | tmem119, cx3cr1, csf1r, itgam, itgax, p2ry12, spi1, aif1 | Kracht et al, 2020, Bennett et al, 2016 |
| Homeostatic microglia | tmem119, p2ry12, p2ry13, adgrg1,tiam1,hist1h2bg,cx3cr1,vista | Kracht et al, 2020; Silva-Gomes et al, |
| Phagocytic/activated microglia | apoe, cd68, axl, hla-dr(s), siglec14, chp1,apoc1, asah1,clec7a | Kracht et al, 2020 |
| Immediate early genes | (c)jun,fos, ccl4,ccl4l2, ccl3l1,dusp1 | Kracht et al, 2020; Silva-Gomes et al, |
| Proliferative microglia | mki67, spc24, iqgap3, cenpf, trim59, birc5 | Kracht et al, 2020 |
| Temporally-restricted phagocytic genes but could contain neuronal transcripts | meg3, kif5a, map1b,sox11,cxadr,sox4,igf2bp1 | Kracht et al, 2020 |
| MRPL23 clusters | mrpL23, ac004556.1, polr2e, cnbp, rplp0 | Kracht et al, 2020 |
| PARP4 | parp4, parp4p2, al354798.1 | Kracht et al, 2020 |
| MTX1 | mtx1, mtx1p1, gpx1, hspa8, rpl35 | Kracht et al, 2020 |
| HBG2 | hbg2, hba2, hgb1, hba1, hbb | Kracht et al, 2020 |
| ZP3 | pompzp3, zp3 | Kracht et al, 2020 |
| NAMPT | NAMPT, NAMPTP1 | Kracht et al, 2020 |
| Myeloid cluster 1 (non-microglia) and border associated macrophages | lyve1, f13a1, rnase1, mrc1, dab2, ms4a family genes | Kracht et al, 2020; Silva-Gomes et al, |
| Myeloid cluster 2 (non-microglia) and border associated macrophages | S100 A9, lilra5, s100a8, lyz, dock5, ms4a family genes | Kracht et al, 2020, Silva-Gomes et al, |
| Immunocompetent microglia | itgax, clec7a, pkm, axl, tyrobp, p2ry12, cx3cr1, c3, cd68 | Kracht et al, 2020 |
| Migrating microglia | ifngr1 | Boghozian et al, 2023 |
| Apoptotic microglia | caspase3, bax, p53, fas, tnf | Sierra et al., 2013 |
| Myelin related microglia | itgax, clec7a, gpnmb, spp1 | Butovsky et al, 2018 |
| **Microglia markers** | **Genes** | **Morphological type** |
| Molecular markers, may not appear unless upregulated (common to all microglia) | tmem119, cx3cr1, csf1r, itgam, itgax, p2ry12, spi1, aif1 | Ramified |
| Homeostatic microglia markers | tmem119, p2ry12, p2ry13, adgrg1,tiam1,hist1h2bg,cx3cr1,vista,clec7a | Ramified |
| Microglia markes of phagocytosis/activation (could be common to all microglia clusters) | apoe, cd68, axl, hla-dr(s), siglec14, chp1,apoc1, asah1 | Phagocytic/clustered |
| Proliferative microglia but could be in all clusters, but one should stand out according to temporal window considered | mki67, spc24, iqgap3, cenpf, trim59, birc5 | Proliferative |
| Immunocompetent microglia | itgax, clec7a, pkm, axl, tyrobp, p2ry12, cx3cr1, c3,ifgnr1 | Amoeboid/phagocytic |

**Supplementary Table 1: Microglia and myeloid functional gene lists**

| Supplementary Table: Case demographics | | | | | | |
|---|---|---|---|---|---|---|
| **Case** | **GA** | **PCW** | **Sex** | **Source** | **Cause of Death** | **Histology** |
| Fetus 1 | 12 | 10 | F | HIIM | Medical abortion | Normal |
| Fetus 2 | 12 | 10 | F | HIIM | Medical abortion | Normal |
| Fetus 3 | 13 | 11 | M | OBB | Termination of pregnancy | Normal |
| Fetus 4 | 13 | 11 | F | HIIM | Termination of pregnancy | Normal |
| Fetus 5 | 13 | 11 | F | HIIM | Termination of pregnancy | Normal |
| Fetus 6 | 14 | 12 | F | OBB | Termination of pregnancy | Normal |
| Fetus 7 | 14 | 12 | F | HDBR | Termination of pregnancy | Cortical haemorrhages |
| Fetus 8 | 14 | 12 | n/k | HIIM | Medical abortion | Normal |
| Fetus 9 | 15 | 13 | M | HDBR | Termination of pregnancy | Normal |
| Fetus 10 | 15 | 13 | n/k | HIIM | Medical abortion | Normal |
| Fetus 11 | 16 | 14 | M | OBB | Termination of pregnancy | Normal |
| Fetus 12 | 16 | 14 | M | HDBR | Termination of pregnancy | Cortical haemorrhages |
| Fetus 13 | 16 | 14 | M | HDBR | Termination of pregnancy | Cortical haemorrhages |
| Fetus 14 | 17 | 15 | M | HDBR | Termination of pregnancy | Cortical haemorrhages |
| Fetus 15 | 17 | 15 | n/k | HIIM | Termination of pregnancy | Normal |
| Fetus 16 | 18 | 16 | M | OBB | Termination of pregnancy | Normal |
| Fetus 17 | 18 | 16 | n/k | HIIM | Termination of pregnancy | Normal |
| Fetus 18 | 19 | 17 | M | OBB | Termination of pregnancy | Normal |
| Fetus 19 | 19 | 17 | F | OBB | Termination of pregnancy | Normal |
| Fetus 20 | 20 | 18 | M | HDBR | Termination of pregnancy | Cortical haemorrhages |
| Fetus 21 | 21 | 19 | M | OBB | Termination of pregnancy | Normal |
| Fetus 22 | 21 | 19 | M | HDBR | Termination of pregnancy | Cortical haemorrhages |
| Fetus 23 | 21 | 19 | M | OBB | Termination of pregnancy | Normal |
| Fetus 24 | 22 | 20 | M | OBB | Miscarriage | Normal |
| Fetus 25 | 23 | 21 | F | HIIM | Termination of pregnancy | Normal |
| Fetus 26 | 27 | 25 | F | HIIM | Placental abruption | Normal |
| Fetus 27 | 27 | 25 | F | HIIM | Placental abruption | Normal |
| Fetus 28 | 33 | 31 | n/k | HIIM | Respiratory distress | Normal |
| Fetus 29 | 38 | 36 | F | HIIM | Cardiorespiratory arrest | Normal |
| Fetus 30 | 38 | 36 | F | HIIM | Pulmonary hypoplasia | Normal |
| Fetus 31 | 40 | 38 | M | OBB | Maternal hypertension | Hypoxic injury |
| Total number of cases shown is n = 31. *F:* female; *GA:* gestational age; *HDBR*: Human Developmental Biology Resource; *HIIM:* Croatian Institute for Brain Research; *M:* male; *n/k:* not known; *OBB*: Oxford Brain Bank; *PCW*: postconceptional week. | | | | | | |

**Supplementary Table 2: Demographics of cases**

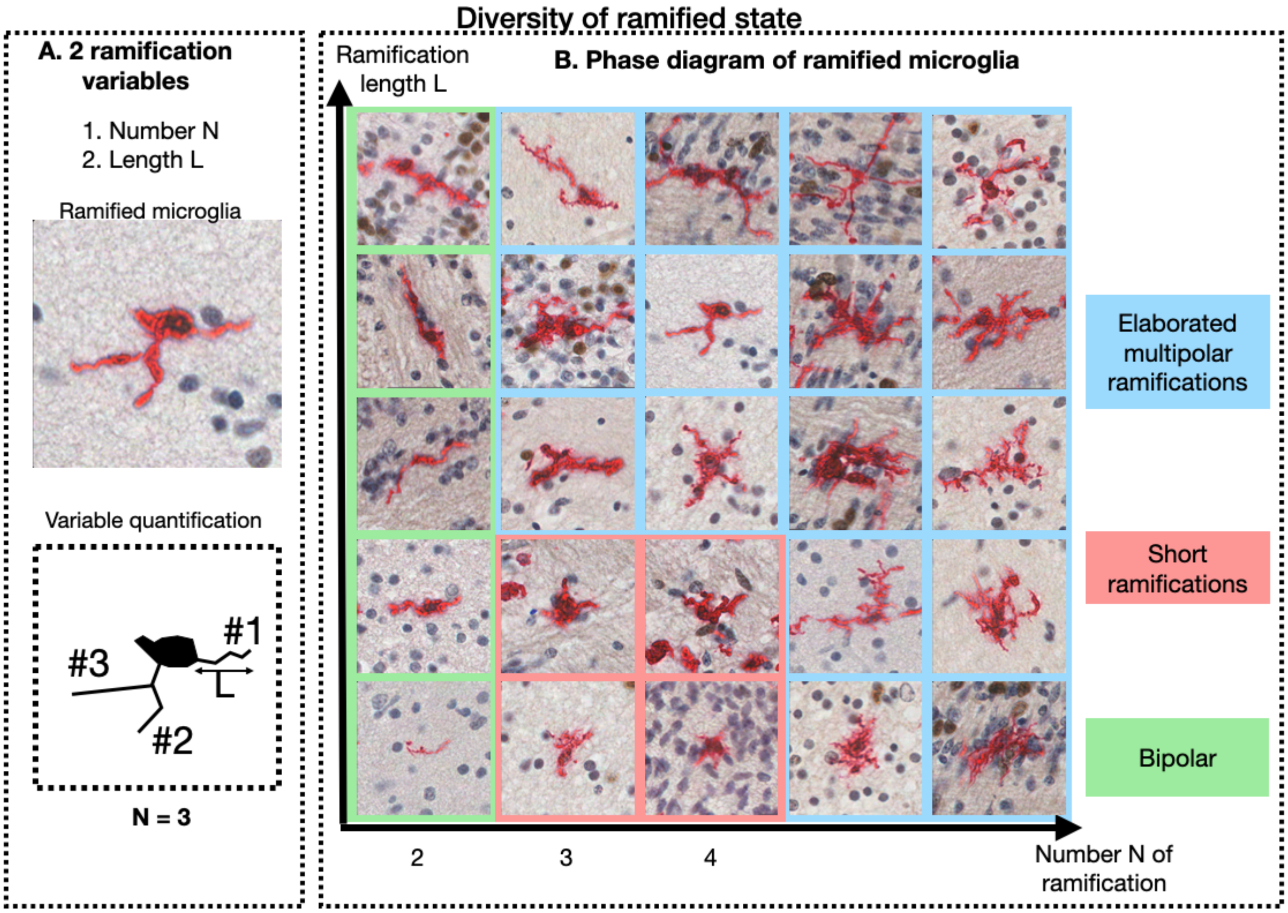

Figure S30: **Ramified cell heterogeneity** **(A)** Characterisation of ramified cells by their number $N$ and length $L$. **(B)** Matrix of Ramified cells distributed according to N and L. Cells can be clustered in 3 groups : bipolar, short ramification, and elaborated multipolar ramifications.

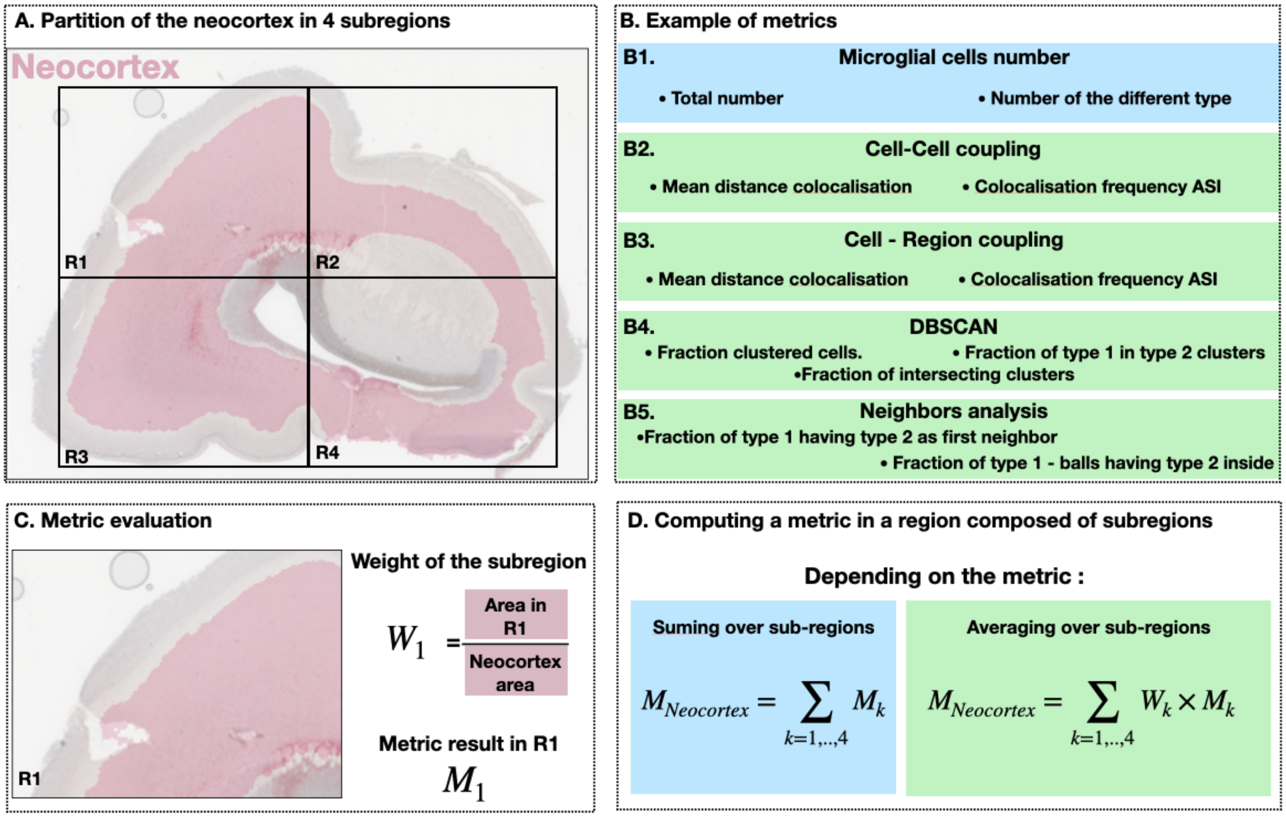

Figure S31: **Metric computation in large regions divided in sub-regions.** **(A)** Neocortex region decomposed into four sub-regions (R1,..,R4). **(B)** Various metrics are presented in categories. **(C)** The metric $M_k$ is calculated on each rectangle $R1, ..R4$, and the weight $W_k$ is the fraction of the region's area in the square. **(D)** The metric $M_k$ computed over the entire image is obtained by summing over the different regions $R_k$, either as a sum of all $M_k$ or weighted by $W_k$ to account for the difference in the areas.
